# Computational design of a versatile, zero-radius proximity labeling enzyme

**DOI:** 10.64898/2026.09.24.754223

**Authors:** Shizhong A. Dai, Peter E. Cavanagh, Albert Qiang, Zijuan Zhang, Qixiang Geng, Maribel Anguiano, Andrew G. Xue, Tsutomu Matsui, Khanh Nguyen, Namrata D. Udeshi, Sam Pellock, Brian Hie, Steven A. Carr, Christina Kim, Nathanael S. Gray, Wei Qin, Alice Y. Ting

**Affiliations:** Department of Genetics, Stanford University.; Department of Biochemistry, Stanford University.; School of Pharmaceutical Sciences, Tsinghua-Peking Center for Life Sciences, Tsinghua University.; Chemical and Systems Biology, Stanford University.; Princeton Neuroscience Institute; Omenn-Darling Bioengineering Institute; Biophysics program, Stanford University; Stanford Synchrotron Radiation Lightsource, SLAC National Accelerator Laboratory, Stanford University.; Broad Institute of MIT and Harvard; Institute for Protein Design; Department of Chemical Engineering and Stanford Data Science, Stanford University.; Arc Institute; Howard Hughes Medical Institute; Departments of Biology, and by courtesy, Chemistry; Chan Zuckerberg Biohub – San Francisco; The Phil & Penny Knight Initiative for Brain Resilience at the Wu Tsai Neurosciences Institute, Stanford University.

## Abstract

The ability to map protein interactomes and organelle proteomes is foundational for achieving a molecular understanding of living cells. Proximity labeling (PL) provides a powerful strategy for this, but existing enzymes and photocatalysts are limited by their spatial resolution, reliance on biotin, and/or *in vivo* compatibility. Here we report FlexID, an engineered promiscuous ligase that catalyzes the rapid attachment of diverse small-molecule probes to proximal endogenous proteins. Critically, FlexID operates through a zero-radius, direct-contact mechanism, offering superior spatial precision compared to existing PL tools. We engineered FlexID by combining the strengths of sequence-and structure-trained computational models to enhance its catalytic activity and structural stability. Biophysical analysis revealed that specific conformational changes in FlexID improve its ability to recognize diverse target proteins while simultaneously preventing the premature release of the reactive intermediate. We demonstrate FlexID’s versatility through *in vivo* proximity labeling, comprehensive organelle proteome mapping, and a high-throughput, fluorescence-based screen for molecular glues. Our work shows that computational methods can be harnessed to create mechanistically distinct PL enzymes and establishes FlexID as a flexible, high-resolution tool for mapping protein interactions and proteomes in living cells.

## Main

PL has emerged as a cornerstone technique for mapping interactomes, organelle proteomes, and subcellular transcriptomes in living cells^1^. The prevailing approach is to use either genetically-encoded enzymes (e.g., TurboID^2^ and APEX^3^) or chemical catalysts^4,5^ to produce highly-reactive, short-lived species that diffuse from the site of generation and covalently tag endogenous proteins and RNA within 1-20 nm (**Figure 1B**). Labeled proteins or RNA are subsequently enriched from cell lysates and identified by mass spectrometry or sequencing. While existing PL methods have produced many valuable interactome and organelle maps^1^, the fact that labeling radius varies across environments (for example, the cell interior has millimolar concentrations of glutathione, a quencher of APEX-and HRP-generated phenoxyl radicals, while the extracellular environment does not), and the difficulty of distinguishing between direct interaction partners and proximal neighbors, fundamentally limits the utility of these methods.

**Figure 1.**
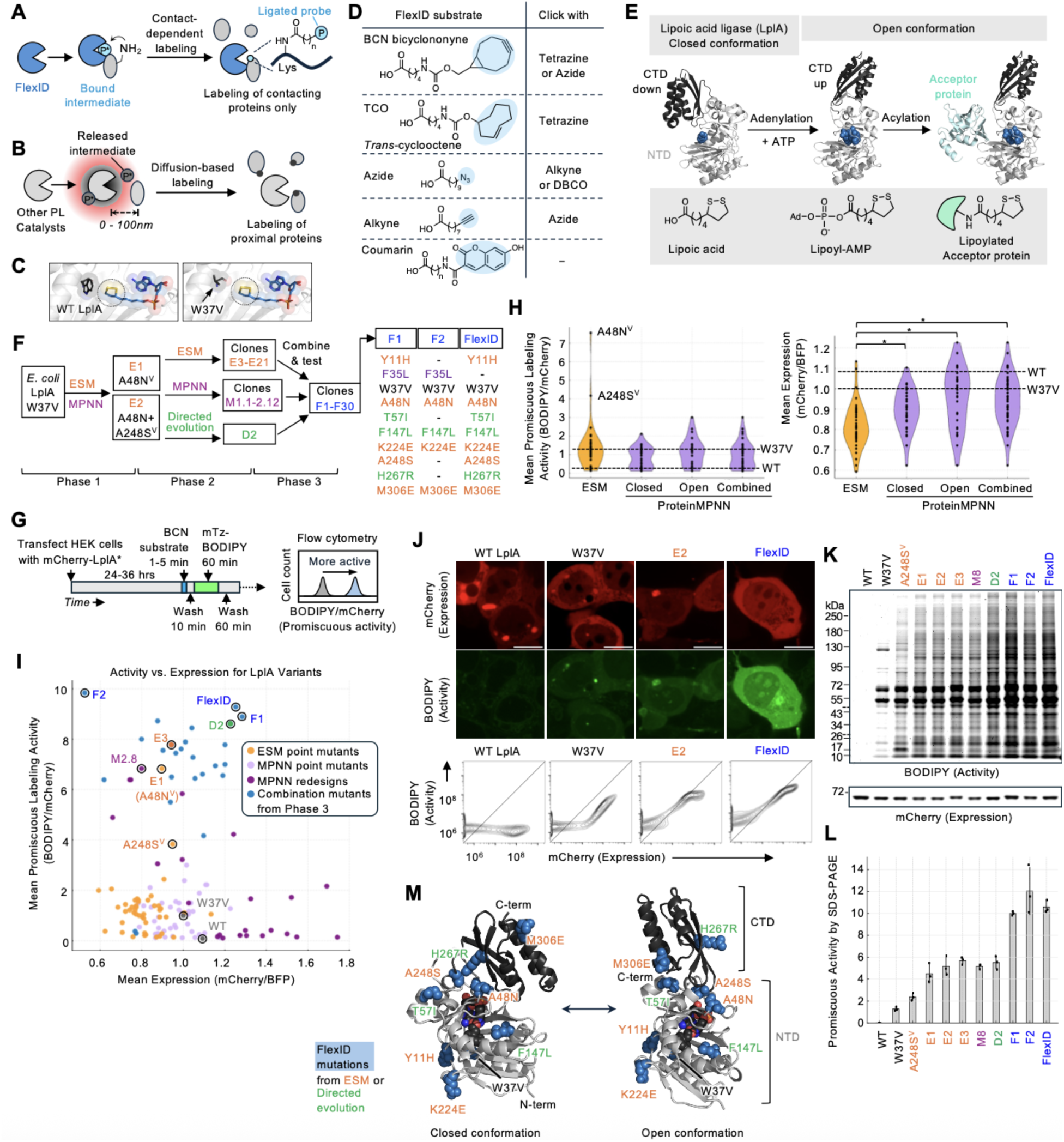
Computational design of FlexID. (**A**) FlexID catalyzes contact-dependent, promiscuous labeling of endogenous proteins with diverse small-molecule probes (P). (**B**) Other proximity labeling (PL) catalysts tag neighboring proteins via release of a diffusible reactive intermediate. P*, reactive intermediate. (**C**) W37V mutation in LplA allows it to accept unnatural probes. Dithiolane ring of bound lipoyl-AMP is circled. (**D**) Substrates of FlexID, including Click-chemistry handles that can be bio-orthogonally derivatized. (**E**) Lipoic acid ligation catalyzed by wild-type *E. coli* LplA, the template for FlexID. Adenylation is catalyzed by the closed conformation, while lipoyl transfer onto acceptor proteins (light blue) is catalyzed by the open conformation. CTD, C-terminal domain. NTD, N-terminal domain. Ad, adenosine. Structures are from PBD IDs 1X2G^25^ (left), 3A7R^26^ (middle), and 3A7A^26^ (right). (**F**) Flow chart showing the engineering of FlexID from W37V LplA in three phases. The V superscript indicates that the mutation is on the W37V template. Mutations in F1, F2, and FlexID are colored by whether they originated from ESM (orange), ProteinMPNN re-design (purple), or directed evolution (green). Detailed characterization of all variants shown in **Extended Data Figures 1-2** and **Supplementary Data 1**, and all sequences provided in **Table 1**. (**G**) Flow cytometry assay used to measure promiscuous labeling activities of mCherry-fused LplA variants in live HEK 293T cells. mTz, methyltetrazine. (**H**) Violin plots of measured activity and expression for ESM and ProteinMPNN-designed LplA point mutants. MPNN mutants are further separated into those designed on the closed (1X2G^25^) vs. open (3A7R^26^) backbone structures of LplA. ESM *N*=48, Open *N*=30, Closed *N*=32, MPNN Combined *N*=51 *p<0.05 (Student’s T-test with Bonferroni correction). (**I**) Activity vs. expression plot (both normalized to template W37V) for all LplA variants. BFP was used as a transfection marker. (**J**) Confocal microscopy of HEK 293T cells labeled by LplA variants with BCN substrate for 5 minutes, followed by live-cell Click with mTz-BODIPY as in (G). Scale bars, 20 µm. Bottom: corresponding flow cytometry plots obtained under the same conditions. (**K**) Promiscuous activities of FlexID and LplA variants in HEK 293T cells with BCN labeling for 5 minutes. After Click with mTz-BODIPY as in (G), lysates were analyzed by SDS-PAGE and in-gel fluorescence. (**L**) Quantification of data in (K). *N*=3, Mean ± SD. (**M**) Open and closed structures of FlexID in complex with BCN-AMP, predicted by Chai-1.

We envisioned that a promiscuous enzyme that labels protein targets by a contact-dependent mechanism, without release of reactive species into solution (**Figure 1A**), might offer a superior, higher-resolution method for dissecting interactomes and spatial proteomes. While the concept of contact-dependent labeling and high sequence promiscuity may seem incompatible, several natural enzymes operate by such a mechanism, including ubiquitin ligases^6^, some kinases^7^, and some histone acetyltransferases^8^. We wondered whether it would be possible to engineer a new PL enzyme that labels its targets by a contact-dependent, zero radius mechanism, yet remains agnostic as to target sequence.

Here we use a combination of engineering approaches – including mutation prediction by a protein language model (ESM^9,19,20^) and the inverse folding model ProteinMPNN^10^ – to create such an enzyme. Our engineered “FlexID” enzyme labels proximal endogenous proteins by a contact-dependent, zero-radius mechanism, and does so with a variety of non-biotin probes, enabling *in vivo* proteome labeling without interference from endogenous biotin or biotinylated proteins. Characterization shows that FlexID is fast (5 minutes), non-toxic, active in multiple mammalian organelles, and produces more spatially specific proteomes than both APEX and TurboID, without a reduction in sensitivity. In addition, we leverage the high spatial specificity and flexible substrate utilization of FlexID to develop a high throughput protein-protein interaction and protein localization screening platform called “PLF” (proximity labeling with fluorescence readout). We use PLF to perform a live-cell screen of molecular glues that recruit endogenous neo-substrates to the E3 ubiquitin ligase cereblon. FlexID provides a powerful and versatile tool for high-resolution mapping of cellular interactomes and organelle proteomes.

### Engineering FlexID for fast, bio-orthogonal proximity labeling in cells

To create FlexID, we selected the enzyme lipoic acid ligase as our template. These enzymes catalyze site-specific, ATP-dependent conjugation of the cofactor lipoic acid onto acceptor proteins in a process essential for energy metabolism, cell growth, and biosynthesis^11^. Notably, the *E. coli* ortholog, LplA, is structurally homologous to *E. coli* BirA (biotin ligase), the template used to engineer TurboID^2^. Furthermore, previous engineering identified an active site residue in LplA (W37, **Figure 1C**) that when mutated to valine enlarges the substrate binding pocket to allow conjugation of diverse unnatural probes in place of lipoic acid, including azides^12,13^, alkynes^12^, bicyclononyne (BCN)^14^, trans-cyclooctene (TCO)^15^, coumarin^16^, and resorufin^17^ (**Figure 1D**). The ability to perform PL with diverse synthetic probes would expand the scope of possible applications, circumvent background from endogenous biotin and endogenous biotinylated proteins (which are especially problematic *in vivo*), and improve the temporal precision of PL.

Lipoic acid ligases are thought to be highly sequence-specific. For example, *E. coli* LplA labels only three proteins in the entire *E. coli* proteome^11^. To test LplA in the mammalian cell setting, we expressed the W37V mutant in the cytosol as a fusion to mCherry and supplied the Click substrate BCN (**Figure 1D**). After in-cell Click derivatization with methyltetrazine-BODIPY, cells were lysed and analyzed by SDS-PAGE and flow cytometry^18^ (**Figure 1G**). **Figures 1J-L** show that some promiscuous tagging of endogenous mammalian proteins was observed, but both the extent and breadth were too low to be useful. In case orthologs from other species may have higher starting promiscuous activities, we cloned 11 diverse lipoic acid ligases as X37V mutants and expressed them in HEK 293T cells to perform the same flow cytometry-based assay (**Extended Data Figures 1A-C**). None of these showed higher promiscuous activity than *E. coli* W37V LplA, though some mesophiles and thermophiles (but not cryophiles) exhibited detectable labeling.

Using *E. coli* W37V LplA as our starting template, we applied two computational strategies to increase promiscuous activity (**Figure 1F**). First, we used the protein language models ESM-1b^19^ and ESM-1v^20^ which are trained on 250 million and 98 million natural protein sequences, respectively. ESM was previously used to identify affinity-enhancing mutations in antibodies^21^ and classify missense variants as pathogenic or benign^22^. We used ESM to predict high-scoring or “evolutionarily plausible” mutations in LplA, reasoning that some of these may enhance promiscuous activity by allowing LplA to sample alternative conformations or even mechanisms used by proteins of related sequence. Second, we applied the inverse folding model (IFM) ProteinMPNN^10^ that is trained on 19,700 high-resolution single-chain structures from the Protein Data Bank (PDB) to recover their wild-type sequences. Mutations predicted by IFMs have been shown to enhance protein stability^23,24^. Solved structures of LplA show the protein in two strikingly different conformations (**Figure 1E**) – a “closed” conformation in which the C-terminal domain (CTD) is oriented downward over the active site^25^, and an “open” conformation in which the CTD is extended and away from the active site^26^. We used both structures as inputs for ProteinMPNN-based mutation prediction, unsure what effect stabilizing mutations on either structure would have on the overall promiscuous activity of the enzyme.

48 ESM-designed mutations and 51 ProteinMPNN-designed mutations (30 on the open structure and 32 on the closed structure, with 11 overlapping) were introduced onto the W37V LplA template (indicated by the V superscript) and tested in HEK 293T cells using the flow cytometry assay in **Figure 1G**. We observed a wide range of promiscuous activities and expression levels (**Figure 1H-I**). In general, ProteinMPNN-predicted mutants showed significantly better expression than mutants predicted by ESM (**Figure 1H** right). However, ESM mutants were on the whole more promiscuous, skewed especially by two variants with 7.6-and 4.2-fold higher promiscuous activities than W37V template (**Figure 1H** left). Though 200 amino acids apart in the linear protein sequence, the A48N and A248S mutations are adjacent in the protein structure, and both map the hinge region between the N-terminal domain (NTD) and CTD (**Figure 1M**). This observation led us to hypothesize that conformational dynamics may play a role in determining LplA’s sequence-specificity, which is explored below.

We moved forward with these two ESM-identified mutations, alone and in combination (clones E1 and E2 in **Figure 1F**), but noticed that they reduced the stability of LplA, giving rise to mCherry-LplA aggregates visible by microscopy (**Figure 1J**), and a decrease in apparent melting temperature as measured by differential scanning fluorimetry (DSF, **Extended Data Figure 1I**). To recover LplA stability while further enhancing its promiscuous activity, in our second phase of engineering (**Figure 1F**), we combined E1 and E2 with: (1) three expression-and stability-enhancing mutations previously identified by yeast display evolution of LplA^27^ (2) other higher-activity and higher stability ESM-identified point mutations from **Figure 1H**, and (3) mutations identified by ProteinMPNN-based redesign of residues surrounding the W37V (**Extended Data Figure 2C**). Of these, we found that the directed evolution mutations were beneficial in eliminating mCherry-LplA puncta in microscopy and BODIPY signal saturation in 2D flow plots (**Extended Data Figure 1F-H**). Mixing and matching mutations (Phase 3 of engineering) produced our best penultimate clones (F1 F2), with mutations contributed from all three strategies (**Figure 1F**).

For our final version of FlexID, we selected F1 for its superior expression but removed ProteinMPNN-designed F35L to slightly enhance its activity (**Figures 1F, I, K-L**). FlexID has 8 mutations relative to W37V LplA template: 5 from ESM and 3 from directed evolution. These mutations are localized throughout the structure (**Figure 1M**). In addition to its high promiscuity, FlexID has other favorable characteristics, including smooth non-punctate expression in HEK293T cells (**Figure 1J**), and higher T_m_ compared to E1 and E2 by DSF (**Extended Data Figure 1I**). In contrast to E1 which exhibits a significant dimeric population, FlexID appears mostly monomeric by size exclusion chromatography-small angle X-ray scattering (SEC-SAXS, **Extended Data Figure 1J**).

### FlexID catalyzes promiscuous labeling by a contact-dependent mechanism

We wished to understand how the mutations in FlexID change the enzyme from sequence-specific to highly promiscuous. The strongest promiscuity-enhancing mutations in FlexID, A48N and A248S, map to the hinge region between LplA’s N-terminal domain (NTD) and CTD (**Figure 1M**), raising the possibility that conformational dynamics play a role in determining LplA’s sequence specificity. Crystal structures show a major rotation of the CTD with respect to the NTD, from the closed conformation when LplA is bound to lipoic acid^26^ to the open conformation when LplA is bound to lipoyl-AMP^26^ or acceptor protein^26^ (**Figure 1E**). The purpose of this conformational rearrangement is to reconfigure the active site from one that catalyzes lipoic acid adenylation to one that catalyzes the second half-reaction, lipoyl transfer onto acceptor proteins; in the open but not closed conformation, the active site is accessible to docked acceptor proteins.

In recent work, we showed that point mutants of LplA computationally predicted to favor the open conformation exhibit higher promiscuous activity, while point mutants biased toward the closed conformation have lower promiscuous activity^18^. This effect could arise if the CTD of LplA acts as a tethered competitive inhibitor that blocks access of acceptor proteins to the active site in the closed conformation (**Figure 2A**). Mutations that favor the open conformation may alleviate this inhibition, increasing affinity and decreasing K_m_ for non-cognate protein substrates, and producing greater promiscuous labeling. On the other hand, mutations that favor the closed conformation would increase CTD competition and produce even more specific labeling than template.

**Figure 2.**
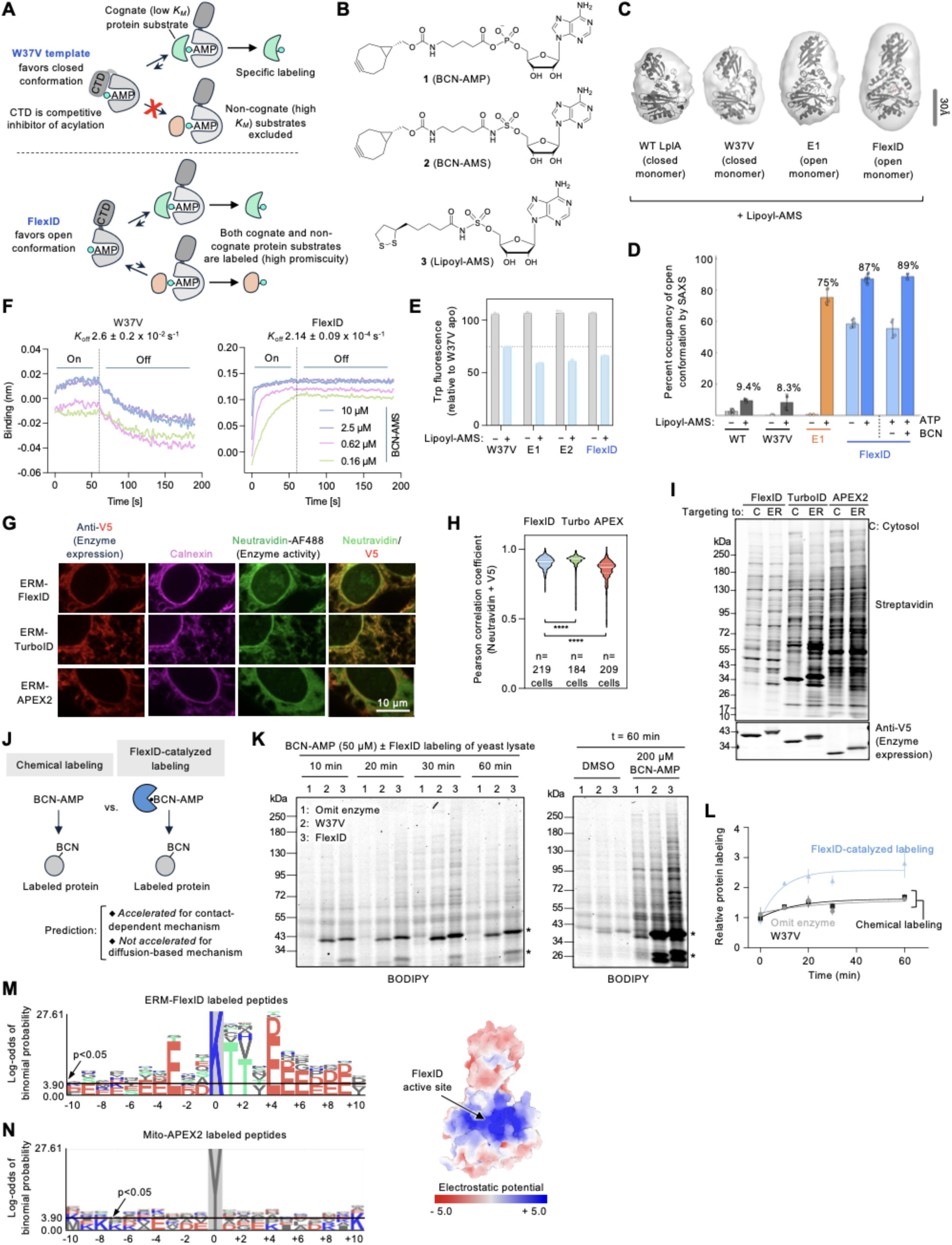
FlexID catalyzes promiscuous labeling via a contact-dependent mechanism. (**A**) Schematic showing how changes in conformational equilibrium between open and closed states could alter LplA promiscuity. Top, W37V prefers the closed adenylate-bound conformation, whose downward CTD position reduces off-target labeling. Bottom, FlexID prefers the open conformation which is more promiscuous. (**B**) Structures of LplA ligands used for biophysical characterization. **1** (BCN-AMS) is a non-hydrolyzable analog of **2** (BCN-AMP). Syntheses in **Extended Data** Fig. 4A**-D**. (**C**) SEC-SAXS analysis of LplA variants. DENSS ab initio modeling of protein envelopes, fit with major LplA conformer detected under +lipoyl-AMS condition. Envelopes are representative of 2-5 independent experiments per variant. (**D**) Fraction of each protein in the open conformation, predicted from SEC-SAXS in the apo state and after lipoyl-AMS binding. FlexID was also tested under +ATP and +ATP+BCN conditions. Conformational occupancy is based on Oligomer modeling from 2-6 independent measurements per variant. Mean ± SD. (**E**) Intrinsic tryptophan fluorescence of purified LplA variants before and after complexation to lipoyl-AMS. This experiment was performed three times. Mean ± SD. (**F**) Kinetics of BCN-AMS dissociation from biotin-immobilized W37V (left) and FlexID (right) by Bio-Layer Interferometry (BLI). This experiment was performed two times with similar results. (**G**) Confocal imaging of FlexID, APEX2, and TurboID targeted to the ER membrane (ERM) facing cytosol in HEK 293T cells. Labeling times were 1 minute for APEX and 10 minutes for TurboID and FlexID. FlexID-catalyzed BCN ligation was detected after fixation by Click with biotin-methyltetrazine. All samples were stained with neutravidin-AF488 and antibodies against V5 (LplA expression tag) and calnexin (endogenous ER marker). Scale bar, 10 μm. This experiment was performed three times with similar results. Additional fields of view in **Extended Data** Fig. 4H (**H**) Pearson correlation coefficients for anti-V5/neutravidin channels. ^∗∗∗∗^p < 0.0001 (Kruskal-Wallis test with Dunn’s nonparametric multiple comparison test). (**I**) Comparison of labeling “fingerprints” for FlexID, TurboID, and APEX2 targeted to the ER membrane or cytosol (“C”) of HEK293T cells. Labeling was performed as in (G), and cell lysates were Clicked with biotin-methyltetrazine for SDS-PAGE analysis. (**J**) Testing if BCN-AMP acylation of endogenous proteins is accelerated by FlexID, as would be expected for a contact-dependent mechanism but not a diffusion-based mechanism. (**K**) Results from experiment in (J). Chemically synthesized BCN-AMP was added to concentrated yeast lysate, either alone or pre-complexed with 25 uM purified W37V LplA or FlexID. After incubation for the indicated times, labeled samples were quenched and Clicked with mTz-BODIPY and analyzed by SDS-PAGE. *Self-labeling of FlexID/W37V. This experiment was performed twice with similar results. Additional analyses and controls in **Extended Data** Fig. 4I**-L**. (**L**) Quantification of lane intensities from (K). (**M**) FlexID substrate sequence preference. Peptides labeled by FlexID-ERM in HEK 293T cells (5 minutes BCN) were enriched and sequenced by mass spectrometry. Amino acid enrichment quantified by pLogo^44^ from 3 biological replicates (*N*=175 peptides). Data in **Extended Data Table 2**. Right: Surface electrostatic potential of FlexID, generated by AlphaFold3 and APBS Electrostatics plugin to PyMOL. (**N)** pLogo^44^ plot of previous data^31^ from 1-minute mito-APEX2 labeling in HEK 293T cells (*N*=273 peptides).

We hypothesized that FlexID may be biased towards the open conformation and exhibit increased promiscuity through this mechanism. Two mutations in FlexID are predicted by our “Conformational Biasing” computational method^18^ to be open-biasing (A48N from ESM and H267R from directed evolution; **Extended Data Figure 3**). In addition, Boltz-2^28^ predicts FlexID’s structure to be in the open conformation when bound to adenylate, whereas W37V LplA is predicted to be closed (**Extended Data Figure 4E**). To test this experimentally, we first used SEC-SAXS to measure the conformational distribution of different LplA variants (**Figures 2C-D** and **Extended Data Figure 5**). In the apo state, the enzyme was almost entirely in the closed conformation. After complexation with lipoyl-AMS, a non-hydrolyzable analog of lipoyl-AMP (**Figure 2B**), W37V LplA opened slightly, to 8.3%. FlexID, on the other hand, showed significantly greater occupancy of the open conformation in both the apo form (58%) and after binding to lipoyl-AMS (87%). We also used tryptophan fluorescence^18^ to probe the conformational transition of both enzymes upon lipoyl-AMS binding. FlexID showed a greater change in tryptophan fluorescence than W37V, consistent with a greater change in open state occupancy (**Figure 2E**).

These data support a model in which FlexID’s mutations shift its conformational preference to the open state, enhancing promiscuous labeling of non-cognate substrates through increasing access to bound AMP intermediate as shown in **Figure 2A**. Importantly, this mechanism for promiscuity involves direct transfer of bound lipoyl-AMP to protein substrate rather than its release into solution – a contrast to the mechanism believed to be used by other PL enzymes such as TurboID, APEX, and BioID. For example, the R118G mutation in BioID reduces the affinity of the enzyme for biotin-AMP by 400-fold compared to wild-type BirA^29,30^. We used bio-layer interferometry (BLI) to measure the off-rate of a non-hydrolyzable analog of BCN-AMP (BCN-AMS, structure in **Figure 2B**) from FlexID. **Figure 2F** shows that the off-rate was slow, at 2.14 ± 0.09 x 10^-4^ s^-1^. Interestingly, this measured dissociation is even slower (by ∼100-fold) than that of W37V LplA. An explanation is suggested by the Boltz-2-predicted structures of these enzymes in complex with BCN-AMS (**Extended Data Figures 4E-F**). In the open FlexID structure, the adenylate-binding loop is stabilized through formation of a β-sheet between β11 and β13 (pLDDT > 0.9). By contrast, β11 is partially disordered in the closed structure of W37V LplA bound to BCN-AMS (0.7 < pLDDT < 0.9). These differences suggest that FlexID, in its preferred open conformation, may engage the adenylate intermediate more tightly than W37V LplA does in its closed conformation, resulting in a major decrease in off-rate.

We performed several additional experiments to probe the model that FlexID’s promiscuous labeling occurs via a contact-dependent mechanism. First, we used cell imaging to compare the spread of FlexID-labeled proteins to those labeled by APEX and TurboID. We found that FlexID-tagged proteins were tightly colocalized with the enzyme, consistent with contact-dependent labeling, whereas APEX labeled proteins were the most diffuse (**Figures 2G-H and Extended Data Figure 4H)**. Second, we used streptavidin blotting to compare the banding pattern (“fingerprints”) of proteins labeled by FlexID, TurboID, or APEX targeted to either the cytosol or the ER membrane facing cytosol. Because these compartments are overlapping, APEX - which labels via a diffusible biotin-phenoxyl radical - produced highly similar fingerprints, consistent with larger labeling radius and consequent tagging of overlapping proteomes (**Figure 2I**). By contrast, FlexID and TurboID fingerprints were more distinct between the two compartments, implying shorter labeling radii.

Third, we assessed whether FlexID could accelerate labeling of highly-concentrated yeast cell lysates – mimicking cytosolic protein concentrations in live HEK cells – when supplied with chemically-synthesized intermediate BCN-AMP. In a diffusion-based labeling mechanism, the enzyme is not needed after it generates acyl-adenylate. However, we observed that addition of FlexID dramatically enhanced labeling of lysate proteins by BCN-AMP, and also produced a banding pattern distinct from that of chemical labeling by BCN-AMP - supporting the model that FlexID participates in the acylation step of catalysis (**Figures 2J-L, Extended Data Figure 4I-L**). Finally, we sequenced FlexID-labeled peptides after BCN labeling at the ER membrane of HEK 293T cells (**Figure 2M**). In contrast to peptides labeled by APEX^31^, which show no amino acid preferences for residues flanking the labeled tyrosine (**Figure 2N**), FlexID peptides are enriched in negatively-charged amino acids surrounding the labeled lysine, complementing the positively-charged surface of LplA’s active site vestibule. The significant overrepresentation of glutamate at the –3 position also mirrors the natural substrate preference of wild-type LplA^26^. Overall, our data are consistent with a model in which FlexID labels endogenous proteomes through direct contact with active site-bound AMP intermediate, rather than its release into solution.

### Characterization of FlexID for proximity labeling applications

We first performed a timecourse to compare the speed of FlexID-catalyzed labeling to TurboID and APEX2 (**Figures 3A** and **Extended Data Figure 6A**). APEX2 was the fastest while TurboID and FlexID were comparable. Next, we assessed the activity of FlexID in multiple cellular compartments, using both SDS-PAGE and microscopy. **Figures 3B** shows strong labeling with BCN in 5 minutes in the mitochondrial matrix, outer mitochondrial membrane (OMM), ER membrane (ERM) facing cytosol, nucleus, and cytosol. Labeling in the ER lumen was weaker and required 30 min of BCN incubation. SDS-PAGE of labeled lysates showed distinct banding patterns across compartments, suggesting that unique proteomes are tagged in each locale (**Figure 3C**).

**Figure 3.**
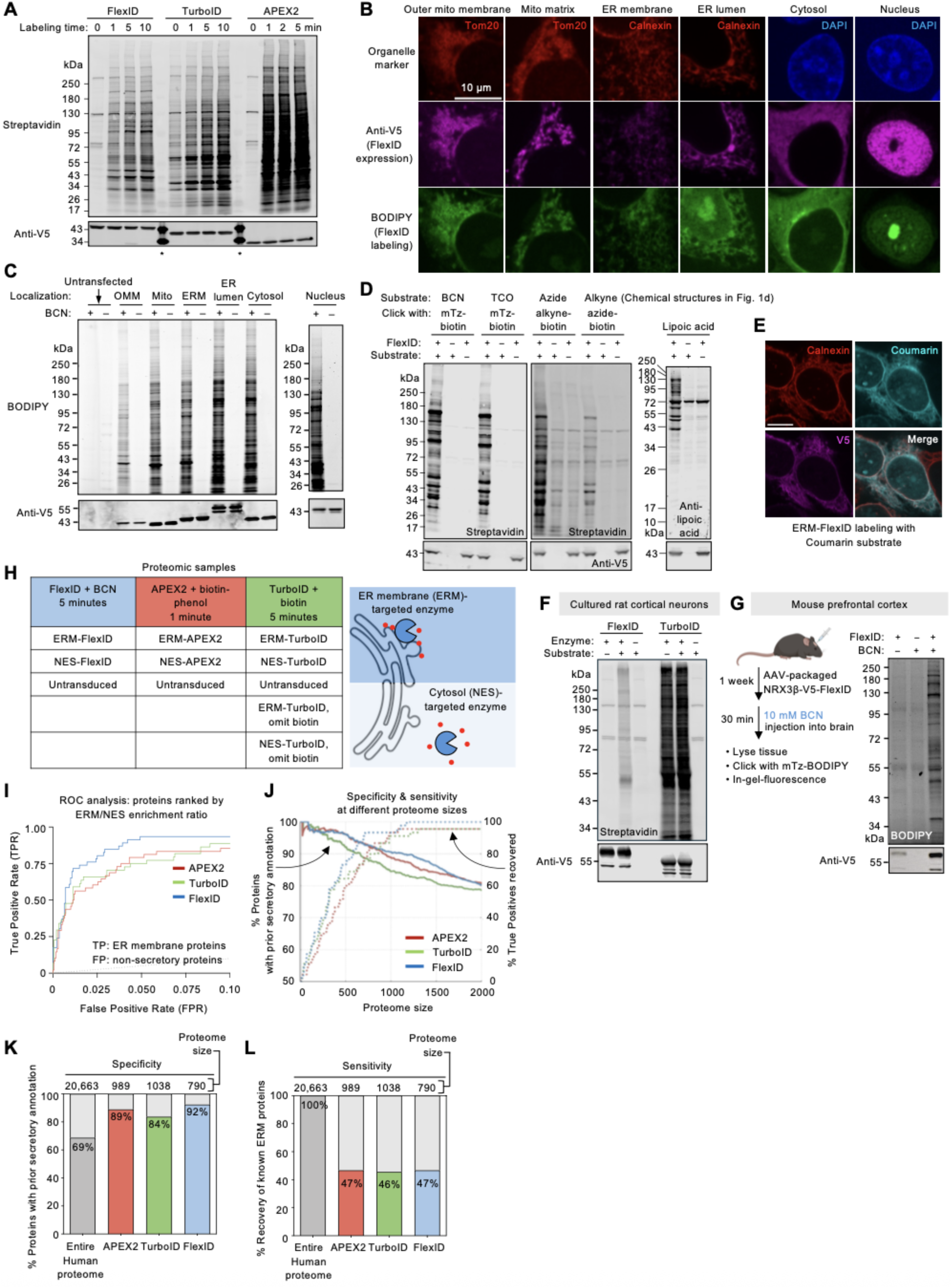
FlexID labeling of spatial proteomes. (**A**) Time course of proximity labeling (PL) catalyzed by FlexID with BCN, TurboID with biotin, and APEX2 with biotin-phenol. Enzymes targeted to the ER membrane facing cytosol of HEK 293T cells. BCN detected by Click with biotin-methyltetrazine after cell lysis. TurboID shows labeling even without exogenous biotin addition. Anti-V5 detects PL enzyme expression. *Molecular weight ladder. This experiment was performed twice with similar results. Same experiment with cytosol-targeted enzymes in **Extended Data** Figure 6A. (**B**) Confocal imaging of FlexID-catalyzed PL in various HEK 293T organelles. All samples labeled live with BCN for 5 minutes, except ER lumen FlexID which used 30 minutes. Click with mTz-BODIPY in live cells followed by washing, fixation, and imaging. Tom20 and calnexin are endogenous mitochondrial and ER markers, respectively. This experiment was performed >3 times with similar results. (**C**) SDS-PAGE analysis of samples labeled as in (B). BCN detected by Click with mTz-BODIPY after cell lysis. OMM, outer mitochondrial membrane facing cytosol. ERM, ER membrane facing cytosol. Anti-V5 blot detects FlexID expression. This experiment was performed >3 times with similar results. (**D**) FlexID uses diverse substrates. HEK 293T cells expressing ERM-FlexID were labeled live for 5 minutes with the indicated substrates (chemical structures in **Figure 1D**). Labeled proteins detected via Click with the indicated reagents after cell lysis. This experiment was performed >3 times with similar results. (**E**) 10-minute FlexID labeling with coumarin substrate (structure in **Figure 1D**). This experiment was performed twice with similar results. (**F**) FlexID labeling in primary rat cortical neurons. FlexID and TurboID were targeted to the neuronal membrane facing cytosol via fusion to Neurexin-3β, and labeled for 5 min with BCN (FlexID) or biotin (TurboID). BCN detected by Click with mTz-biotin after cell lysis. Anti-V5 detects PL enzyme expression. This experiment was performed twice with similar results. (**G**) *In vivo* labeling with FlexID. Neurexin3β-V5-FlexID was expressed in the prefrontal cortex of adult mice. This experiment was performed twice with similar results. (**H**) Design of proteomic experiment comparing FlexID, APEX, and TurboID-catalyzed PL at the ER membrane facing cytosol (ERM) of HEK 293T cells. Cytosolic constructs, fused to NES (nuclear export sequence) were used as spatial references. All samples were prepared in triplicate and analyzed by data-independent acquisition (DIA). (**I**) ROC (receiver operating characteristic) analysis comparing FlexID, TurboID and APEX proteomic data from (H). For every possible ERM/NES enrichment ratio cut-off, the true positive rate (TPR) was plotted against the false positive rate (FPR). True positives are 90 well-established ER membrane proteins; false positives are proteins lacking secretory annotation (**Table 3**). (**J**) Specificity (solid lines) and sensitivity (dashed lines) as a function of proteome size, for each PL enzyme. (**K-L**) Specificity (K) and sensitivity (L) of FlexID, TurboID, and APEX-mapped ERM proteomes, based on ROC-determined cut-offs (**Extended Data** Figure 7E and **Table 4**).

FlexID can be targeted to various baits and cell compartments through genetic fusion. However, we noticed that FlexID was sensitive to tagging at its C-terminal end, probably because such fusion can impede the conformational transition of its C-terminal domain (**Figures 1E, M**). We tested a series of flexible linkers, and identified a 12 amino acid Gly-Ser linker that preserves FlexID activity when mCherry is fused to its C-terminal end (**Extended Data Figures 6B-D**).

The ability to use non-biotin probes is especially useful in biological settings where endogenous biotin is abundant, such as neuron cultures. TurboID-catalyzed labeling occurs in neurons whether or not exogenous biotin is supplied (**Figure 3F**), preventing temporal control over the labeling time window. We tested FlexID’s ability to use diverse non-biotin substrates, as its template, W37V LplA, accepts many alternative substrates in place of lipoic acid^12–17^. Similarly, we found that FlexID can catalyze promiscuous labeling with lipoic acid, linear alkynes and azides, BCN, trans-cyclooctene (TCO), and coumarin (**Figure 3D-E**). Each Click-based probe is detected by its corresponding Click reaction partner in lysate (**Figure 1D**). When we expressed FlexID in neuron cultures, we observed that labeling depends strictly on addition of exogenous probe (BCN), in contrast to TurboID, and strong signal is detected after just 5 minutes of probe incubation (**Figure 3F**). The non-toxic conditions of FlexID labeling also permitted us to perform PL in the living mouse brain, with FlexID targeted to the neuronal membrane in the prefrontal cortex by AAV transduction, and BCN supplied via direct intracranial injection (**Figure 3G**). Promiscuously tagged endogenous proteins were detected only in the presence of both enzyme and probe, 30 min after BCN administration.

We then performed a proteomic experiment using FlexID to evaluate its specificity and depth of coverage (sensitivity) when mapping an “open compartment” - the ER membrane facing the cytosol (**Figure 3H**). HEK 293T cells were incubated with BCN for 5 min, then lysed and Clicked with mTz-biotin for streptavidin-based capture. We also prepared a sample targeting FlexID to the cytosol to serve as a spatial reference. For direct comparison to TurboID and APEX2, we prepared equivalent samples, treated with biotin for 5 minutes, or biotin-phenol and H_2_O_2_ for 1 minute, respectively (**Figures 3H, Extended Data Figure 7**). After streptavidin enrichment of all samples, on-bead trypsin digestion, and data-independent acquisition of mass spectra, we analyzed the results using ROC (receiver operating characteristic) curves, plotting the detected proteins – ranked by ERM/cytosol enrichment ratio – by true positive rate (TPR) and false positive (FPR) rate at every possible enrichment ratio cut-off. **Figures 3I-K** show that our FlexID dataset was the most specific, better able to retain true positive proteins over false positive proteins than TurboID and APEX2 at every proteome size. The increase in spatial specificity did not come at the expense of sensitivity, as all PL enzymes recovered a similar fraction (∼47%) of well-established ERM proteins (**Figure 3L**). The sensitivity of PL-based mapping is in large part limited by steric access, since labeling is performed in living cells with macromolecular complexes and membranes intact; thus, a considerable fraction of true positive proteins are shielded from labeling by any PL enzyme.

An additional advantage of FlexID apparent from the proteomic data is its temporal precision compared to TurboID. **Extended Data Figure 7L** shows that the ROC curves for TurboID are similar whether or not exogenous biotin is supplied, indicating that significant labeling occurs over ∼20 hours using the endogenous biotin present in culture media (also supported by Western blots in **Figures 3A,F**). Finally, we evaluated whether FlexID’s enhanced spatial specificity could obviate the need for a spatial reference control (e.g., FlexID-NES). The ROC curves based on ERM vs. untransduced enrichment ratio or ERM intensity vs. total protein abundance in **Extended Data Figures 7J-K** show that while APEX2 cannot separate true positives from false positives at all, FlexID can do so, albeit with reduced effectiveness compared to using the ERM/NES enrichment ratio. Thus, the orthogonal substrate utilization and enhanced spatial specificity of FlexID provide distinct benefits over TurboID and APEX2 for spatial proteomic experiments.

### FlexID for high-throughput protein interaction screening in cells

High-throughput evaluation of protein-protein interactions (PPIs) is critical for drug discovery and for understanding the effects of mutations or genetic perturbations (e.g., CRISPRi). Existing PPI detection methods are constrained by the need for dual tagging and overexpression of prey and bait proteins (e.g., NanoBRET), the requirement for protein purification and chemical conjugation (e.g., TR-FRET^32^), or the difficulty of capturing weak or transient interactions (co-immunoprecipitation). PL can address these limitations, but when coupled to mass spectrometry-based readout, it is too expensive and labor-intensive to scale to a large number of conditions. We capitalized on the high spatial resolution and low background of FlexID to develop a high-throughput screening (HTS) method for PPIs based on PL with fluorescence readout instead (“PLF”; **Figure 4A**). In PLF, a bait fused to FlexID catalyzes in-cell PL with BCN probe. After lysis and Click with a fluorescent probe, antibody-coated beads are used to capture endogenous protein targets (“prey”), which will appear fluorescent if the captured proteins were proximal to FlexID during the 5-minute live cell labeling window. The assay requires minimal material (one well of a 384-well plate per condition) and is rapid and inexpensive to perform.

**Figure 4.**
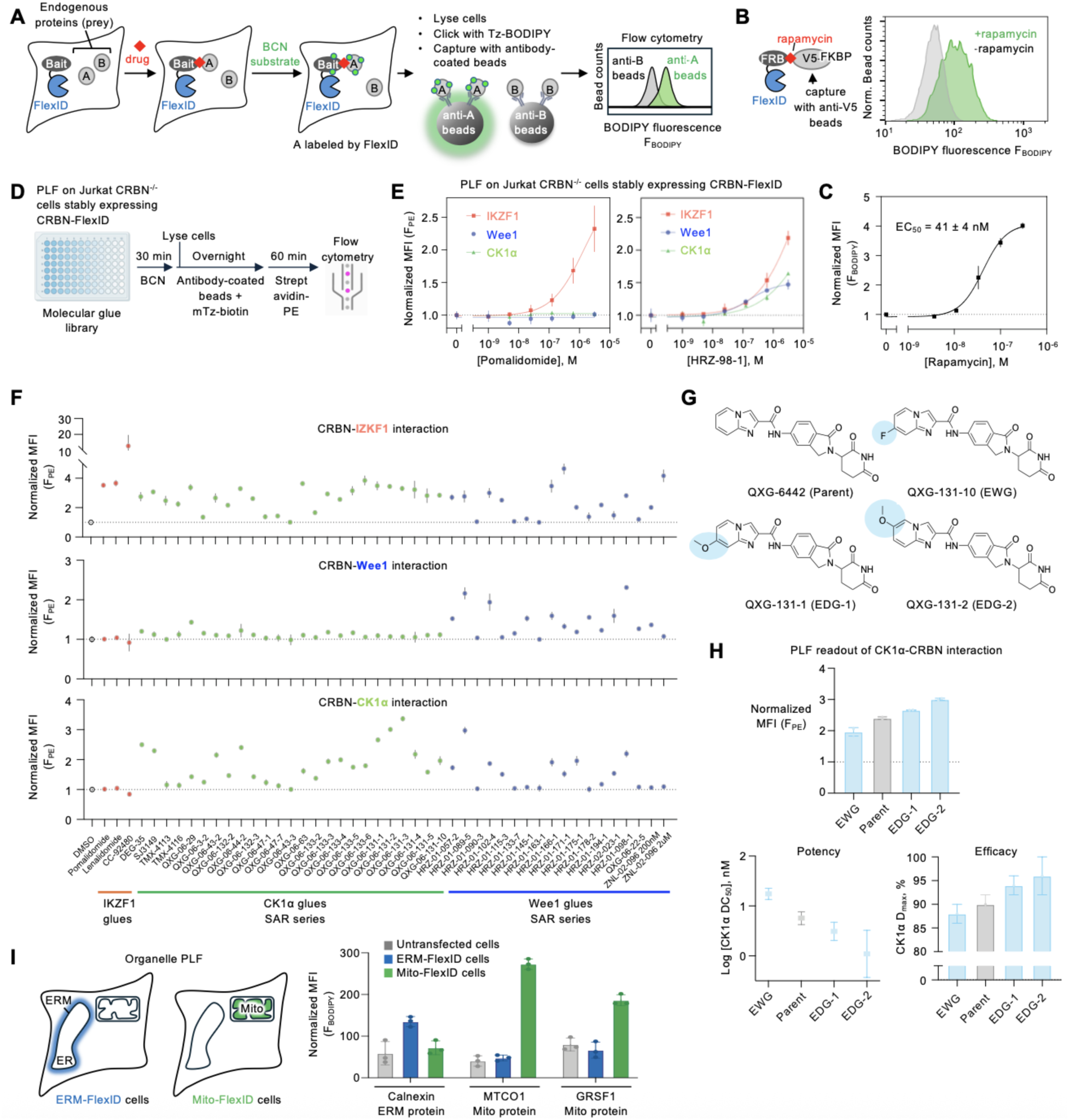
Proximity labeling with fluorescence readout (PLF) for high-throughput screening of protein-protein interactions in living cells. (**A**) PLF schematic. A bait protein is fused to FlexID, which tags endogenous interacting “prey” proteins with BCN. After cell lysis, specific prey proteins are captured with antibody-coated beads, and flow cytometry indicates which ones were proximal to FlexID-bait in living cells (protein A but not B in this example). (**B**) PLF detects rapamycin-induced interaction between FlexID-FRB and V5-FKBP12-APEX. (**C**) Rapamycin dose-response curve for experiment in (B). *N = 2.* Mean ± SD. MFI, mean fluorescence intensity. (**D**) Scheme for high-throughput PLF screen of 46 molecular glues against 3 endogenous neo-substrates in Jurkat cells expressing CRBN-FlexID. mTz-biotin, methyltetrazine-biotin. PE, phycoerythrin. (**E**) PLF dose-response for two molecular glues targeting Ikaros (IKZF1) to Cereblon (CRBN)-FlexID. Beads coated with antibodies against IKZF1, Wee1, and CK1α were used to detect BCN/mTz-biotin/streptavidin-PE-labeled endogenous proteins following the procedure in (D). *N = 2.* Mean ± SD. (**F**) Results from experiment in (D). Beads coated with antibodies against IKZF1 (top), Wee1 (middle), and CK1α (bottom) were used. SAR, structure-activity relationship. *N = 2.* Mean ± SD. (**G**) SAR series based on the molecular glue QXG-06442 and its derivatives with electron withdrawing (EWG) or electron donating (EDG) groups. (**H**) PLF shows that electron donating/withdrawing groups tune the CRBN–CK1α interaction in a manner correlated with glue potency and efficacy. (**I**) PLF can be used to determine the sub-compartment localization of endogenous proteins. HEK 293T cells expressing FlexID targeted to either the ER membrane facing cytosol (ERM) or mitochondrial matrix were labeled with BCN for 5 minutes, then Clicked with tetrazine-BODIPY after lysis. Right: the indicated endogenous proteins are captured with antibody-coated beads and analyzed by flow cytometry as in (A). *N* = 3. Mean ± SD. Background from IgG control beads is subtracted for each sample. Additional data in **Extended Data Fig. 8J-K.**

First, we demonstrated PLF using FRB fused to FlexID. Upon rapamycin treatment, co-transfected V5-tagged FKBP12 was selectively labeled and captured by anti-V5 beads in a dose-dependent manner (**Figures 4B-C**). Next, we used FlexID to profile molecular glue–induced recruitment of endogenous neo-substrates to the E3 ubiquitin ligase Cereblon (CRBN). To preserve physiological expression level, we stably expressed a CRBN–FlexID fusion in CRBN⁻/⁻ Jurkat cells (**Extended Data Figure 8B**). Substrate protein ubiquitination and degradation were inhibited by MLN4924, an inhibitor of Cullin ligase-dependent ubiquitination^33^. To enhance signal-to-noise, BCN-labeled proteins were Clicked with mTz–biotin and stained with streptavidin–phycoerythrin. We first compared target recruitment by two molecular glues with differing specificity profiles: pomalidomide, which selectively recruits IKZF1 to CRBN; and HRZ-98-1, which engages multiple targets including IKZF1, CK1α, and Wee1^34^. Following the PLF workflow, each glue induced the recruitment of expected neo-substrates to CRBN-FlexID in a sensitive and dose-dependent manner (**Figure 4D-E**).

We then expanded our PLF screen to a library of 46 degraders consisting of IKZF1-targeted molecular glues, a CK1α-targeted glue SAR (structure-activity relationship) series, a Wee1-targeted glue SAR series, and a Wee1-directed PROTAC (**Table 5**). The library spans a range of activities (from inactive to potent) and target specificities. We found that the screen was both sensitive and reproducible (**Figures 4F** and **Extended Data Figure 8C**). Moreover, the PPI strength measured by PLF strongly correlated with each degrader’s potency, efficacy, and specificity, as measured by a HiBiT degradation assay^34,35^ (**Extended Data Figures 8D-E**). The large dynamic range of PLF also enabled us to evaluate substitution effects among closely-related degrader derivatives (**Figure 4G-H** and **Extended Data Figures 8H-I**) and discriminate between molecular glue and PROTAC modes of action through visualization of the Hook effect at elevated compound concentrations (**Extended Data Figures 8F**).

Because PLF detects PPIs, it reports on a step upstream of the actual degradation event, in contrast to assays like HiBiT that report on degradation. This fact enabled us to detect, for example, an intrinsic affinity between CRBN and endogenous CK1α^36^ that is diminished by treatment with the potent CRBN binder CC-92480^37^ (**Extended Data Figures 8G**), an observation that would have been invisible to HiBiT. Furthermore, we observed by PLF that several CK1α-targeting glues strongly recruit endogenous IKZF1 to CRBN (**Figure 4F**), despite having been shown to not degrade IKZF1^35^. This observation reinforces the idea that substrate recruitment alone is insufficient to induce target degradation.

Another potential application of PLF is to rapid screening of protein subcellular localization (**Figure 4I**). To test this concept, we targeted FlexID to either the ER membrane facing the cytosol, or to the mitochondrial matrix. After 5-minutes of live cell labeling with BCN, we used antibody-coated beads to enrich the endogenous proteins calnexin, MTCO1, and GRSF1. Calnexin, which resides on the ER membrane, was fluorescently tagged by ERM-FlexID but not mito-FlexID. The mitochondrial matrix proteins MTCO1 and GRSF1, on the other hand, were fluorescently tagged by mito-FlexID but not ERM-FlexID.

These examples illustrate that PLF can be used for rapid, low-cost, and scalable profiling of live cell PPIs and subcellular localization, with high specificity and sensitivity down to endogenous levels.

## Discussion

The power of proximity labeling lies in its spatial resolution and compatibility with living cells, and recent methods using photocatalysts to generate more short-lived reactive intermediates such as carbenes and nitrenes to reduce labeling radius have advanced the field^4,5,38^. FlexID offers a different paradigm for spatial resolution improvement through use of a contact-dependent labeling mechanism in which the reactive intermediate is not released from the enzyme. We showed that this “zero-radius” approach produces more spatially specific proteomes than both APEX and TurboID, while bypassing the high biotin background of TurboID labeling.

Among PL catalysts, PafA ligase which is based on a bacterial form of ubiquitin ligase also catalyzes contact-dependent proteome labeling^39^. However, the ligated “probe” is a peptide that must be genetically expressed in cells, and labeling occurs over many hours to days. FlexID, by contrast, has a temporal resolution of 5 minutes and labeling can be initiated by a variety of small-molecule probes, including bio-orthogonal Click handles.

We were surprised to find in our characterization of FlexID, TurboID, and APEX2 that TurboID showed evidence of a shorter labeling radius than APEX2 in multiple assays. It is possible that TurboID also labels proteins at least partially through a contact-dependent mechanism, contradicting the widespread view that biotin-AMP released from TurboID’s active site is solely responsible for labeling. TurboID lacks the C-terminal domain that gates acceptor protein access in LplA, but possesses a unique N-terminal domain that binds to DNA. Its detailed mechanism will be the subject of a future study. Nevertheless, the background from endogenous biotin and biotinylated proteins hampers TurboID’s utility compared to both FlexID and APEX.

The requirement for direct contact with protein substrates did not appear to diminish FlexID’s ability to capture relevant proteomes compared to APEX and TurboID. This may be because 72.5% of human proteins contain the EXXK motif that FlexID prefers, while 97.3% have a negatively charged sidechain within five residues of lysine. Sensitivity is likely to also be influenced by FlexID’s expression level and copy number within the ER membrane. Since FlexID was not fused to any native ER protein but merely targeted to the ERM via a transmembrane sequence, it is clear that transient, non-cognate protein-protein contacts can be captured by FlexID under the conditions used here. Overall, the speed, spatial specificity, low background, probe versatility, non-toxicity, and *in vivo* compatibility of FlexID make it an attractive alternative to other PL technologies such as APEX and TurboID. However, future improvements in enzyme stability, activity in some compartments (such as the ER lumen), and hydrophobicity of the small-molecule probes will make the technology even more robust.

We used several different computational approaches to engineer and understand FlexID. The protein language model ESM-1b^19^/1v^20^ provided our first breakthrough discovery of promiscuity-enhancing mutations. ESM-predicted mutations also conferred other benefits, including increased expression (F231L, A248S) and reduced aggregation (Y11T, Y11H), likely stemming from loss of a dimerization interface^40^. It is intriguing that the ESM-predicted mutation A248S increased promiscuous activity like A48N, but was not scored as open-biasing; this raises the possibility that A248S increases LplA promiscuity through a different and unknown mechanism. Our findings highlight the power of protein language models to uncover beneficial mutations with diverse functional effects. We also used ProteinMPNN^10^ in this study, which helped to identify expression-enhancing mutations. However, using ProteinMPNN to redesign the active site shell of LplA mostly created instability, evident from early plateauing of flow cytometry curves (**Supplementary data 1**). These redesigned sequences tended to change highly-conserved residues in the hydrophobic substrate binding pocket. Finally, Conformational Biasing^18^ provided a mechanistic explanation for FlexID’s enhanced promiscuity, likely conferred through stabilization of the open conformation that has reduced K_m_ for non-cognate protein substrates. Our results showcase the complementary strengths of computational tools that are trained on different datasets and customized for different functions.

FlexID enabled the development of PLF (proximity labeling with fluorescence readout), a more targeted and vastly more scalable version of PL for PPI detection that offers the same benefits of live cell compatibility, high spatial resolution, rapid tagging, and ability to capture endogenous prey. The large dynamic range in PLF observed with FlexID would have been difficult to achieve with lower-resolution or higher-background PL enzymes like APEX and TurboID. We showed that PLF is suitable for screening a specific set of PPIs across a large number of conditions. For evaluation of molecular glue degraders, PLF is distinguished from the widely-used HiBiT^34,35^ assay due to its gain-of-signal format and mechanistic-agonistic identification of compounds that directly engage target proteins. However, PLF’s general ability to report on PPIs should make it applicable to many other drug classes including PPI modulators that act through non-degradative mechanisms (e.g., trametiglue^41^, immunophilin recruiters^42,43^, and transcriptional/epigenetic CIPs^45^).

## Acknowledgements

A.Y.T. is grateful for funding from the GPCR collaborative of the St. Jude Children’s Research Hospital, CIRM Center for Neuropsychiatric Stem Cell Proteomics, NIH (RC2DK129964), Chan Zuckerberg Biohub – San Francisco, Phil and Penny Knight Initiative for Brain Resilience, and Stanford Bio-X. P.E.C., A.G.X. and M.A. are both supported by NSF Graduate Research Fellowships. A.G.X is also supported by NIH T32GM136568. S.A.D. is supported by Schmidt Science Fellows (in partnership with the Rhodes Trust) and NIH K99CA297018. W.Q. is grateful for funding from the National Key Research and Development Program of China (No. 2024YFA1308000), the National Natural Science Foundation of China (22477066 and 92478128), Beijing Natural Science Foundation (JQ25018), the Fundamental Research Funds from Beijing National Laboratory for Molecular Sciences (BNLMS202301) and the Shenzhen Medical Research Fund (B2401004). Use of the Stanford Synchrotron Radiation Lightsource, SLAC National Accelerator Laboratory, is supported by the U.S. Department of Energy, Office of Science, Office of Basic Energy Sciences under Contract No. DE-AC02-76SF00515. The SSRL Structural Molecular Biology Program is supported by the DOE Office of Biological and Environmental Research, and by the National Institutes of Health, National Institute of General Medical Sciences (P30GM133894).

We thank Kathy Le for help with yeast cytosolic extracts, Jay Kittler and Dan Schwartz for pLogo, Woong Sub Byun for sharing CRBN^-/-^ Jurkat cells, and Hlib Razumkov for sharing Wee1-targeting molecular glues. Charles Xu, Matt Ravalin, Derek Tan, Zhe Zhuang, Tangpo Yang, Haoyuan Wang, Zixuan Jiang, Dain Brademan, Chang Lin, Chu Zheng, Varun Shanker, Mateo Sanchez Lopez, Shuo Han, Reika Tei, Joseph Fox, and Kelvin Cho provided valuable guidance and discussion. We also thank Onn Brandman, Dan Herschlag, Pehr Harbury, Wei Wei, and Jon Long for useful discussions.

## Author contributions

A.Y.T. and P.E.C. conceived of FlexID. A.Y.T., S.A.D., and P.E.C. conceived the overall project. P.E.C. engineered FlexID with help from S.A.D. and A.Q. P.E.C. and B.L.H. designed the ESM library. A.G.X. and P.E.C. designed ProteinMPNN libraries. S.J.P. and P.E.C. used ProteinMPNN to redesign LplA’s active site. S.A.D. synthesized LplA ligands. S.A.D. and P.E.C. performed Trp fluorescence and yeast lysate labeling experiments. P.E.C. and T.M. performed SEC-SAXS and A.G.X. analyzed the data. S.A.D. performed BLI and DSF assays. A.Q. characterized FlexID for proximity labeling with guidance from S.A.D. and P.E.C. S.A.D., P.E.C. and A.Y.T. designed proteomic experiments, S.A.D. and P.E.C. prepared proteomic samples, Z.Z. processed proteomic samples, and S.A.D., P.E.C., and Z.Z. analyzed proteomic data. K.N., N.D.U., S.A.D., and P.E.C. performed preliminary proteomic studies. M.A. and S.A.D. performed mouse experiments. S.A.D. performed neuron culture experiments. S.A.D. performed PLF experiments with help from A.Q. and Q.G. W.Q., C.K., N. G. and A.Y.T. provided supervision. A.Y.T, S.A.D, and P.E.C. wrote the manuscript with input from A.G.X.

## Data availability

Source data for Figure 1H-I are provided in the paper in Extended Data Table 1. Source Data for Source data for Figure 2C-D are provided in the paper in Extended Data Table 6. Source Data for Figure 2M are provided in the paper in Extended Data Table 2. Source data for Figure 3I-L and Extended Data Figure 7C-L are provided in the paper in Extended Data Table 4. Source data for Figure 4F-H and Extended Data Figure 8C-I are provided in the paper in Extended Data Table 5. The original mass spectra may be downloaded from MassIVE (http://massive.ucsd.edu) after publication. Supplementary 2D flow cytometry data and SEC-SAXS data for LplA variants are available for download at Zenodo at: https://doi.org/10.5281/zenodo.17583062. Any additional data that support the findings of this study are available from the corresponding author upon reasonable request.

