## Supplementary Information for "Computational design of a versatile, zero-radius proximity labeling enzyme"

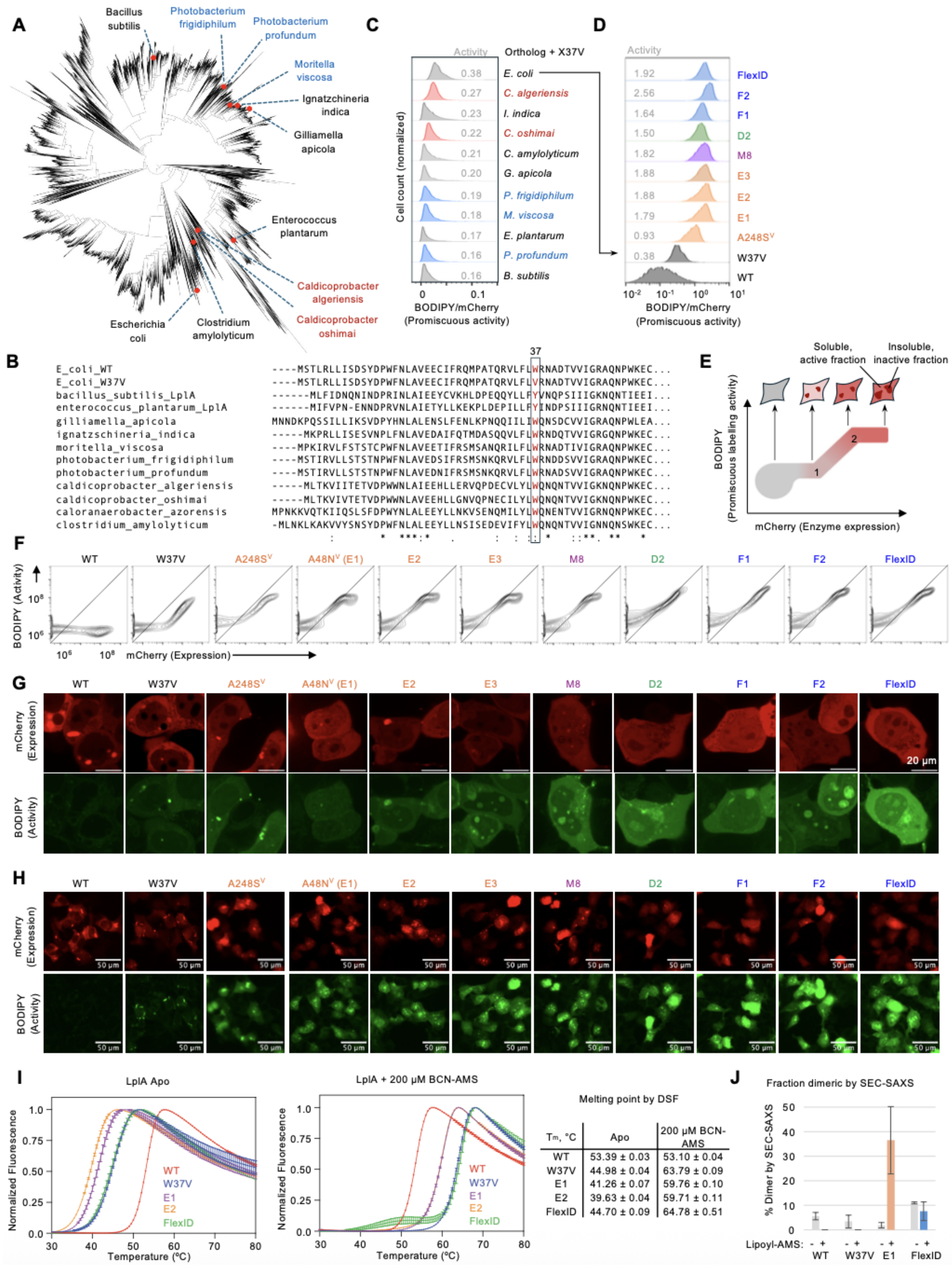


| Tm, °C | Apo | 200 µM BCN-AMS |
|---|---|---|
| WT | 53.39 ± 0.03 | 53.10 ± 0.04 |
| W37V | 44.98 ± 0.04 | 63.79 ± 0.09 |
| E1 | 41.26 ± 0.07 | 59.76 ± 0.10 |
| E2 | 39.63 ± 0.04 | 59.71 ± 0.11 |
| FlexID | 44.70 ± 0.09 | 64.78 ± 0.51 |

1

**Extended Data Figure 1. Engineering FlexID** (related to Figure 1)**.** (**A**) Bacterial phylogenetic tree, highlighting LplA orthologs tested. Thermophiles (red), cryophiles (blue), and mesophiles (black) were selected. (**B**) Introduction of X37V mutation into all LplA orthologs by sequence alignment, in order to allow accommodation of BCN substrate. (**C**) Promiscuous activities of LplA orthologs from (B). Labeling was performed with BCN for 5 minutes as in **Figure 1G**. (**D**) 1D Flow cytometry histograms for W37V, FlexID, and engineering intermediates. This experiment was performed 3 times with similar results. (**E**) Interpretation of 2D flow cytometry plots from assay in **Figure 1G**. BODIPY saturation at high LplA expression level likely reflects enzyme aggregation, which limits promiscuous labeling extent. (**F**) 2D flow cytometry histograms for samples in (D). (**G**) Live-cell confocal microscopy of the same LplA variants. This is an expanded version of **Figure 1J**. This experiment was performed 2 times with similar results. (**H**) Same as (G) but with 20x magnification to visualize more cells in each field of view. (**I**) Melting temperatures ($T_m$) for LplA variants measured via differential scanning fluorimetry (DSF) in the apo state (left) or bound to BCN-AMS (right). $N = 2$. Mean ± SD. (**J**) Dimerization propensity of LplA variants measured by SEC-SAXS. Details in **Extended Data Figure 5**. $N = 2\text{-}5$. Mean ± SD. Sequences of all LplA variants are listed in **Table 1**.

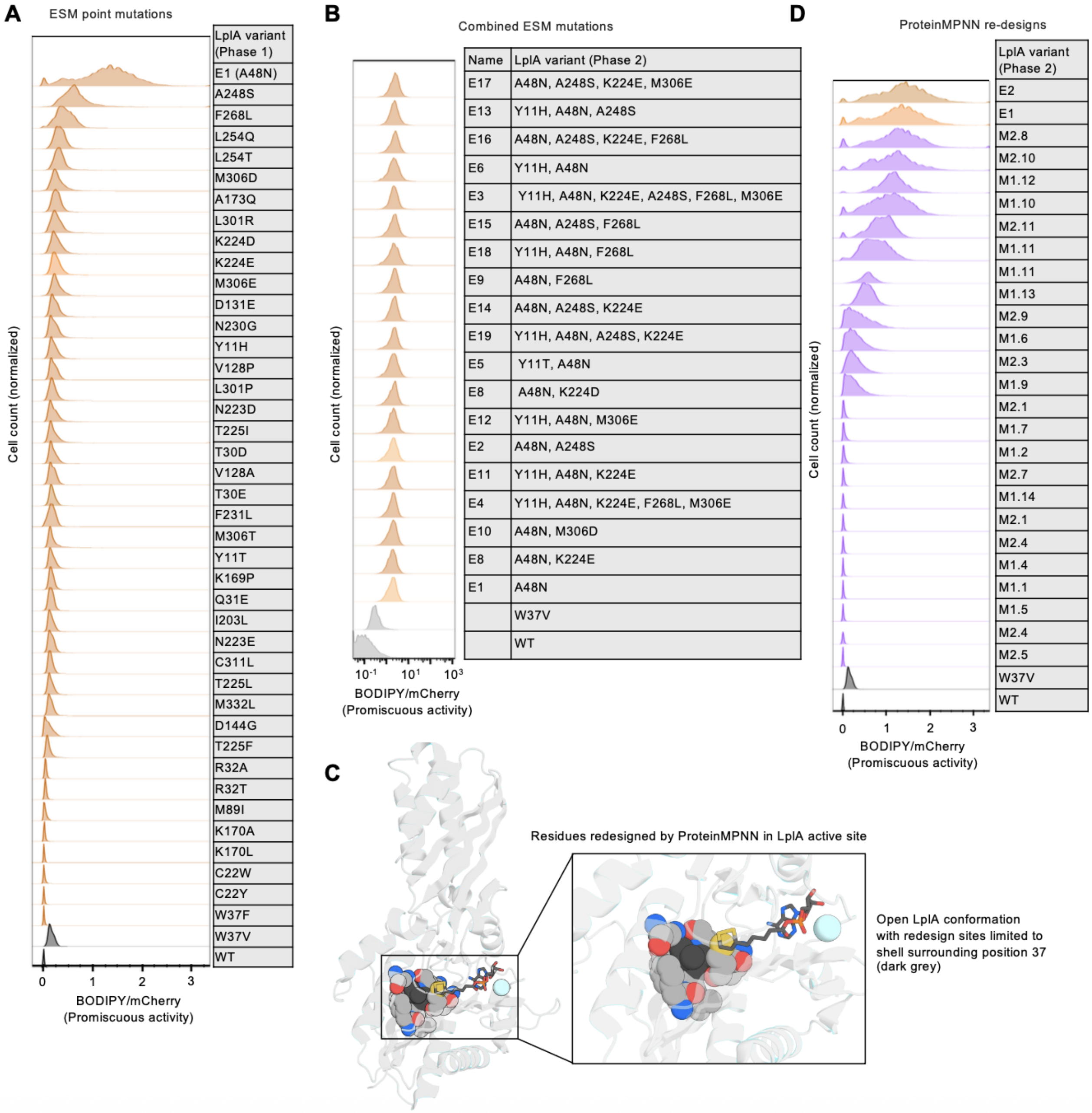
A
ESM point mutations
LpIA variant (Phase 1)
E1 (A48N)
A248S
F268L
L254Q
L254T
M306D
A173Q
L301R
K224D
K224E
M306E
D131E
N230G
Y11H
V128P
L301P
N223D
T225I
T30D
V128A
T30E
F231L
M306T
Y11T
K169P
Q31E
I203L
N223E
C311L
T225L
M332L
D144G
T225F
R32A
R32T
M89I
K170A
K170L
C22W
C22Y
W37F
W37V
WT
Cell count (normalized)
0 1 2 3
BODIPY/mCherry
(Promiscuous activity)
B
Combined ESM mutations
Name | LpIA variant (Phase 2)
E17 | A48N, A248S, K224E, M306E
E13 | Y11H, A48N, A248S
E16 | A48N, A248S, K224E, F268L
E6 | Y11H, A48N
E3 | Y11H, A48N, K224E, A248S, F268L, M306E
E15 | A48N, A248S, F268L
E18 | Y11H, A48N, F268L
E9 | A48N, F268L
E14 | A48N, A248S, K224E
E19 | Y11H, A48N, A248S, K224E
E5 | Y11T, A48N
E8 | A48N, K224D
E12 | Y11H, A48N, M306E
E2 | A48N, A248S
E11 | Y11H, A48N, K224E
E4 | Y11H, A48N, K224E, F268L, M306E
E10 | A48N, M306D
E8 | A48N, K224E
E1 | A48N
W37V
WT
Cell count (normalized)
10⁻¹ 10¹ 10³
BODIPY/mCherry
(Promiscuous activity)
C
Residues redesigned by ProteinMPNN in LpIA active site
Open LpIA conformation
with redesign sites limited to
shell surrounding position 37
(dark grey)
D
ProteinMPNN re-designs
LpIA variant (Phase 2)
E2
E1
M2.8
M2.10
M1.12
M1.10
M2.11
M1.11
M1.11
M1.13
M2.9
M1.6
M2.3
M1.9
M2.1
M1.7
M1.2
M2.7
M1.14
M2.1
M2.4
M1.4
M1.1
M1.5
M2.4
M2.5
W37V
WT
Cell count (normalized)
0 1 2 3
BODIPY/mCherry
(Promiscuous activity)

**E**

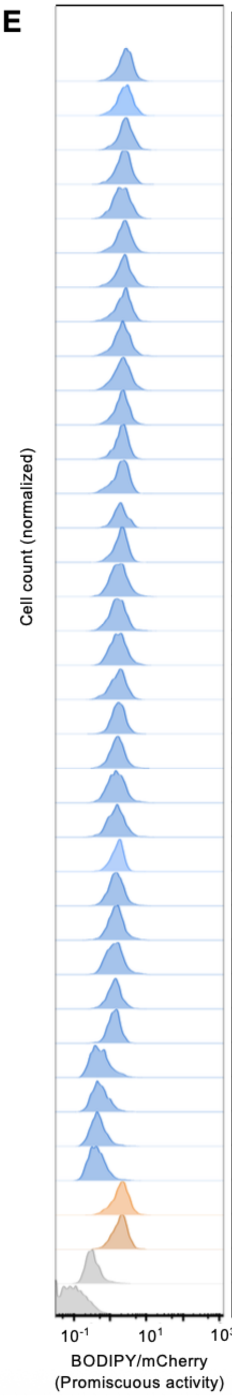


| Name | LplA variant (Phase 3) |
|---|---|
| F7 | Y11H, A48N, F147L, K224E, H267R, M306E |
| F5 | Y11H, F35L, A48N, F147L, K224E, H267R, M306E |
| F4 | Y11H, A48N, F147L, K224E, A248S, H267R, M306E |
| F2 | F35L, A48N, F147L, K224E, M306E |
| F13 | Y11H, F35L, A48N, T57I, F147L, K224E, A248S, H267R, F268L, M306E |
| F6 | Y11H, F35L, A48N, F147L, K224E, A248S, H267R, M306E |
| F27 | F35L, W37V, A48N, T84L, F147L, K224E, A248S, H267R, M306E |
| F28 | F35L, W37V, A48N, F147L, K224E, H267R, M306E |
| F19 | F35L, W37V, A48N, F147L, K224E, A248S, H267R, M306E |
| FlexID | Y11H, A48N, T571, F147L, K224E, A248S, H267R, M306E |
| F28 | F35L, A48N, F147L, K224E, H267R, F268L, M306E |
| F10 | F35L, A48N, T84L, F147L, K224E, A248S, H267R, F268L, M306E |
| F21 | F35L, W37V, F147L, K224E, A248S, H267R, M306E |
| F9 | Y11H, A48N, T57I, F147L, K224E, A248S, H267R, F268L, M306E |
| E20 | Y11H, A48N, K224E, F268L, M306E |
| F3 | Y11H, F35L, A48N, T57I, F147L, K224E, H267R, M306E |
| F1 | Y11H, F35L, A48N, T57I, F147L, K224E, A248S, H267R, M306E |
| F16 | S8L, Y11H, F35L, W37V, A48N, T57I, T84L, F147L, K224E, A248S, H267R, M306E |
| F20 | Y11H, F35L, W37V, A48N, T57I, F147L, K224E, H267R, M306E |
| F30 | F35L, W37V, A48N, K224E, H267R, M306E |
| F22 | S8L, Y11H, F35L, W37A, A48N, T571, T84L, F147L, K224E, A248S, H267R, M306E |
| F8 | Y11H, F35L, A48N, T57I, F147L, K224E, H267R, F268L, M306E |
| F26 | Y11H, F35L, W37V, A48N, T571, F147L, K224E, A248S, H267R, M306E |
| F29 | S8L, Y11H, F35L, W37A, A48N, T84L, K224E, A248S, F268L, M306E |
| F12 | S8L, Y11H, F35L, W37A, A48N, TB4L, F147L, K224E, A248S, H267R, F268L, M306E |
| F24 | S8L, Y11H, F35L, W37A, A48N, T84L, K224E, H267R, M306E |
| D2 | A48N, T571, F147L, A248S, H267R |
| F23 | S8L, Y11H, F35L, W37A, A48N, T57I, T84L, F147L, K224E, H267R, M306E |
| F17 | S8L, Y11H, F35L, W37A, A48N, T84L, K224E, A248S, H267R, M306E |
| F15 | Y11H, F35L, W37V, A48N, T571, K224E, M306E |
| F25 | Y11H, W37A, A48N, T57I, F147L, K224E, A348S, H267R, M306E |
| F25 | Y11H, W37A, A48N, T57I, F147L, K224E, A248S, H267R, M306E |
| F18 | S8L, Y11H, F35L, W37A, A48N, T57I, T84L, K224E, M306E |
| E2 | A48N, A248S |
| E1 | A48N |
| W37V | W37V |
| WT | WT |

**Extended Data Figure 2. Flow cytometry data for complete set of ESM-designed, ProteinMPNN-designed, and combination LplA mutants.** (**A**) Flow cytometry histograms for all ESM-designed LplA point mutations (Phase 1 of engineering). Labeling was performed as in

**Figure 1G** with 5 minutes of BCN. (**B**) Flow cytometry histograms of all combined ESM mutations (from Phase 2 of engineering). (**C**) Pymol rendering of LplA in open conformation (PBD 3A7R[1]), with residues selected for ProteinMPNN redesign shown as grey spheres (W37V mutation is shown in black). (**D**) Flow cytometry histograms of all ProteinMPNN redesigns, along with the starting templates for these designs, E1 and E2 (from Phase 2 of engineering). (**E**) Flow cytometry histograms of all combination mutants from Phase 3 of FlexID engineering, along with the starting templates WT, W37V, E1, and E2. All 2D flow cytometry plots shown in **Supplementary Data 1**.

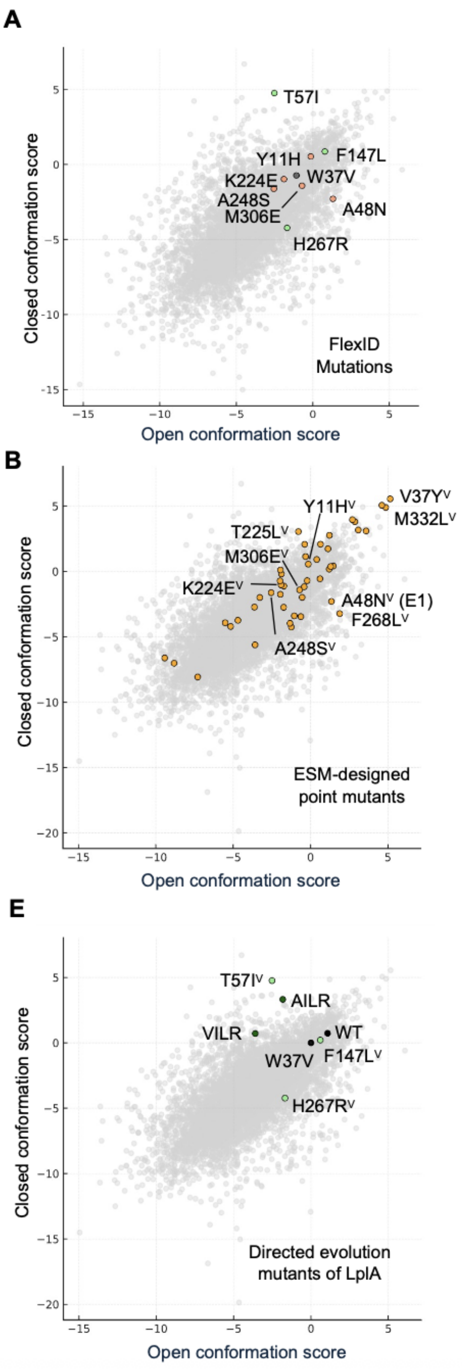


**Extended Data Figure 3. Conformational Biasing (CB)$^2$ analysis of LplA variants.** (A) CB scores for the 9 mutations in FlexID. ESM-derived mutations shown in orange and DE-derived mutations shown in green. Closed and open conformation scores based on ProteinMPNN, using the closed and open structures of LplA (PDB 1X2G[1] and 3A7R[3], respectively). (**B**) CB scores for FlexID and LplA engineering intermediates. Sequences of variants in **Table 1**. (**C**) CB scores for ESM-predicted point mutants of LplA. (**D**) CB scores for ProteinMPNN active site redesigns. The most active designs, M2.8 and M2.10, are highlighted. (**E**) CB scores for directed evolution mutants of LplA.

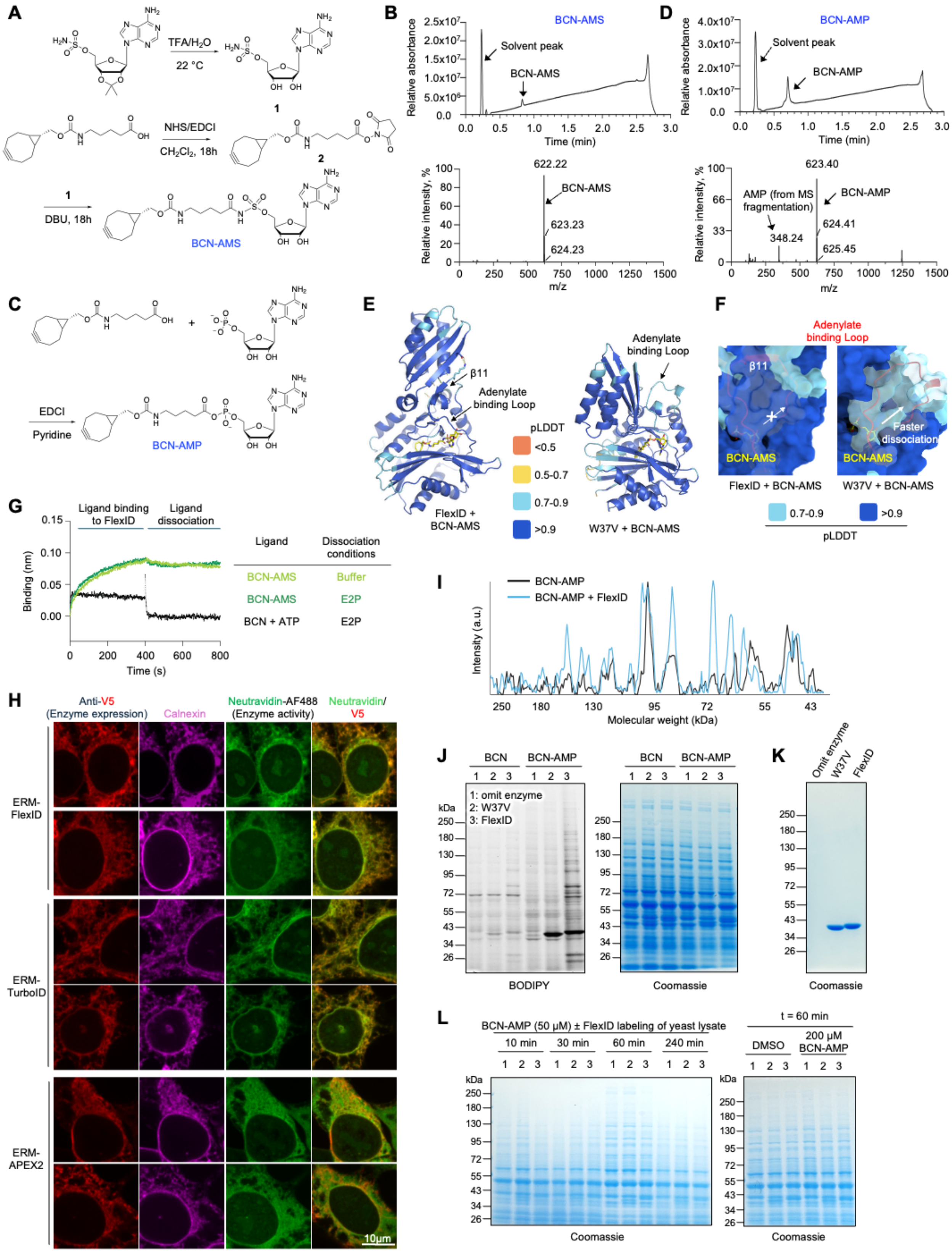
A
TFA/H2O
22 °C
1
NHS/EDCI
CH2Cl2, 18h
2
DBU, 18h
BCN-AMS
B
BCN-AMS
Solvent peak
BCN-AMS
Relative absorbance
Time (min)
622.22
623.23
624.23
Relative intensity, %
m/z
C
EDCI
Pyridine
BCN-AMP
D
BCN-AMP
Solvent peak
BCN-AMP
Relative absorbance
Time (min)
623.40
AMP (from MS fragmentation)
348.24
624.41
625.45
Relative intensity, %
m/z
E
β11
Adenylate binding Loop
pLDDT
<0.5
0.5-0.7
0.7-0.9
>0.9
FlexID + BCN-AMS
W37V + BCN-AMS
F
Adenylate binding Loop
β11
BCN-AMS
Faster dissociation
FlexID + BCN-AMS
W37V + BCN-AMS
0.7-0.9
>0.9
pLDDT
G
Ligand binding to FlexID
Ligand dissociation
Binding (nm)
Time (s)
Ligand
Dissociation conditions
BCN-AMS
Buffer
BCN-AMS
E2P
BCN + ATP
E2P
H
Anti-V5 (Enzyme expression)
Calnexin
Neutravidin-AF488 (Enzyme activity)
Neutravidin/ V5
ERM-FlexID
ERM-TurboID
ERM-APEX2
10μm
I
BCN-AMP
BCN-AMP + FlexID
Intensity (a.u.)
Molecular weight (kDa)
J
BCN
BCN-AMP
1 2 3
1: omit enzyme
2: W37V
3: FlexID
kDa
BODIPY
Coomassie
K
Omit enzyme
W37V
FlexID
Coomassie
L
BCN-AMP (50 μM) ± FlexID labeling of yeast lysate
10 min
30 min
60 min
240 min
t = 60 min
DMSO
200 μM BCN-AMP
kDa
Coomassie
Coomassie

1

**Extended Data Figure 4. FlexID characterization and contact-dependent mechanism** (related to Figure 2)**.** (**A**) Synthesis of BCN-AMS. TFA, Trifluoroacetic acid. NHS, N-Hydroxysuccinimide. EDCI, 1-Ethyl-3-(3-dimethylaminopropyl)carbodiimide. DBU, 1,8-Diazabicyclo(5.4.0)undec-7-ene. (**B**) LC-MS of purified BCN-AMS. MS (ESI) for $C_{26}H_{36}N_7O_9S$ [M + H]+: m/z calculated, 622.23; found, 622.22. (**C**) Synthesis of BCN-AMP. (**D**) LC-MS of purified BCN-AMP. MS (ESI) for $C_{26}H_{36}N_6O_{10}P$[M + H]+: m/z calculated, 623.22; found, 623.40. (**E**) Boltz-2-predicted structures of FlexID and W37V LplA in complex with BCN-AMS. (**F**) Close-up view of the predicted adenylate binding pocket in both FlexID and W37V LplA. In FlexID, the exit route of BCN-AMS is blocked by a stabilized adenylate-binding loop, whereas in W37V, the same loop is partially disordered. (**G**) Development of BLI (Bio-Layer Interferometry) assay for measuring ligand dissociation from FlexID. E2P is a natural acceptor protein for wild-type LplA. This experiment was performed once. (**H**) Additional fields of view from experiment in **Figure 2G**. (**I**) Line-scan analysis from lysate labeling experiment in **Figure 2K** (for $4^{th}$ and $6^{th}$ lanes on gel at right), showing distinct banding patterns for proteins chemically vs. enzymatically labeled by BCN-AMP. (**J**) Control for experiment in **Figure 2K** showing that pre-mixing 1 mM lipoyl-AMS with lysate prevents added enzyme from using hydrolyzed BCN acid to complete BCN labeling. This experiment was performed twice with similar results. (**K**) Coomassie-stained gels of purified W37V LplA and FlexID used for experiment in **Figure 2K**. (**L**) Coomassie-stained gels from experiment in **Figure 2K**.

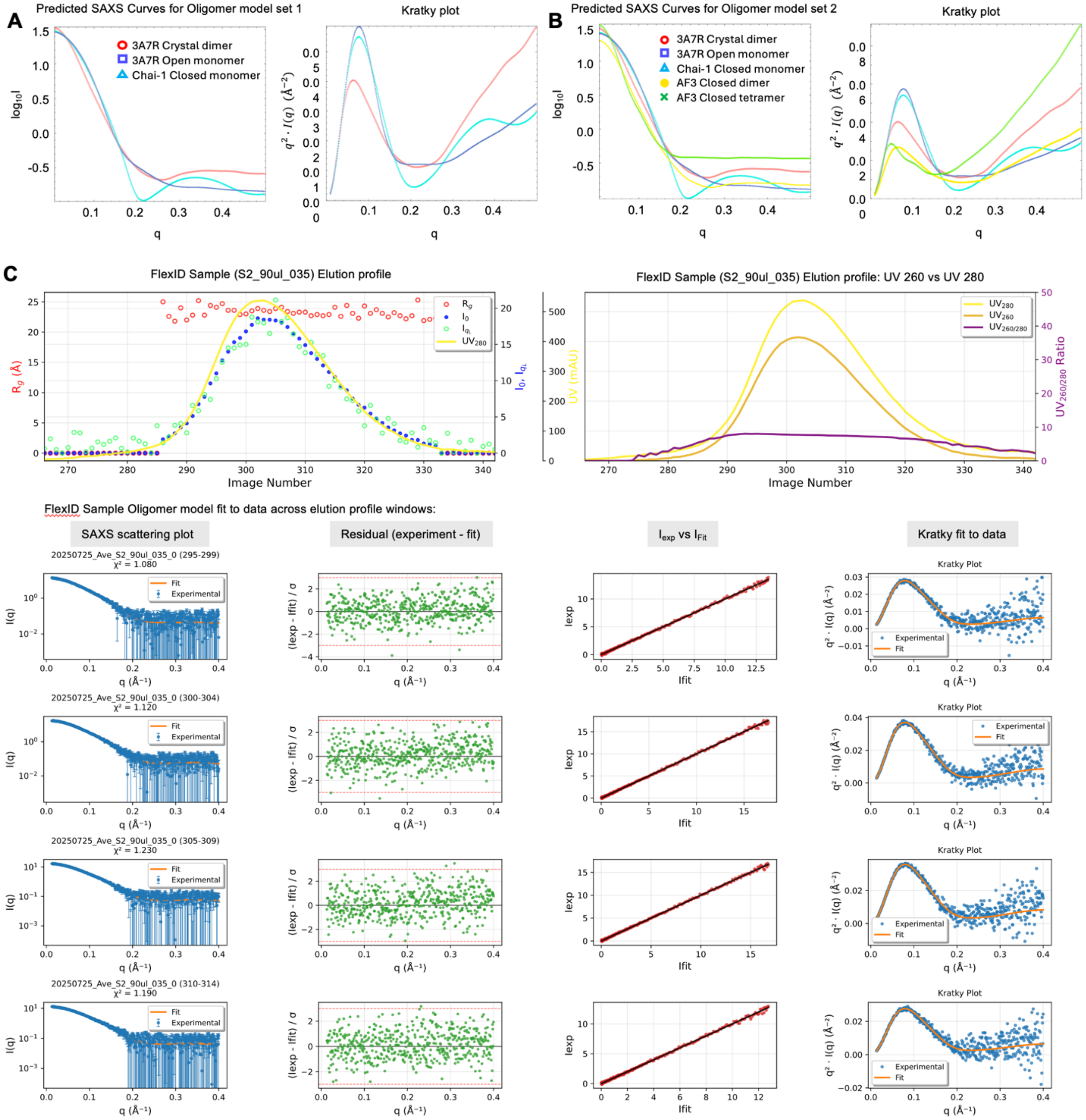


**Extended Data Figure 5. SEC-SAXS analysis of LplA.** (**A**) Superposition of the predicted SAXS scattering curves for the three major LplA species expected: open monomer (from PDB ID: 3A7R), closed monomer (from Chai-1 prediction), and open dimer (from PDB 3A7R[1] crystal lattice). Both Log(I) vs q and Kratky representations are shown. (**B**) Same as (A), but additional predicted SAXS scattering curves for closed tetramer and closed dimer (both based on AlphaFold3 prediction) are included. (**C**) Example SEC-SAXS data for FlexID + lipoyl-AMS. Top shows SEC elution profile with $I_O$, $I_q$, $R_g$ plotted at left, and $UV_{280}$, $UV_{260}$, and $UV_{260/280}$ at right. SAXS scattering curves are shown at bottom left for different windows of analysis, overlaid with Oligomer fits to data. For each analysis window, residual (data – model) vs q plots, $I_{Exp}$ vs $I_{Fit}$ plots, and Kratky plots are given. Complete SAXS data for other LplA variants provided in **Supplementary Data 2**.

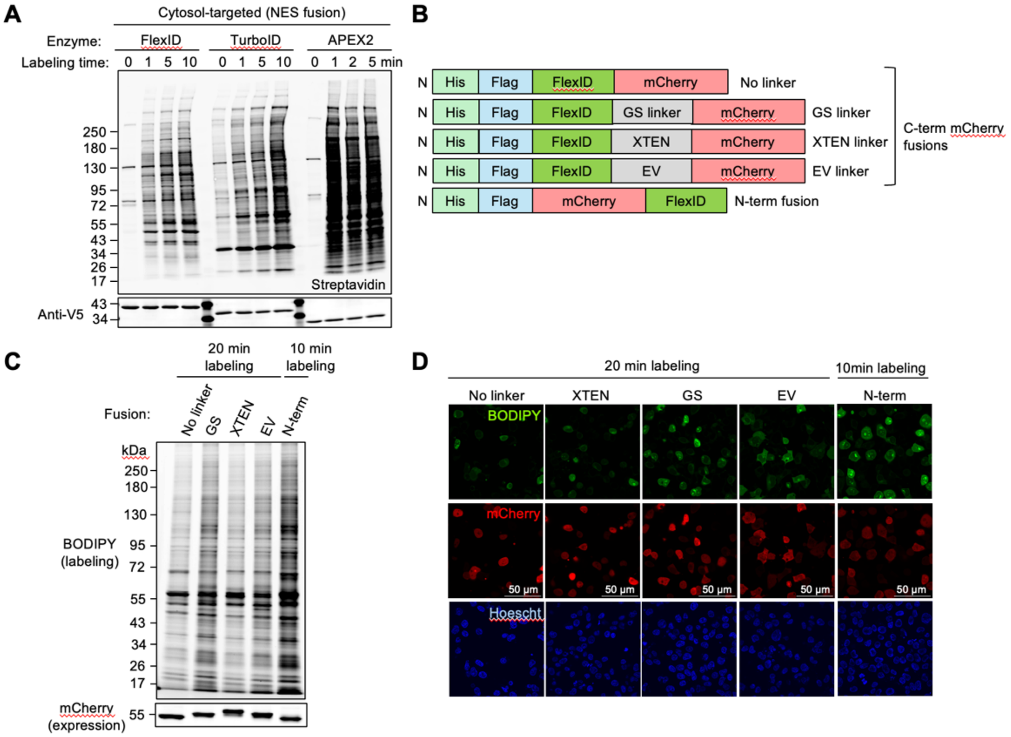


**Extended Data Figure 6. FlexID time course and C-terminal tagging** (related to Figure 3)**.** (**A**) Time course as in Figure 3A but using PL enzymes targeted to the cytosol (via NES) rather than ERM. This experiment was performed twice with similar results. (**B**) Domain structures of different FlexID-mCherry fusions tested in (C) and (D). (**C**) Comparison of FlexID fusions in (B). Transiently transfected HEK 293T cells were labeled with BCN for 10 or 20 minutes, then lysed. Lysates were Clicked with mTz-BODIPY and analyzed by in-gel fluorescence. This experiment was performed 3 times with similar results. (**D**) Confocal microscopy comparing FlexID fusions in (B). HEK 293T cells prepared as in (C) were treated live with 200 nM mTz-BODIPY, stained with Hoescht, and imaged live. This experiment was performed 3 times with similar results.

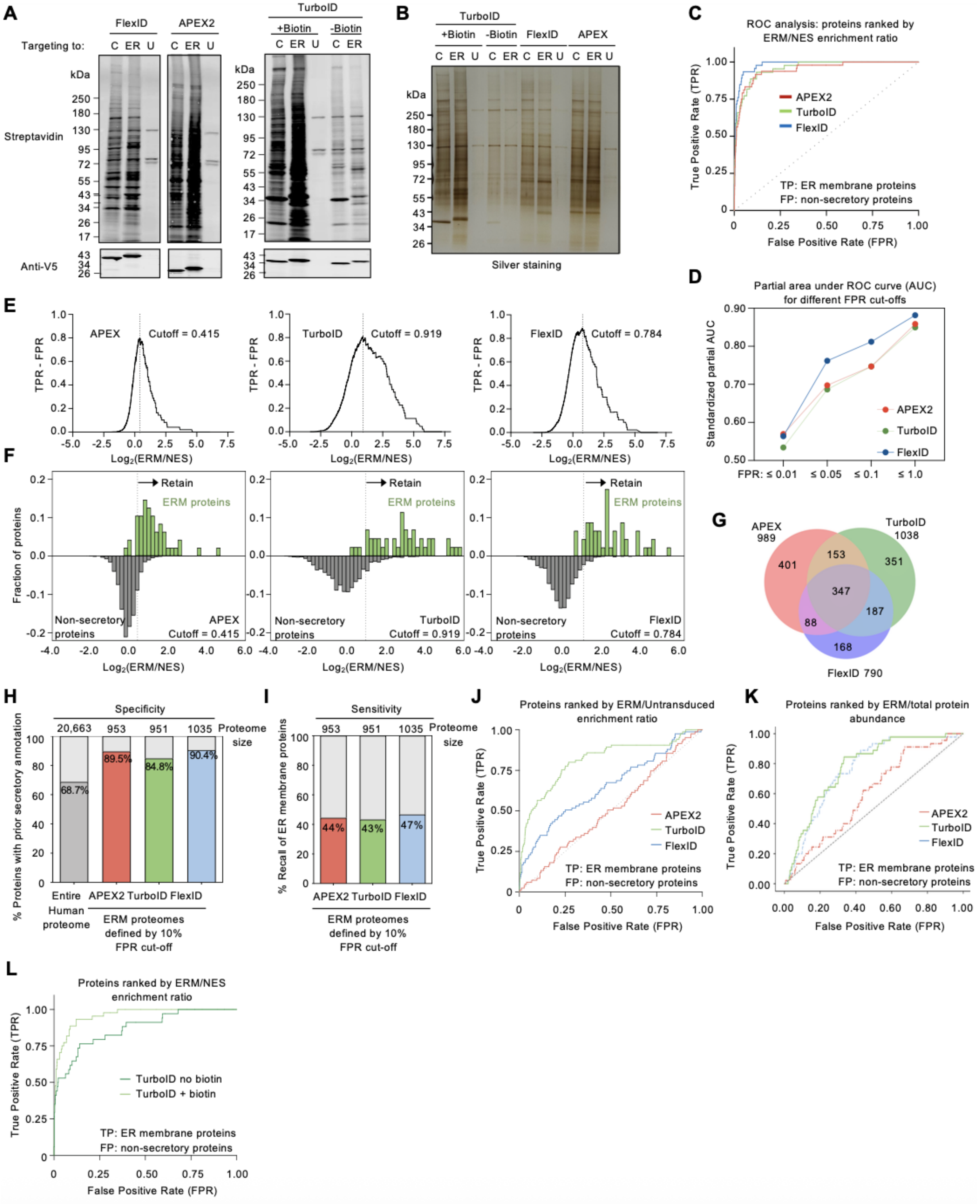


**Extended Data Figure 7. FlexID 5-minute proteomic experiment** (related to Figure 3)**.** (**A**) Streptavidin blots of whole cell lysates from proteomic experiment in **Figure 3H**. Anti-V5 blot detects PL enzyme expression. C: cytosolic/NES. ER: ER membrane. U: Untransduced. (**B**) Silver stained gel of streptavidin-enriched proteomic samples from **Figure 3H**. (**C**) Expanded version of ERM/NES ROC curve from **Figure 3I**, showing FPR up to 1.0. (**D**) Partial area under ROC curve in (C), at different FPR thresholds. (**E**) Determination of ROC-based cut-offs for APEX2, TurboID, and FlexID 5-minute ERM proteomes. Cutoff is made at the log2 ERM/NES intensity ratio at which TPR-FPR is maximal. (**F**) Histograms showing the distribution of true positives (green; proteins with prior ER membrane annotation, **Table 3**) and false positives (grey; non-secretory proteins that should not be labeled by ERM-ligase fusions, **Table 3**) by log2 ERM/NES intensity ratio. Dashed vertical lines show ROC-based cut-offs from (E). (**G**) Venn diagram showing overlap of FlexID 5-minute, TurboID 5-minute, and APEX2 1-minute ERM proteomes from **Figures 3H-L**. (**H-I**) Alternative proteomes based on 10% FPR cut-off instead of ROC-based cut-off (see Methods). The same mass spectrometry data was filtered according to FPR only. Specificity (H) and sensitivity (I) calculated as in **Figures 3K-L**. (**J-K**) Alternative ROC curves based on ERM/untransduced enrichment ratio (J) or ERM/total protein abundance ratio (K) instead of ERM/NES enrichment ratio as in **Figure 3I**. Without a spatial reference, APEX cannot enrich true positives over false positives. TurboID specificity is enhanced by its ability to label proteins with endogenous biotin over 20 hours of expression. FlexID labeling is restricted to a 5-minute window. In (K), HEK lysate DIA proteomic data[4] was used as the denominator to normalize for differences in total protein abundance. (**L**) ERM/NES ROC curves comparing TurboID-enriched proteins in the presence vs. absence of added biotin.

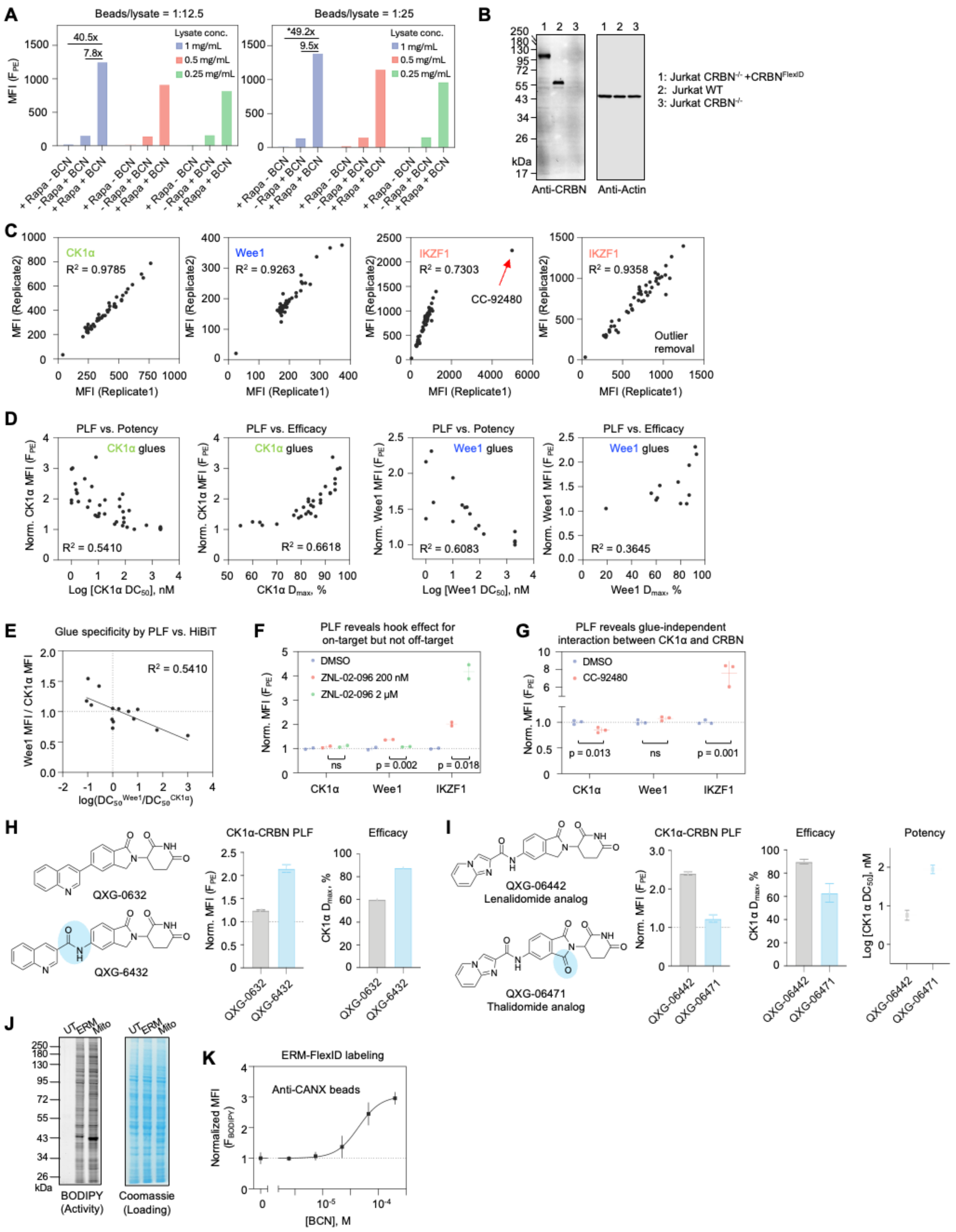
A
Beads/lysate = 1:12.5
Beads/lysate = 1:25
MFI (F_PE)
Lysate conc.
1 mg/mL
0.5 mg/mL
0.25 mg/mL
40.5x
7.8x
*49.2x
9.5x
+ Rapa - BCN
- Rapa + BCN
+ Rapa + BCN
B
1: Jurkat CRBN-/- +CRBNFlexID
2: Jurkat WT
3: Jurkat CRBN-/-
kDa
Anti-CRBN
Anti-Actin
C
CK1α
R2 = 0.9785
Wee1
R2 = 0.9263
IKZF1
R2 = 0.7303
CC-92480
IKZF1
R2 = 0.9358
Outlier removal
MFI (Replicate1)
MFI (Replicate2)
D
PLF vs. Potency
PLF vs. Efficacy
CK1α glues
Wee1 glues
R2 = 0.5410
R2 = 0.6618
R2 = 0.6083
R2 = 0.3645
Norm. CK1α MFI (F_PE)
Norm. Wee1 MFI (F_PE)
Log [CK1α DC50], nM
CK1α Dmax, %
Log [Wee1 DC50], nM
Wee1 Dmax, %
E
Glue specificity by PLF vs. HiBiT
R2 = 0.5410
Wee1 MFI / CK1α MFI
log(DC50Wee1/DC50CK1α)
F
PLF reveals hook effect for on-target but not off-target
DMSO
ZNL-02-096 200 nM
ZNL-02-096 2 μM
Norm. MFI (F_PE)
ns
p = 0.002
p = 0.018
CK1α
Wee1
IKZF1
G
PLF reveals glue-independent interaction between CK1α and CRBN
DMSO
CC-92480
Norm. MFI (F_PE)
p = 0.013
ns
p = 0.001
CK1α
Wee1
IKZF1
H
QXG-0632
QXG-6432
CK1α-CRBN PLF
Efficacy
Norm. MFI (F_PE)
CK1α Dmax, %
I
QXG-06442
Lenalidomide analog
QXG-06471
Thalidomide analog
CK1α-CRBN PLF
Efficacy
Potency
Norm. MFI (F_PE)
CK1α Dmax, %
Log [CK1α DC50], nM
J
UT
ERM
Mito
kDa
BODIPY
(Activity)
Coomassie
(Loading)
K
ERM-FlexID labeling
Anti-CANX beads
Normalized MFI (F_BODIPY)
[BCN], M

1

**Extended Data Figure 8. Additional data related to PLF experiments** (related to Figure 4)**.** (**A**) Optimization of PLF assay using FKBP/FRB constructs from **Figure 4B**. BCN-labeled proteins were Clicked with mTz-biotin, captured on anti-V5 antibody-coated beads, and detected by staining with streptavidin-phycoerythrin (PE). (**B**) Characterization of Jurkat cells used in molecular glue screen. (**C**) Reproducibility of the PLF screen in **Figure 4F** across two biological replicates. Pearson correlation was used for analyses. (**D**) Correlation between protein-protein interaction strength, as measured by PLF, and degradation potency or efficacy, for CK1α and Wee1 glues. Spearman correlation was used for analyses (**E**) Correlation between PLF data (y-axis) and HiBiT degradation data (x-axis) for measuring the specificity of 14 CK1α-targeting molecular glues. All glues with HiBiT data on both CK1α and Wee1 binding were included in this analysis. Non-degraded targets in the HiBiT assay were assigned a $DC_{50}$ of 1 μM. Spearman correlation was used for analyses. (**F**) ZNL-02-096, a PROTAC targeting Wee1, also engages IKZF1 as an off-target. A dose-dependent hook effect is observed for Wee1, while IKZF1 engagement increases monotonically. $N = 2$. Mean ± SD. p (two-tailed Student's t tests) (**G**) CK1α is an endogenous substrate of CRBN that binds weakly even without molecular glues. CC-92480, a potent CRBN binder does not directly engage CK1α or Wee1 but uncovers CRBN–neo-substrate interactions by competing with endogenous CK1α binding. $N = 3$. Mean ± SD. p (two-tailed Student's t tests) (**H**) Adding an amide bond to CK1α molecular glue QXG-0632 enhances the CRBN–CK1α interaction as well as drug efficacy. (**I**) The lenalidomide scaffold exhibits stronger CRBN binding than the thalidomide scaffold, resulting in enhanced potency and efficacy for CK1α degradation. (**J**) SDS-PAGE analysis of ERM-FlexID and mito-FlexID samples used for PLF in **Figure 4I**. BCN-labeled proteins were detected via Click with mTz-BODIPY following cell lysis. Equal protein loading was confirmed by Coomassie staining. (**K**) BCN dose–response for calnexin labeling with ERM-

FlexID, detected by PLF. Beads coated with anti-calnexin antibody were used to capture BCN/mTz-BODIPY–labeled endogenous proteins. $N = 2$. Mean ± SD.

**Supplementary Data and Code Availability**

**Supplementary data 1.** 2D Flow cytometry data for all LplA variants tested. Data are shown for (**A**) LplA orthologs, (**B**) ESM point mutants, (**C**) ProteinMPNN point mutants, (**D**) ESM combined mutants (E3-21), Directed evolution variant (D2), MPNN active-site redesigns (M1-24) and final combined engineering variants (F1-30). The black line in each plot shows the shape of the template (W37V LplA).

**Supplementary data 2.** SEC-SAXS Elution profiles and Oligomer fits. PDFs for each LplA variant tested, showing elution profile and associated SAXS scattering curves with Oligomer-predicted model fits. Data are plotted in both Log(I) vs q and Kratky plot format to better emphasize relevant differences, as well as the residual (data – model) for each fit.

**Supplementary data 3.** Structures used for SEC-SAXS Oligomer analysis. PDB files for all LplA structural models used for Oligomer analysis of the SAXS data.

**Supplementary data 4.** Code used to analyze SEC-SAXS data.

These files can be downloaded at: https://doi.org/10.5281/zenodo.17583062

**Extended Data Table 1. LplA mutant sequences and activities (related to Figure 1).**

- Tab 1: Full LplA sequences for all variants tested. Flow cytometry quantification of promiscuous activity and expression level.
- Tab 2: All LplA point mutations scored by ESM1b/1v set of 6 models, and proteinMPNN pseudo-log-likelihoods for all single point mutants
- Tab 3: Column definitions.

**Extended Data Table 2. FlexID labeling site mapping (related to Figure 2M).**

- Tab 1: FlexID-labeled sites identified by superTOP-ABPP[5]. Peptides identified from all three biological replicates were consolidated for pLogo analysis.
- Tab 2: MS-identified BCN-labeled peptides from Replicate 1.
- Tab 3: MS-identified BCN-labeled peptides from Replicate 2.
- Tab 4: MS-identified BCN-labeled peptides from Replicate 3.
- Tab 5: Column definitions.

**Extended Data Table 3. True positive and false positive lists used to filter and analyze proteomic data (related to Figure 3 and Extended Data Figure 7).**

- Tab 1: ERM Gold TP: True positive list of 90 well-established ERM proteins[6]. Used to determine ROC cut-offs and to calculate ERM proteome sensitivity.
- Tab 2: ERM TP: Larger true positive list of 14,198 human proteins with secretory annotation in GOCC, Human Protein Atlas, Phobius, Human Plasma Proteome Database, or literature. Used to calculate ERM proteome specificity.

- Tab 3: Non-secretory FP: False positive list of 6,465 proteins with non-secretory annotation in GOCC, Human Protein Atlas, Human Plasma Proteome Database, or literature. This set was generated by excluding the 14,198 secretory proteins in Tab 2 from the 20,663 proteins in the reference human proteome (Tab 5), with the remaining proteins designated as the non-secretory subset. Used to determine ERM/NES ROC cut-offs.
- Tab 4: Mito matrix TP: 162 soluble mitochondrial matrix proteins[7]. Used to determine ROC cut-offs and to calculate 1min Mito matrix proteome sensitivity.
- Tab 5: Non-mitochondrial FP: Proteins with non-mitochondrial annotation. Previously curated false positive list of 2185 human proteins that are not annotated to be mitochondrial. Used to determine 1min Mito/NES ROC cut-offs.
- Tab 6: Mitochondrial TP: mitochondrial annotated proteins. A list of 3817 human proteins present in MitoCarta3.0[8] or annotated with the following Gene Ontology term: GO:0005739, but excluding any proteins also present in Tab 5. Used to calculate 1min Mito proteome specificity.
- Tab 7: Reference human proteome (Uniprot: UP000005640, updated September 1, 2025).
- Tab 8: ERM TP for ERM_UT: True positive list of 90 well-established ERM proteins[6] and 25 additional true ERM proteins that are identified with all three PL enzymes with a FDR <0.01. Used to analyze ROC of ERM/untransduced enrichment.
- Tab 9: Column definitions.

**Extended Data Table 4. FlexID ERM proteome (related to Figure 3).**

- Tab 1: FlexID 5-minute ERM proteome using ROC-based cutoff (790 proteins)
- Tab 2: APEX 1-minute ERM proteome using ROC-based cutoff (989 proteins)

- Tab 3: TurboID 5-minute ERM proteome using ROC-based cutoff (1038 proteins)
- Tab 4: FlexID 5-minute ERM proteome using 10% FPR cutoff (1035 proteins)
- Tab 5: APEX 1-minute ERM proteome using 10% FPR cutoff (953 proteins)
- Tab 6: TurboID 5-minute ERM proteome using 10% FPR cutoff (951 proteins)
- Tab 7: Unfiltered mass spectrometry data from 5-minute FlexID labeling
- Tab 8: Unfiltered mass spectrometry data from 1-minute APEX labeling
- Tab 9: Unfiltered mass spectrometry data from 5-minute TurboID labeling
- Tab 10. Unfiltered mass spectrometry data from 5min PL labeling. Used to analyze ROC of ERM/untransduced enrichment. Missing values from the untransduced samples were imputed with MinDet (See Methods)
- Tab 11. Unfiltered mass spectrometry data from 5min TurboID labeling in the absence or presence of Biotin. Used to analyze ROC of TurboID ERM/NES enrichment for both +/- Biotin conditions. Note that the raw intensity values in Tables 10–11 differ from those in Tables 7–9 because they were generated from independent DIA-NN runs (see Methods).
- Tab 12. Column definitions

**Extended Data Table 5. PLF molecular glue screening data (related to Figure 4).**

- Tab 1: Normalized PLF values were determined for each compound targeting IKZF1, Wee1, and CK1α
- Tab 2: Raw median fluorescence intensity of IKZF1, Wee1, CK1α-immobilized beads under each treatment condition
- Tab 3: Efficacy and potency data for compounds obtained from the HiBiT assay
- Tab 4. Column definitions

**Extended Data Table 6. SEC-SAXS data (related to Figure 2).**

- Tab 1: Averaged SEC-SAXS data across all replicates per condition
- Tab 2: SEC-SAXS complete data with individual replicates
- Tab3: Column definitions

**Table of genetic constructs used in this study**

| Name | Promoter | Details | Used for |
|---|---|---|---|
| His-FlexID-FLAG-mCherry | CMV | Enzymes: FlexID-mCherry fusion<br>His: HHHHHH<br>FLAG: DYKDDDDK | Expression of FlexID–mCherry in HEK293T |
| His-FLAG-FlexID-GS-mCherry | CMV | Enzymes: FlexID, mCherry<br>GS Linker: GSSGSSGSSGSS<br>His: HHHHHH<br>FLAG: DYKDDDDK | Expression of FlexID–mCherry in HEK293T |
| His-FLAG-FlexID-XTEN-mCherry | CMV | Enzymes: FlexID, mCherry<br>XTEN linker: SGSETPGTSESATPES<br>His: HHHHHH<br>FLAG: DYKDDDDK | Expression of FlexID–mCherry in HEK293T |
| His-FLAG-FlexID-EV-mCherry | CMV | Enzymes: FlexID, mCherry<br>EV Linker:<br>SAGGSAGGSAGGSAGGSAGG<br>His: HHHHHH<br>FLAG: DYKDDDDK | Expression of FlexID–mCherry in HEK293T |
| pTRE3G-OMM-V5-FlexID | TRE3G | Target: outer mitochondrial membrane<br>OMM: AKAP1 mito transmembrane domain, MAIQLRSLFPLALPG LLALLGWWWFFSRKK<br>V5: GKPIPNPLLGLDST | Transient and stable expression of FlexID targeted to the OMM in HEK293T |

| | | | |
|---|---|---|---|
| pTRE3G-NLS-V5-FlexID | TRE3G | Target: nucleus<br>NLS: 3x SV40 NLS, SRADPKKKRKVDPKKKRKVDPKKKRKV<br>V5: GKPIPNPLLGLDST | Transient and stable expression of FlexID targeted to the nucleus in HEK293T |
| pTRE3G-mito-V5-FlexID | TRE3G | Target: mitochondrial matrix<br>Mito: human COX4 MTS 1-24aa, MLATRVFSLVGKRAISTSVCVRAH<br>V5: GKPIPNPLLGLDST | Transient and stable expression of FlexID targeted to the mito matrix in HEK293T |
| pTRE3G-V5-ERL-FlexID | TRE3G | Target: ER lumen<br>ERL: Human Sec61b 2-96aa, PGPTPSGTNVG SSGRSPSKAVAARAAGSTVRQRKNASCGTRSAGRTTSAGTGGMWRFYTEDSPGLKVGPVPVLVMSLLFIASVFMLHIWGKYTRS<br>V5: GKPIPNPLLGLDST | Transient and stable expression of FlexID targeted to the ER lumen in HEK293T |
| pTRE3G-ERM-linker-V5-Enzyme | TRE3G | Target: ER membrane<br>ERM: C1 P450 1-27aa, MDPVVVLGLCLSCLLLLSLWKQSYGGG | Transient and stable expression of FlexID targeted to the ER membrane in HEK293T |

| | | | |
|---|---|---|---|
| | | Linker:<br>GSGSGGRGSGSGSGSGSGSGSGG<br>ASGS<br>V5: GKPIPNPLLGLDST<br>Enzyme: FlexID, TurboID, or APEX2 | |
| pTRE3G-V5-NES-Enzyme | TRE3G | Target: Cytosol<br>V5: GKPIPNPLLGLDST<br>NES: LQLPPLERLTLD<br>Enzyme: FlexID, TurboID, or APEX2 | Transient and stable expression of FlexID targeted to the cytosol in HEK293T |
| pYFJ16 6xHis-TEVcs-FlexID | T5 | IPTG induced LplA expression | *E. Coli* expression and purification of FlexID |
| pYFJ16 6xHis-TEVcs-LplA (W37V) | T5 | IPTG induced LplA expression | *E. Coli* expression and purification of LplA(W37V) |
| pYFJ16 6xHis-TEVcs-E1 (LplA | T5 | IPTG induced LplA expression | *E. Coli* expression and purification of E1 (LplA W37V/A48N) |

| | | | |
|---|---|---|---|
| W37V/A48N<br>) | | | |
| pYFJ16<br>6xHis-<br>TEVcs-E2<br>(LplA<br>W37V/A48N/<br>A248S) | T5 | IPTG induced LplA expression | *E. Coli* expression and purification of E2 (LplA W37V/A48N/A248S) |
| pYFJ16<br>6xHis-<br>TEVcs-LplA<br>(WT) | T5 | IPTG induced LplA expression | *E. Coli* expression and purification of LplA(WT) |
| pETDuet-T7-<br>6xHis-<br>TEVcs-<br>AviTag-<br>XTEN-<br>FlexID T7-<br>BirA | T7 | IPTG induced co-expression of FlexID and BirA,<br>BirA and LplA are under two different T7 protomers.<br>AviTag-XTEN-FlexID is biotinylated in situ. | *E. Coli* expression and purification of Biotinylated FlexID |
| pETDuet-T7-<br>6xHis-<br>TEVcs-<br>AviTag- | T7 | IPTG induced co-expression of LplA (W37V) and BirA,<br>BirA and LplA are under two different T7 protomers. | *E. Coli* expression and purification of Biotinylated LplA (W37V) |

| | | | |
|---|---|---|---|
| XTEN-LplA(W37V) T7-BirA | | AviTag-XTEN-FlexID is biotinylated in situ. | |
| pAAV NRXN3b(1-52)-HA-NRXN3b (538-591)-GSlinker-V5-NES-FlexID | Synapsin | Target: Neuron membrane<br>NRXN3b (1-52) and (538-591)<br>HA: YPYDVPDYA<br>GS linker:<br>GSGSTSGSGSGGSGGSGGSSGG<br>V5: GKPIPNPLLGLDST<br>NES: LQLPPLERLTLD | Figure 3F and 3G for AAV-induced expression in neurons and mouse brain |
| FRB-Flag-FlexID | CMV | Bait: FRB domain<br>Enzyme: FlexID<br>FLAG: DYKDDDDK | Figure 4B, 4C, and Extended Data Figure 8A. Transient expression of FRB-fused FlexID in HEK293T |
| V5-FKBP-APEX | CMV | Prey: FKBP-APEX<br>V5: GKPIPNPLLGLDST | Figure 4B, 4C, and Extended Data Figure 8A. Transient expression of V5-FKBP-APEX2 in HEK293T |
| pTRE3G-CRBN-3xFlag-FlexID | TRE3G | Bait: CRBN<br>Enzyme: FlexID<br>3xFLAG:<br>DYKDHDGDYKDHDIDYKDDDDK | Figure 4D and 4F. Stable expression of CRBN-FlexID fusion protein to in the CRBN$^{-/-}$ Jurkat cells |

**Complete amino acid sequence of FlexID**

MSTLRLLISDSHDPWFNLAVEECIFRQMPATQRVLFLVRNADTVVIGRNQNPWKECNIR
RMEEDNVRLARRSSGGGAVFHDLGNTCFTFMAGKPEYDKTISTSIVLNALNALGVSAEA
SGRNDLVVKTVEGDRKVSGSAYRETKDRGLHHGTLLLNADLSRLANYLNPDKKKLAA
KGITSVRSRVTNLTELLPGITHEQVCEAITEAFFAHYGERVEAEIISPNETPDLPNFAETFA
RQSSWEWNFGQSPAFSHLLDERFTWGGVELRFDVEKGHITRAQVFTDSLNPAPLEALAG
RLQGCLYRADELQQECEALLVDFPEQEKELRELSAWMAGAVR. Mutations relative to wild-type *E. coli* LplA (in red): Y11H, W37V, T57I, A48N, F147L, K224E, A248S, H267R, M306E.

**Complete nucleotide sequence of FlexID** (mammalian codon-optimized)

ATGAGCACCCTGAGACTGCTGATCAGCGACAGCCACGACCCCTGGTTCAACCTGGC
CGTGGAGGAGTGCATCTTCAGACAGATGCCCGCCACCCAGAGAGTGCTGTTCCTGGT
GAGAAACGCCGACACCGTGGTGATCGGCAGAAACCAGAACCCCTGGAAGGAGTGC
AACATCAGAAGAATGGAGGAGGACAACGTGAGACTGGCCAGAAGAAGCAGCGGCG
GCGGCGCCGTGTTCCACGACCTGGGCAACACCTGCTTCACCTTCATGGCCGGCAAGC
CCGAGTACGACAAGACCATCAGCACCAGCATCGTGCTGAACGCCCTGAACGCCCTG
GGCGTGAGCGCCGAGGCCAGCGGCAGAAACGACCTGGTGGTGAAGACCGTGGAGG
GCGACAGAAAGGTGAGCGGCAGCGCCTACAGAGAGACCAAGGACAGAGGCCTGCA
CCACGGCACCCTGCTGCTGAACGCCGACCTGAGCAGACTGGCCAACTACCTGAACC
CCGACAAGAAGAAGCTGGCCGCCAAGGGCATCACCAGCGTGAGAAGCAGAGTGAC
CAACCTGACCGAGCTGCTGCCCGGCATCACCCACGAGCAGGTGTGCGAGGCCATCA

CCGAGGCCTTCTTCGCCCACTACGGCGAGAGAGTGGAGGCCGAGATCATCAGCCCCAACGAGACCCCCGACCTGCCCAACTTCGCCGAGACCTTCGCCAGACAGAGCAGCTGGGAGTGGAACTTCGGCCAGTCCCCCGCCTTCAGCCACCTGCTGGACGAGAGATTCACCTGGGGCGGCGTGGAGCTGAGATTCGACGTGGAGAAGGGCCACATCACCAGAGCCCAGGTGTTCACCGACAGCCTGAACCCCGCCCCCCTGGAGGCCCTGGCCGGCAGACTGCAGGGCTGCCTGTACAGAGCCGACGAGCTGCAGCAGGAGTGCGAGGCCCTGCTGGTGGACTTCCCCGAGCAGGAGAAGGAGCTGAGAGAGCTGAGCGCCTGGATGGCCGGCGCCGTGAGATAA

**Table of antibodies used in this study**

| Antibody | Source | Vendor | Catalog Number | Dilution(s) |
|---|---|---|---|---|
| Anti-V5- Alexa Fluor™ 647 | Mouse | Invitrogen | 451098 | WB: 1:2500 |
| Anti-V5 | Mouse | Invitrogen | R960CUS | PLF: 1 μg for 1 μL beads, IF: 1:1000 |
| Anti-V5 | Rabbit | Cell Signaling Technology | 13202S | IF: 1:1000 |
| Anti-Calnexin | Rabbit | Invitrogen | PA534754 | PLF: 1 μg for 1 μL beads, IF: 1:1000 |
| Anti-MTCO1 | Rabbit | Abcam | ab203912 | PLF: 1 μg for 1 μL beads |

| | | | | |
|---|---|---|---|---|
| Anti-GRSF1 | Rabbit | Abcam | ab205531 | PLF: 1 μg for 1 μL beads |
| Anti-Tom20 | Mouse | Santa Cruz Biotechnology | sc-17764 | IF: 1:1000 |
| Anti- Lipoic Acid | Rabbit | MilliporeSigma | 437695 | WB: 1:5000 |
| Anti-β-Actin | Mouse | Cell Signaling Technology | 3700S | WB: 1:5000 |
| Anti-CRBN | Rabbit | Cell Signaling Technology | 71810S | WB: 1:1000 |
| Anti-Wee1 | Rabbit | Cell Signaling Technology | 13084S | PLF: 1 μg for 1 μL beads |
| Anti- CSNK1A1 | Rabbit | Proteintech | 55192-1-AP | PLF: 1 μg for 1 μL beads |
| Anti- IKAROS | Rabbit | Invitrogen | PA5-85570 | PLF: 1 μg for 1 μL beads |
| Anti-mouse-AlexaFluor568 | Goat | Invitrogen | A-21124 | IF: 1:2000 |
| Anti-mouse-AlexaFluor647 | Goat | Invitrogen | A-21241 | IF: 1:2000 |
| Anti-rabbit-AlexaFluor568 | Goat | Invitrogen | A-11036 | IF: 1:2000 |

| | | | | |
|---|---|---|---|---|
| Anti-rabbit-AlexaFluor647 | Goat | Invitrogen | A-21245 | IF: 1:2000 |
| Anti-mouse-IRDye 680RD | Goat | LI-COR Biosciences | 92668070 | WB: 1:10000 |
| Anti-rabbit-IRDye 800CW | Goat | LI-COR Biosciences | 92632211 | WB: 1:10000 |
| IRDye® 800CW Streptavidin | N/A | LI-COR Biosciences | 92632230 | WB: 1:10000 |

**Methods**

### Methods

#### Cloning and mutagenesis

Genes and gene fragments were PCR-amplified using Q5 polymerase (NEB) and vectors were digested using restriction enzymes. All fragments were gel-purified and ligated using Gibson assembly (NEB), then transformed into stable competent *E. coli* (NEB) or XL1-Blue competent *E. coli* (Agilent). The FlexID gene was codon-optimized for mammalian expression[9]. Plasmids were checked by whole plasmid sequencing. See **Table of Genetic Constructs Used in this Study** for further details.

Point mutants were cloned by Gibson assembly into the W37V LplA mammalian expression vector. PCR primers were designed to enable amplification of the LplA plasmid in two halves, with one junction falling within the LplA gene, such that this junction could be used to introduce point mutations. Mutagenic primers were designed using primerX (https://www.bioinformatics.org/primerx/ ). Cloned variants were confirmed by whole plasmid sequencing.

LplA variants combining multiple mutants were cloned by up to 6-way Gibson assembly of fragments containing the desired mutations. BamHI and EcoRI-digested vector was combined with PCR-amplified fragments using Gibson assembly.

#### Selection of LplA orthologs and generation of bacterial phylogenetic tree

LplA orthologs were selected to sample orthologs with similar alignment scores to *E. coli* LplA, while covering a broad range of bacterial phylogeny. We prioritized selecting some orthologs with prior experimental evidence of catalyzing the complete two-step lipoyl adenylation and lipoyl transfer reaction[10,11]. Structural and sequence alignment enabled identification of the analogous X37 position in all LplA orthologs, which was mutated to valine in each construct to expand the active site to accept BCN as a substrate.

The bacterial phylogenetic tree was downloaded from Genome Taxonomy Database (GTDB) along with metadata containing descriptions of all species listed in the tree. Using a Python script, the tree was pruned from ~136,000 tips to ~15,000 tips by matching tip names (listed in GTDB accession number) to corresponding NCBI organism names, and dropping entries without well characterized NCBI organism names (Genus, Species). Species with multiple strains were deduplicated by selecting one strain to serve as a representative. The tree was then visualized in R using the ape package, and tips were colored to highlight species for which LplAs were selected for testing.

Orthologs were cloned by gene-synthesis of the complete LplA ortholog genes, followed by Gibson assembly into the digested mCherry-LplA expression plasmid backbone, and confirmed by whole-plasmid sequencing.

**ESM design of LplA point mutants**

We followed the method established in Hie et al.[12] for the implementation of ESM1b/1v, wherein we select amino acid substitutions recommended by a combination of 6 language models[12]: the ESM1b collection of 5 models and ESM1v[13]. Code and details are available at: https://github.com/brianhie/efficient-evolution. We take as input the LplA wild-type and W37V sequences in separate runs, and accept all mutations predicted by any of the models to be above the WT / W37V sequences in log-likelihood. In the case of LplA, this was 48 point-mutations.

**ProteinMPNN design of LplA point mutants**

We selected ProteinMPNN[14] point mutations based on those with the highest pseudo-log-likelihood conditioned on the structure backbone coordinates, for each of LplA's conformational states (open, PDB 3A7R; and closed, chain A of PDB 1X2G). We take as input the WT LplA sequence and accept only the ~35 top-scored mutations predicted towards each of the conformations. WT and W37V sequence inputs yielded very similar results[2]. Gain-of-lysine variants were largely excluded to avoid the possibility of increased self-labelling activity, which might lead to inaccurate measurement of the promiscuous activity.

**ProteinMPNN re-design of LplA active site**

ProteinMPNN redesign[14] was carried out using the LplA structure in the open conformation (PDB ID: 3A7R) and sequence design was limited to the first shell surrounding the W37V mutation. Redesigns were made in the context of the E1 or E2 sequences, and the W37 position was allowed to be designed to V, A, I, or L. Structures of resulting sequences were predicted with AlphaFold using multiple sequence alignments generated with HHblits[15]. Predictions were filtered for high confidence (pLDDT > 90) and designs were manually inspected to ensure integrity of the active site.

**Mammalian cell culture**

HEK293T cells (CRL-3216, ATCC) and Lenti-X 293T cells (632180, Takara) were cultured as monolayers in complete growth medium: Dulbecco's Modified Eagle Medium (DMEM, Corning) supplemented with 10% Fetal Bovine Serum (FBS, VWR). Jurkat cells and CRBN-KO Jurkat cells (Gifts from Dr. Woong Sub Byun) were cultured as suspension in RPMI-1640 Medium (11875119, Thermo Scientific) supplemented with 10% Fetal Bovine Serum (FBS, VWR). All cell lines were cultured at 37 °C under 5% $CO_2$. For experimental assays, HEK293T cells were grown in 6-well, 12-well, 24-well, or 96-well plates pretreated with 20 μg/mL human fibronectin (Millipore) for at least 10 min at 37 °C.

**Transient expression of LplA/FlexID in HEK 293T cells**

24 well plates were coated with 200 μL human fibronectin (HFN) for 30 minutes at 37 °C. HFN was aspirated, and then HEK293T cells were plated at 1 x $10^5$ cells per well. Following 12-24h of cell growth to 60-80% confluency, transfections were performed. For each well, 500 ng of the mCherry-FlexID plasmid and 4 μL of PEI transfection reagent were combined in 50 μL of DMEM without FBS or antibiotics, incubated for 20 minutes at room temperature, and then added to each well of a 24-well plate dish. Cells were grown for 24 - 48 hours after transfection to allow for LplA expression, and mCherry signal was confirmed by imaging.

**Flow cytometry assay for LplA promiscuous activity in HEK 293T cells**

Following successful transfection of cells, reagents necessary for labeling were prepared. A 200 mM working stock of the labeling probe Endo-BCN Pentanoic Acid (BCN) (Broadpharm, BP-24361) was prepared by dilution of solid BCN in DMSO. A labeling solution of 200 μM BCN in DMEM+FBS+penicillin/streptomycin (1000x dilution) was prepared from this. Next, a stock solution of 1 mM methyltetrazine-BODIPY (Conjuprobe, CP-4018) was prepared in DMSO, then diluted 5000x in complete growth media. Both labeling and BODIPY solutions were pre-warmed to 37 °C, along with additional complete media for washes. The growth media was removed from the transfected HEK293T cells, after which 0.5 mL of BCN solution was added to each well. The cells were moved to a 37 °C cell culture incubator for 5 minutes, and then labeling media was removed by aspiration and immediately replaced with warmed complete media. Three additional 5-minute washes with warmed complete media were performed to remove excess BCN substrate. Afterwards, the media was aspirated and replaced with 0.5 mL/well of the fluorophore-click solution (200 nM mTz-BODIPY) and cells were incubated at 37 °C for 45 minutes. Media was then removed by aspiration and cells were quickly washed twice in pre-warmed complete media. Three additional 15-minute washes (15 min incubation between media addition and aspiration) in complete media were performed to remove unbound fluorophore. Once media from the final wash was removed, cells were incubated for 1 minute in 100uL of enzyme-free cell dissociation buffer at 37°C, and then lifted with an additional 100uL flow buffer (PBS + 3% FBS) and transferred to a 96-well plate. Cells were pelleted once by spinning at 400g for 5 minutes at 4°C, and then resuspended in flow buffer before analyzing on a BioRad ZE5 flow cytometer. BFP was detected using a 405 nm laser, with a 460/22 emission filter. mTz-BODIPY was detected using a 488 nm laser, with a 509/24nm emission filter. mCherry was detected using a 561 nm laser, with 615/24 nm emission filter. BFP was detected using a 405 nm laser, with a 460/22 nm emission. Forward and side scatter were detected using the 488 laser with a 488/10 emission filter. Each sample was agitated for 5 s, and run with a flow rate of 1 µl/s.

### Quantitation of flow cytometry data

The following gates were used for the flow cytometry assay to assess LplA promiscuous activity in HEK293T cells:

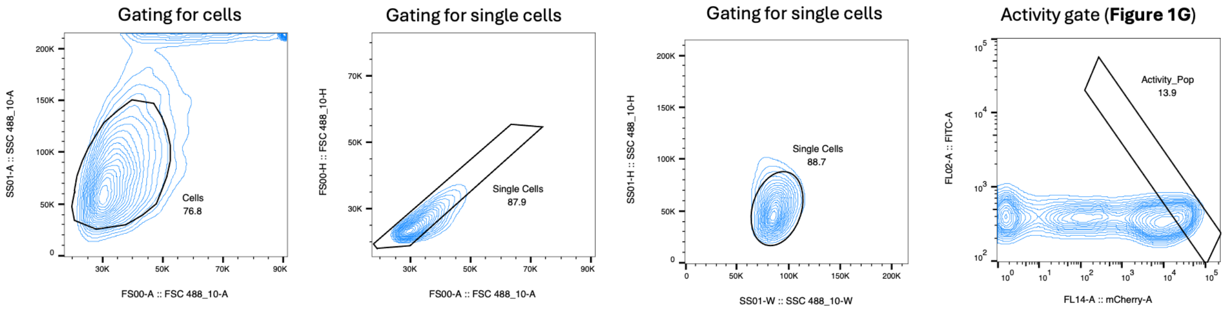


Single cells were gated by forward scatter/side scatter profiles as above. After gating, cells were plotted by mCherry against FITC channels, and a diagonal gate was used to capture the linear region for the mCherry:BODIPY curve. Cells within this gate were analyzed for median BODIPY/mCherry signal to quantify the promiscuous labelling activity of each variant. All data was analyzed using the software package FlowJo.

### SDS-PAGE analysis of LplA promiscuous activity

HEK293T cells transiently or stably expressing FlexID or its variants under a pCMV or a pTRE3G promoter were incubated or induced with 100–300 ng/mL doxycycline for 24-48 hours. After validation of expression of transfected constructs, the cell growth media was removed and labeling was carried out by addition of pre-warmed labeling media (200 µM BCN in complete DMEM) for 5 minutes at 37C. Labelling was quenched with 3x washes with ice-cold PBS.

To lyse cells for protein enrichment, RIPA buffer was supplemented with 1:100 dilutions of protease inhibitor cocktail (PIC) and phenylmethylsulfonyl fluoride (PMSF). Labeled cells were resuspended by pipetting in RIPA buffer, and cell suspensions were incubated on ice for 10 minutes at 4°C. Following lysis, cell debris was removed by centrifuging the lysate at 20, 000 × g for 10 minutes at 4°C. The supernatant was collected for subsequent analyses.

To visualize BCN-labeled proteins with in gel fluorescence, lysates were further labelled via the inverse-electron demand Diels-Alder Cycloaddition (IEDDAC) reaction of BCN with methyl-tetrazine (mTz, mTz-BODIPY, or mTz-JF647). mTz-Dye was added to the clarified lysate at a final concentration of 4 µM and allowed to react for 2 hours at 4°C with vortexing at 850 rpm. For samples in which the fluorescence of protein tags (mCherry and YFP) needed to be preserved, unreacted methyl-tetrazine dye was removed by incubating the lysates with TCO-agarose. Specifically, 1 µL of TCO-agarose was added per 50 µL of lysate, followed by incubation for 1 h at 4 °C with continuous agitation (850 rpm). The resin-treated lysates ("clicked and purified") were subsequently combined with protein loading buffer at a 5:1 ratio and mixed thoroughly without boiling. For all other samples, unreacted methyl-tetrazine dye was removed by methanol–chloroform precipitation (4:1, v/v). Precipitated lysates were collected, air-dried, and resuspended in 3× protein loading buffer prior to analysis.

Lysate was loaded per well of gradient SDS-PAGE gels (4-12%), and run in SDS running buffer (Tris-Glycine) for 45 minutes at 180V. Gels were then washed briefly in milliQ water, and then imaged on a Typhoon biomolecular imager. mTz-BODIPY was imaged using a 480nm laser, with a 520nm emission filter. YFP and mCherry were imaged using a 532nm laser, with 610nm emission filter. JF-647 was imaged with a 633nm laser, using a 670nm emission filter.

In gel fluorescence data was analyzed and quantified using FIJI. Labelling and expression were background-subtracted, and the labelling was normalized to expression level for each variant to calculate the expression-normalized activity. Then the expression-normalized activity of each variant is divided by the expression-normalized activity of the template W37V to calculate the relative activity for each variant.

To visualize BCN-labeled proteins with streptavidin blotting, lysates were labelled with 4 μM mTz-Biotin for 2 hours at 4°C with vortexing at 850 rpm. Unreacted mTz-Biotin was removed by methanol–chloroform precipitation (4:1, v/v). Precipitated lysates were collected, air-dried, and resuspended in 3× protein loading buffer prior to analysis. Samples were then loaded on 4-12% SDS–PAGE gels, transferred onto PVDF membranes, and stained with Ponceau S (Sigma) to visualize total protein loading. Blots were blocked in 5% (w/v) BSA in 1× tris-buffered saline with Tween (TBST) (Teknova) for 60 min at room temperature, incubated with mouse anti-V5-AF647 antibody (1:2, 500) against FlexID's epitope tag and streptavidin-IRDye800 (1:10, 000) in TBST for 1 h at room temperature, washed three times with TBST for 5 min each, and imaged on an Odyssey CLx imager (LICOR). See antibody sources and dilutions used in the Supplementary Information.

**Fluorescence microscopy screening for FlexID Engineering**

48 well glass-bottom plates were coated with 100uL human fibronectin (HFN) for 30 minutes at 37 °C. HFN was aspirated, and then HEK293T cells were plated at 4.5E4 cells per well. Following 12-24h of cell growth to 60-80% confluency, transfections were performed. For each well, 250ng of the mCherry-LplA plasmid and 2uL of PEI transfection reagent were combined in 25uL of DMEM without FBS or antibiotics, incubated for 20 minutes at room temperature, and then added

to each well of the 48-well plate dish. Cells were grown for 24 - 48 hours after transfection to allow for LplA expression.

Labelling was carried out as in the flow-cytometry assay. The cell growth media was removed and labeling was carried out by addition of pre-warmed labeling media (200 µM BCN in complete DMEM) for 5 minutes at 37C, and then immediately replaced with warmed complete media. Three additional 5-minute washes with warmed complete media were performed to remove excess BCN substrate. Afterwards, the media was aspirated and replaced with 0.5mL/well of the fluorophore-click solution (200nM mTz-BODIPY) and cells were incubated at 37°C for 45 minutes. Media was then removed by aspiration and cells were quickly washed twice in pre-warmed complete media. Three additional 15-minute washes (15 min incubation between media addition and aspiration) in complete media were performed to remove unbound fluorophore. Once media from the final wash was removed, cells were washed 3x in PBS, and treated with 4% paraformaldehyde for 15 minutes at room temperature. Cells were again washed 3x in PBS, and imaged using an Olympus APX100 microscope with a 20x objective. mTz-BODIPY was detected using a 480 nm laser, with a 520nm emission filter, and mCherry was detected using a 532 nm laser, with 610 nm emission filter. Images were analyzed using FIJI.

The following combinations of laser excitation and emission filters were used for various fluorophores: Bodipy (491 laser excitation; 528/38 emission), mCherry (561 laser excitation; 617/73 emission). Acquisition times ranged from 100 to 500 *ms*. All images were collected using SlideBook (Intelligent Imaging Innovations) and processed using FIJI/ImageJ.

**PL enzyme imaging comparison**

HEK 293T cells stably expressing FlexID, TurboID, and APEX2 targeted to ER membrane were plated onto glass bottom 24-well plates at 60% confluency and induced with 300 ng/mL dox for 20 hours. TurboID expressing cells were then labeled for 10 minutes at 37 °C with 200 μM biotin and quenched with 3x cold DPBS wash. APEX2 expressing cells were pre-incubated for 30 minutes with 500 μM biotin-phenol and labeling was initiated by the addition of 1mM $H_2O_2$ for 1 minute. APEX2 labeling was quenched with a 3x wash with a solution of 10 mM ascorbate, 5mM Trolox, 10 mM sodium azide in DPBS, followed by 3x cold DPBS wash. FlexID expressing cells were labeled for 10 minutes at 37 °C with 200 μM BCN and quenched with 3x cold DPBS wash.

All cells were subsequently fixed and permeabilized for 10 minutes at 4 °C with methanol pre-chilled at -20C. Following fixation, FlexID labeled cells were clicked with 4 μM mTz-biotin for 1 hour at room temperature. All cells were then blocked overnight at 4 °C with 1% BSA in DPBS. Primary staining was performed for 1 hour at room temperature with 1:1000 mouse-anti-V5 and 1:1000 rabbit anti-calnexin in 1% BSA in DPBS. Following 3x5 minute washes in DPBS, secondary staining was performed with 1:2000 mouse-AF568, 1:2000 rabbit-AF647, and 1:2000 NaV-AF488 for 1 hour at room temperature in 1% BSA in DPBS. Cells were imaged by confocal fluorescence microscopy (Zeiss). Images were collected with Slidebook (Intelligent Imaging Innovations).

To quantify colocalization between PL enzyme expression and labeling signals, three fields of view showing labeling, expression, and calnexin staining were acquired for each ERM-PL enzyme across three biological replicates. TIFF images were processed with Cellpose[16] to segment expressing cells into individual ROIs using a radius of 220. These ROIs were applied to all corresponding TIFF channels, and the Pearson correlation coefficient between expression and labeling signals was calculated using the JACoP plugin. Statistical analyses were performed using Kruskal-Wallis test with Dunn's nonparametric multiple comparison test (****$p < 0.0001$).

**Immunofluorescence detection of FlexID activity in different subcellular compartments**

HEK 293T cells stably expressing FlexID targeted to the outer mitochondrial membrane (OMM), mitochondrial matrix (mito), ER membrane (ERM), ER lumen (ERL), cytosol, and nucleus were plated at 70% confluency and induced with 300 ng/mL of dox. After 20 hours of expression, all cells except ERL-FlexID cells were labeled for 10 minutes at 37 °C with 200 μM BCN. ERL-FlexID cells were labeled for 30 minutes at 37 °C with 200 μM BCN. After labeling, cells were clicked with 200 nM mTz-BODIPY for 40 minutes at 37 °C, then washed in DMEM + 10% FBS for 1 hour. Cells were then fixed with 4% PFA for 15 minutes at room temperature, washed 3x with DPBS, and permeabilized with pre-chilled methanol at -20 °C for 5 minutes. Following a 3x DPBS wash, cells were blocked for 1 hour at room temperature with 1% BSA in DPBS. OMM and mito cells were stained with 1:1000 mouse anti-TOMM20 and 1:1000 rabbit anti-V5 for 1 hour at room temperature with 1% BSA in DPBS. ERM and ERL cells were stained with 1:1000 rabbit anti-calnexin and 1:1000 mouse anti-V5 for 1 hour at room temperature with 1% BSA in DPBS. Secondary staining was performed with 1:2000 rabbit-AF647 and mouse-AF568 for OMM and mito cells and 1:2000 mouse-AF647 and rabbit-AF568 for ERM and ERL cells. All cells were stained for 5 minutes with DAPI, washed with DPBS, and imaged by confocal fluorescence microscopy.

**Immunofluorescence detection of C-terminal tagged FlexID activity**

HEK293T cells were plated onto 24-well glass-bottom plates at 60% confluency and transfected with 500 ng of DNA encoding each LplA–mCherry variant. After 20 hours of expression, cells expressing C-terminal LplA–mCherry fusions were labeled with 200 μM BCN for 20 minutes at 37 °C. Cells expressing N-terminal mCherry–LplA fusions were labeled with 200 μM BCN for 10 minutes at 37 °C. All cells were then reacted with 200 nM mTz-BODIPY for 40 minutes at 37 °C, followed by a washout in DMEM + 10% FBS for 1 hour. Nuclei were stained with Hoechst for 5 minutes, cells were washed three times in HBSS, and live-cell imaging was performed by confocal fluorescence microscopy.

**Synthesis of BCN-AMS and BCN-AMP**

Unless otherwise noted, all reagents were purchased from commercial suppliers and used without further purification. Reactions were monitored using a Waters Acquity UPLC/MS system (Waters PDA eλ Detector, QDa Detector, Sample manager─FL, Binary Solvent Manager) using Acquity UPLC BEH C18 column (2.1 × 50 mm, 1.7 μm particle size): solvent gradient = 85% A at 0 min, 1% A at 1.7 min; solvent A = 0.1% formic acid in water; solvent B = 0.1% formic acid in Acetonitrile; flow rate: 0.6 mL/min. Analytical thin layer chromatography (TLC) was performed on Merck silica gel 60 F254 TLC glass plates and analytes were visualized by fluorescence quenching (using 254 nm light). Purification of reaction products was carried out by preparative RP-HPLC using Waters SunFireTM Prep C18 column (19 × 100 mm, 5 μm particle size) with a gradient of 10–90% methanol in water over 40 min (45 min run time) at a flow of 40 mL/min.

<u>5'-Sulfamoyladenosine</u> (**3**)

20 mg of 2′,3′-isopropylidene sulfamoyladenosine (Chempartner, 0.05 mmol) was dissolved in trifluoroacetic acid (TFA, 0.7 mL) and water (0.3 mL) and stirred at room temperature for 1 hour. TFA was then removed under reduced pressure, and the residue was neutralized to pH 7 using $NaHCO_3$. The resulting aqueous solution was lyophilized directly without further purification.

2,5-dioxopyrrolidin-1-yl 5-(((bicyclo[6.1.0]non-4-yn-9-ylmethoxy)carbonyl)amino)pentanoate (**4**)

1-Ethyl-3-(3-dimethylaminopropyl)carbodiimide hydrochloride (EDCI·HCl, 71 mg, 0.37 mmol) was added to a solution of endo-BCN-pentanoic acid (100 mg, 0.34 mmol) and N-hydroxysuccinimide (43 mg, 0.37 mmol) in dichloromethane (DCM, 1.0 mL). The reaction mixture was stirred at 20 °C for 18 h. After completion, the mixture was diluted with ethyl acetate and washed successively with water, saturated aqueous $NaHCO_3$, and brine. The organic layer was dried over anhydrous $Na_2SO_4$, filtered, and concentrated under reduced pressure to afford the crude product **4** (54 mg, 37.6%), which was used in the next step without further purification.

BCN-AMS (**1**)

**4** (25 mg, 0.0641 mmol) was dissolved in dry DMF (1 mL), and 1,8-diazabicyclo[5.4.0]undec-7-ene (DBU, 22 μL, 0.15 mmol) was added along with **3** from the entire reaction. The reaction mixture was stirred at 4 °C, then gradually warmed to room temperature and stirred overnight. The reaction mixture was concentrated under reduced pressure, and the residue was purified by HPLC. The collected fractions were lyophilized to afford the desired product as a white solid DBU salt (6.71 mg, 17.3%).

BCN-AMP

1-Ethyl-3-(3-dimethylaminopropyl)carbodiimide hydrochloride (EDCI·HCl, 1.3 g, 6.8 mmol) was added to a solution of endo-BCN-pentanoic acid (100 mg, 0.34 mmol) and adenosine-5′-monophosphate disodium salt (120 mg, 0.31 mmol) in water (0.75 mL) and pyridine (0.75 mL). The reaction mixture was stirred at 4 °C overnight. The mixture was diluted with water, washed twice with ethyl acetate, and purified by HPLC. The collected fractions were lyophilized to yield the desired product as a white solid (7.46 mg, 3.9%).

**Expression and purification of LplA from *E. coli***

Chemically competent BL21 (DE3) pLysS *E. coli* cells (69451-4, EMD Millipore) were thawed on ice and then transformed with plasmids encoding LplA variants following the producer's recommended transformation protocol. Transformed cells were plated onto LB carbenicillin agar plates and incubated overnight at 37 °C. After incubation and observation of colonies, 5–10 colonies were picked and inoculated into 5 mL of LB-carbenicillin medium and then shaken at 220 RPM at 37 °C for over 6 hours until the culture became visibly cloudy. The 5 mL overnight culture was added to 500 mL of LB- carbenicillin in a 2-liter flask. The culture was shaken at 37 °C until an A600 of 0.5 was reached, approximately 2–3 hours depending on the starting culture. The flask was then moved to a room-temperature shaker for cooling, and isopropyl-β-D-thiogalactopyranoside (IPTG) was added to a final concentration of 100 μg/mL. Induction was performed by shaking the flask at 220RPM overnight at room temperature (~8-16 h). In situ biotinlyation of Avi-tagged LplA were triggered by adding 50 μM of Biotin after induction.

Cells were pelleted by centrifugation at 5,000 × g for 10 min at 4 °C. The supernatant was discarded, and the bacterial pellet was kept on ice. For each sample, 100 μL of 100 mM PMSF and 100 μL protease inhibitor cocktail were added to 20 mL Bacterial Protein Extraction Reagent (B-PER). The pellet was lysed by thoroughly resuspending in 10 mL of the prepared B-PER. After thorough resuspension, an additional 10 mL of the prepared B-PER was then added to ensure thorough suspension. The suspension was gently agitated for 10 min at 4 °C. The lysate was transferred to high-speed centrifuge tubes and centrifuged at maximum speed for 60 min at 4 °C. The resulting supernatant, containing the cell lysate, was collected in a 50 mL conical tube. Nickel-NTA resin for purification was prepared by loading approximately 1 mL of packed resin into a Poly-Prep column, followed by washing with five column volumes of ice-cold nickel-binding buffer (50 mM Tris, pH = 7.8 and 300 mM NaCl). 20 mL of ice-cold nickel-binding buffer was added to the cell lysate, and the prepared resin was transferred to the lysate tube. The mixture was gently agitated for 20 min at 4 °C to facilitate His-tagged protein binding to the resin. The cell lysate/resin mixture was loaded onto a Poly-Prep column, avoiding the column drying out. The resin was washed with ten column volumes of nickel NTA wash buffer (50 mM Tris, pH = 7.8, 10 mM imidazole and 300 mM NaCl) to remove non-specifically bound proteins. Bound protein was eluted using 3 mL of nickel-NTA elution buffer (50 mM Tris, pH = 7.8, 100 mM imidazole and 300 mM NaCl). DTT and glycerol were added to a working concentration of 2 mM and 5% v/v respectively. His-TEV protease (Berkeley QB3 macro-lab) (1:50 molar ratio) was added to the eluate for cleavage, and the sample was incubated overnight at 4 °C.

The eluate was then diluted 20-fold to achieve DTT and imidazole concentrations of <0.2 mM and <5 mM, respectively. Nickel-NTA resin (~1 mL packed per ~5 mg protein) was prepared and washed, then used to bind the reaction mixture for 20 min with gentle agitation at 4 °C. The suspension was loaded onto the column, and the flow-through, containing tag-less LplA, was collected. The resin was washed with one column volume of nickel-binding buffer, and the flow-through was again collected and combined. DTT and glycerol concentrations were re-adjusted to 2 mM and 5% v/v, respectively. The protein solution was concentrated to <500 μL using a protein concentrator (Amicron Ultra, MW 10 kDa, UFC801024, EMD Millipore), maintaining a concentration below 10 mg/mL of protein to minimize aggregation. The sample was centrifuged at 20,000 × g for 10 min, and the supernatant was loaded onto a preequilibrated SEC column. A280 positive fractions were collected, and again the protein solution was concentrated using a protein concentrator (A micron Ultra, MW 10 kDa, UFC801024, EMD Millipore), maintaining a concentration below 10 mg/mL of protein to minimize aggregation. Protein concentration was then quantified using a DTT-compatible BCA assay (23250, Thermo Scientific), and protein samples were aliquoted, flash-frozen in liquid nitrogen, and stored at –80C until use.

**SEC-SAXS**

SEC-SAXS experiments were conducted at the Stanford Synchrotron Radiation Light Source (SSRL) Bio-SAXS beamline 4-2, BL4-2[17]. SEC-SAXS data were collected at High-Resolution mode using a Superdex 75 Increase 3.2/300 column (Cytiva) using DTT-supplemented SEC running buffer (40 mM Tris pH 7.5, 5mM $MgCl_2$, 150 mM NaCl, 5mM DTT, 0.5mM EDTA).

Protein samples without N-terminal His-tags were prepared using the purification method described above, and concentrated by spin-filtration to 1-10 mg/mL in SEC running buffer. We added lipoyl-AMS (or lipoyl-AMP for controls) to a final concentration of 500 μM at 4 °C, 20 minutes prior to SEC-SAXS data collection. Samples with ATP or ATP and BCN were incubated at 4 °C with 500 μM BCN and 1mM ATP for 20 minutes prior to SEC-SAXS data collection.

500 images were acquired with 2-second exposure every 5 seconds at a flow rate of 0.05 mL/min. To minimize potential sample-cell fouling, the x-ray shutter was closed after the 100th

image (blank data collection) and the sample-cell was immediately washed in preparation for the following fractions of interest. The shutter was then re-opened to resume the image acquisitions at the fractions of interest (as determined during test-runs to establish the expected elution point for LplA and variants). Data collection was conducted using the SEC-SAXS tab of BL4-2 *Blu-Ice*[17], and data reduction and initial analyses were performed using the BL4-2 automated SEC-SAXS data processing and analysis pipeline, *SECPipe* (https://www-ssrl.slac.stanford.edu/smb-saxs/node/1860). It implements the programs *SASTOOL* (https://www-ssrl.slac.stanford.edu/smb-saxs/node/1914) and *ATSAS AUTORG*[18]. The program *GNOM* was used for the indirect Fourier transform to estimate the distance distribution function *P*(r). Evolving factor analysis (EFA) was used for data inspection using the program *BioXTAS RAW*[19]. The data were plotted as I(q) versus q, where $q = 4\pi\sin(\theta)/\lambda$, $2\theta$ is the scattering angle, and $\lambda$ is the wavelength of the X-ray. Images were grouped into 5-image windows and SAXS scattering curves averaged across each window.

We next analyzed all samples using Oligomer to estimate the volume fractions of open and closed conformations[20]. We analyzed all windows of 5 images for which every daughter image had an average $R_g$ greater than 10, in order to select for the elution peak of LplA (as LplA has an expected $R_g$ of 22 at minimum). These scattering curves were trimmed by 6 datapoints from the beginning and cutoff at a q value of 0.4 prior to Oligomer prediction of fractional occupancy. Based on our previously-established analysis method[2] we selected a set of models for Oligomer analysis consisting of (1) a Chai-1 predicted closed monomer, (2) the crystal structure of the open conformation monomer from PDB 3A7R, (3) the crystal structure of the open conformation dimer from PDB 3A7R, (4) an AlphaFold3-predicted closed tetramer, and (5) an AlphaFold3-predicted closed dimer. For each sample, the Oligomer predictions were weighted by the average UV280 for the window of analysis, and then averaged, to generate a weighted-average oligomer predicted set of volume fractions. Samples were then grouped by their identity, and volume fractions 2 and 3 were added to give the total open conformation fractional occupancy for each sample.

*Ab initio* electron density maps of all samples were calculated using the program *DENSS*[21], with a final map calculated by averaging 20 independent runs.

**Differential Scanning Fluorimetry (DSF)**

LplA variants were diluted in DSF buffer (1× PBS, pH 7.5, 5 mM $MgCl_2$) to a final concentration of 5 µM and incubated with a 4-fold serial dilution of BCN-AMS (final concentrations: 200 µM to 0). SYPRO Orange (5×; 1:1000, Invitrogen S6650) was included in all reactions. Reactions (10 µl each) were dispensed in two replicates into a white 384-well microtiter plate (Axygen PCR-284-LC480WNFBC, Lot 09819000). Fluorescence was monitored on a Bio-Rad CFX96 qPCR system using the FRET channel, with measurements recorded every 30 s at 0.5 °C intervals from 12.5 °C to 95 °C. Fluorescence data were normalized to the maximum signal, plotted as normalized fluorescence versus temperature, and fitted with a sigmoidal dose–response (variable slope) equation in GraphPad Prism. Melting temperatures ($T_m$) were defined as the inflection point, corresponding to the temperature at which the signal reached 50% between the curve's bottom and top plateaus.

**Biolayer interferometry (BLI)**

BLI was used to measure BCN-AMS off-rate from immobilized enzymes (**Figure 2F**). We generated site-specifically biotinylated LplAs by expressing Avi-LplA together with *E. coli* BirA in Chemically competent BL21 (DE3) pLysS *E. coli* cells (69451-4, EMD Millipore) following the same protocol as shown in the **Expression and purification of LplA from *E. coli* section**. In situ biotinylation of Avi-tagged LplA were triggered by adding 50 μM of Biotin after induction. Two linker strategies were used to immobilize FlexID on the SA sensors (Avi-XTEN-FlexID or Avi-GGGGS-FlexID). Both linkers performed similarly, and Avi-XTEN-FlexID was selected for the final analysis. Complete biotinylation was confirmed with NeutrAvidin gel shift assay. 1 μM of Avi-FlexID was mixed with 5 μM of NeutrAvidin Protein (31000, Thermo Scientific) in PBS buffer and incubated for 60 min at room temperature. Samples were mixed with loading buffer without heating and directly analyzed by SDS-PAGE gel followed by Coomassie staining.

100 nM biotinylated LplA proteins in loading buffer (PBS pH = 7.4, 0.05% Tween 20, 5 mM $MgCl_2$, 0.1% DMSO) were immobilized to give a relative intensity of 1 nm on streptavidin biosensors. Association (t = 0-60 s) and dissociation (t = 60 -190 s) cycles of compounds were started by dipping sensors into BCN-AMS solutions and control buffer. Binding signals were double reference-subtracted from empty tip control and DMSO control. The resulting data were analyzed based on a 1:1 binding model from which $k_{on}$ and $k_{off}$ values were obtained. LplA undergoes a significant conformational change upon binding to the reaction intermediate, leading to an increase in the BLI signal. This signal serves as a readout for intermediate binding. To ensure that the rate-limiting step is the release of the intermediate rather than the conformational change itself, we monitored BCN-AMP binding and hydrolysis in the presence of E2P. In situ BCN-AMP generation was initiated by dipping FlexID-immobilized sensors into plate wells containing 200 μM BCN and 1 mM ATP. Ligand dissociation and hydrolysis were then initiated by transferring the sensors into wells containing either 5 μM E2P or loading buffer. Binding signals were double reference–subtracted using empty-tip and DMSO controls. BCN-AMP was rapidly consumed by E2P, and the BLI signal quickly returned to baseline, indicating that the rate of intermediate release, rather than the conformational change, is the rate-limiting step.

**Tryptophan fluorescence quenching**

Purified LplA variants were diluted to 5 µM in assay buffer (DPBS supplemented with 1 mM DTT, 5 mM $MgCl_2$, 0.05% Tween-20, and 5% glycerol). 90 µL of each protein solution were dispensed into black 96-well plates. Baseline intrinsic fluorescence was measured at room temperature five times using a Tecan plate reader (excitation/emission = 290/340 nm). Subsequently, 10 µL of 1 mM Lipoyl-AMS or buffer control was added, and fluorescence was monitored every ~10 s for 5 min under identical settings. Relative fluorescence (%) was calculated as post-addition fluorescence divided by baseline fluorescence.

**Generation of stable mammalian cell lines**

Stable cell lines were generated through infection by lentivirus. To produce lentivirus, LentiX cells were cultured in 10-cm dishes and transiently transfected with 10.584 µg lentiviral plasmid, 7.56 µg psPAX2, 3.276 µg pMD2.G packaging plasmids using 64.26 µL Mirus TransIT-LT1 transfection reagent (MIR 2300, Mirus) in 2.52 mL OptiMEM. After 6 hours of transfection, remove all volume from each plate and replace slowly with DMEM supplemented with 1X ViralBoost reagent (VB100, Alstem Cell Advancements). After 18 hours of incubation at 37 °C under 5% $CO_2$, cell supernatant was filtered through 0.45-mm filters, concentrated using lentivirus precipitation solution (VC100, ALSTEM) at 4 °C overnight, and centrifuged at 1,500g for 30 min at 4 °C to collect virus pellets. The virus pellets were resuspended in 1000 µl cold Opti-MEM for transduction of cells. 1 - 30 µL of concentrated virus was added on to a 6-well dish of approximately 50% confluent HEK293T or Jurkat cells at 0.5 x $10^6$ cells per mL. After 2 days, cells were passaged and selected with 1 µg/mL of puromycin for 1 week.

### Comparison of PL enzyme fingerprints by streptavidin blot

HEK 293T cells stably expressing FlexID, TurboID, and APEX2 targeted to the cytosol and ER membrane were induced with 300 ng/ml dox at 80% confluency for 20 hours. FlexID and TurboID expressing cells were then labeled for 10 minutes at 37 °C with 200 μM BCN and biotin respectively, then quenched with 3x washes with ice-cold DPBS. APEX2 expressing cells were pre-incubated for 30 minutes with 500 μM biotin-phenol and labeling was initiated by the addition of 1mM $H_2O_2$ for 1 minute. APEX2 labeling was quenched with a 3x wash with a solution of 10 mM ascorbate, 5mM Trolox, 10 mM sodium azide in DPBS, followed by 3x cold DPBS wash. All cells were then lysed in RIPA buffer with 1X PIC and 1mM PMSF for 15 minutes on ice, lysates were clarified by centrifugation and normalized to a protein concentration of 1mg/mL. Lysates from FlexID labeled cells were clicked with 4 μM mtz-biotin for 2 hours at 4 °C and cleaned up through methanol-chloroform precipitation. All samples were boiled for 10 minutes in 1x Protein Loading Buffer and then analyzed by SDS-PAGE gel followed by Western Blot as shown before.

### Yeast lysate labeling with BCN-AMP and purified FlexID

Concentrated yeast lysate was generated by mechanical grinding as previously described[22]. LplA variants were preincubated with BCN-AMP for 15 min at room temperature. In parallel, concentrated yeast lysate was preincubated with lipoyl-AMS and glycerol on ice. The *in vitro* labeling reaction was initiated by combining the LplA–BCN-AMP complex with the yeast lysate mixture and incubating at 37 °C. The final reaction mixture contained 25 μM LplA, 50 or 200 μM BCN-AMP, 1 mM lipoyl-AMS, and 5% glycerol in yeast lysate (>60 mg/mL total protein). A high concentration of lipoyl-AMS was included as a competitive inhibitor to ensure that labeling reflected the LplA-catalyzed BCN-AMP acylation (2nd step) rather than total reaction with free BCN generated by BCN-AMP hydrolysis. Effective competition was verified by incubating 25 μM LplA, 50 μM BCN, and 1 mM lipoyl-AMS in yeast lysate for 4 h. Minimal background labeling was observed under these conditions (**Extended Data Figure 4J**). Reactions were incubated for the indicated times, then quenched by rapid 20-fold dilution into ice-cold DPBS followed by methanol/chloroform precipitation (4:1). Protein pellets were resolubilized in 2% SDS, normalized to 1.5 mg/mL with RIPA buffer, and reacted with 4 μM mTz-BODIPY at 4 °C for 2 h. Samples were cleaned up using TCO-agarose. mTz-BODIPY–labeled proteins were analyzed by in-gel fluorescence and Coomassie staining. Band intensities were quantified in Fiji, and relative labeling was determined by dividing the fluorescence signal by the Coomassie signal after background subtraction. Fingerprint analysis of BCN chemical labeling and FlexID-catalyzed labeling was performed in Fiji after background subtraction (rolling ball radius: 15 pt, **Extended Data Figure 4I**).

**Labeling site mapping for FlexID**

ERM-FlexID transfected HEK293T cells were labeled with 200 μM BCN for 5min at 37 °C. Cells were lysed with RIPA buffer (50 mM Tris, pH 8.0, 150 mM NaCl, 0.1% SDS, 0.5% sodium deoxycholate, 1% Triton X-100) containing an EDTA-free protease inhibitor cocktail (Roche), and whole-cell lysates were normalized by using the BCA protein assay Kit (Pierce) on a microplate reader and adjusted to 2 mg/mL. The lysates were precipitated by methanol and chloroform (4:1). Then the protein pellets were resuspended in 0.4% SDS PBS. The proteins were reduced with 10 mM dithiothreitol (DTT) at 37 °C for 30 min and alkylated with 20 mM iodoacetamide (IAA) at 35 °C for 30 min in dark. After the alkylated proteins were precipitated with methanol–chloroform again, protein pallets were washed with ice cold methanol 3 times and then resuspended in 1 mL PBS buffer with ultrasonication. The protein was digested with 20 ng/μL of trypsin for 16 h at 37 °C. After the digested peptides were mixed with azide coated beads (CY75 beads)[5]. The click reaction was carried out at 37 °C for 2 h. The beads were washed with 1 mL of 8 M urea, 2 × 1 mL of PBS, 5 × 1 mL of $H_2O$ sequentially. Peptides were proceeded to on-beads cleavage by using 200 μL of 2% Formic acid for twice on thermomixer. The eluate containing the peptides was collected for LC-MS/MS analysis. Tryptic peptides were extended to the same length using peptide extender ver.0.2.2 alpha (https://schwartzlab.uconn.edu/pepextend)with the human proteome as a reference. Peptide sequences were acquired from three biological replicates. Peptide sequences were visualized using the pLogo web application[23].

**Comparison of PL enzyme labeling time courses**

HEK 293T cells stably expressing FlexID, TurboID, and APEX2 targeted to the ER membrane were induced with 300 ng/ml dox at 80% confluency for 20 hours. FlexID and TurboID expressing cells were then labeled for 10, 5, 1, and 0 minutes at 37 °C with 200 μM BCN and biotin respectively, then quenched with 3x washes with ice-cold DPBS. APEX2 expressing cells were pre-incubated for 30 minutes with 500 μM biotin-phenol and labeling was initiated by the addition of 1mM $H_2O_2$ for 5, 2, 1, and 0 minutes. APEX2 labeling was quenched with a 3x wash of a solution of 5 mM Trolox, 10 mM sodium azide, and 10 mM sodium ascorbate, followed by 3x washes with ice-cold DPBS. All cells were then lysed and analyzed as shown before.

**SDS-PAGE analysis of FlexID labeling with diverse substrates**

HEK 293T cells stably expressing FlexID targeted to the ER membrane were plated onto 12-well plates at 80% and induced with 300 ng/ml dox. After 20 hours of expression, cells were labeled with either 200 μM BCN for 10 minutes, 200 μM TCO for 10 minutes, 200 μM Azide for 10 minutes, 200 μM Alkyne for 2 hours, or 500 μM lipoic acid for 10 minutes at 37 °C. Labelling was quenched with 3x washes with ice-cold PBS. All cells were then lysed in EDTA-free RIPA buffer with 1x PIC and 1 mM PMSF for 15 minutes on ice, lysates were clarified by centrifugation and normalized to a protein concentration of 1 mg/mL. BCN and TCO labeled lysates were clicked with 4 μM mTz-biotin for 2 hours at 4 °C, while Azide and Alkyne labeled lysates were clicked with 100 μM alkyne-biotin or 100 μM azide-biotin respectively for 2 hours at 4 °C (1 mM $CuSO_4$, 500 μM BTTAA, and 2.5 mM freshly prepared sodium ascorbate). All lysates were cleaned up with methanol-chloroform precipitation. Cells were then analyzed on SDS-PAGE then transferred onto a PVDF for Western blot analysis. Blots were blocked in 5% (w/v) BSA in 1× tris-buffered saline with Tween (TBST) (Teknova) for 60 min at room temperature, incubated with mouse anti-V5-647 antibody (1:2,500) against FlexID's epitope tag and streptavidin-IRDye800 (1:10,000) in TBST for 1 h at room temperature, washed three times with TBST for 5 min each, and imaged on an Odyssey CLx imager (LICOR). For lipoic acid labeled lysates, the blot was incubated first with mouse anti-lipoic acid antibody (1:5,000), washed three times with TBST for 5 min each, stained with V5-AF647 (1:2,500)and streptavidin-IRDye800 (1:10, 000), then washed three times with TBST for 5 min each before imaging.

**SDS-PAGE analysis of FlexID labeling in different subcellular compartments**

HEK 293T cells were plated onto 12-well plates at 80% confluency and transfected with 500 ng of plasmids targeting FlexID to the outer mitochondrial membrane (OMM), mitochondrial matrix (mito), ER membrane (ERM), ER lumen (ERL), cytosol, and nucleus. After 20 hours of expression, all cells except ERL-FlexID cells were labeled for 10 minutes at 37 °C with 200 μM BCN. ERL-FlexID cells were labeled for 30 minutes at 37 °C with 200 μM BCN. All cells were then lysed in RIPA buffer with 1x PIC and 1mM PMSF for 15 minutes on ice, lysates were clarified by centrifugation and normalized to a protein concentration of 1 mg/mL. Normalized lysates were clicked with 4 μM mtz-BODIPY for 2 hours at 4 °C and cleaned up using methanol-chloroform precipitation (4:1). Cells were then analyzed on SDS-PAGE, imaged with a Typhoon Bio-Imager first. Then the SDS-PAGE gel was transferred onto a PVDF membrane for Western blot analysis. Blots were blocked in 5% (w/v) BSA in 1× tris-buffered saline with Tween (TBST) (Teknova) for 60 min at room temperature, incubated with mouse anti-V5-647 antibody (1:10,000) against FlexID's epitope tag in TBST for 1 h at room temperature, washed three times with TBST for 5 min each, and imaged on an Odyssey CLx imager (LICOR).

**SDS-PAGE analysis of C-terminal tagged FlexID labeling in different subcellular compartments**

HEK293T cells were plated onto 24-well glass-bottom plates at 60% confluency and transfected with 500 ng of DNA encoding each LplA–mCherry variant. After 20 hours of expression, cells expressing C-terminal LplA–mCherry fusions were labeled with 200 μM BCN for 20 minutes at 37 °C. Cells expressing N-terminal mCherry–LplA fusions were labeled with 200 μM BCN for 10 minutes at 37 °C. All cells were then lysed in RIPA buffer with 1x PIC and 1mM PMSF for 15 minutes on ice, lysates were clarified by centrifugation and normalized to a protein concentration of 1 mg/mL. Normalized lysates were clicked with 4 μM mTz-BODIPY for 2 hours at 4 °C and cleaned up with methanol-chloroform precipitation (4:1). Cells were then analyzed on SDS-PAGE and imaged with a Typhoon Bio-Imager.

**FlexID labeling with coumarin substrate**

HEK 293T cells stably expressing FlexID targeted to the ER membrane were plated onto glass bottom 24-well plates at 60% confluency and induced with 300ng/ml dox for 20 hours. Cells were then labeled for 10 minutes at 37 °C with 20 μM coumarin-AM2 supplemented with 0.1% Pluronic F-127. After labeling, cells were washed three times with warm DMEM + 10% FBS for 20min each. Cells were then fixed and permeabilized for 10 minutes at 4 °C with methanol pre-chilled at –20 °C. All cells were then blocked overnight at 4 °C with 1% BSA in DPBS. Primary staining was performed for 1 hour at RT with 1:1000 mouse-anti-V5 and 1:1000 rabbit anti-calnexin in 1% BSA in DPBS. Following 3x5 minute washes in DPBS, secondary staining was performed with 1:2000 mouse-568, and 1:2000 rabbit-647 for 1 hour at RT in 1% BSA in DPBS. Cells were then imaged at 100x magnification on a confocal microscope.

**Generation of FlexID AAV1/2**

To generate supernatant AAV, HEK293T cells were cultured in 6-well plate and transfected at approximately 80% confluency in opti-MEM reduced serum medium (Gibco). Per each well, the AAV vector containing the gene of interest (360 ng) and AAV packaging/helper plasmids AAV1 (180 ng), AAV2 (180 ng), and DF6 (720 ng) incubated with 10 μl PEI in 200 μl opti-MEM were used for transfection. After 20 h, the cell medium was replaced with complete DMEM. The cell medium containing the AAV was collected 48 h post transfection and filtered using a 0.45-μm filter.

For in vivo expression, AAV1/2 was produced in a large scale (3× 15-cm plates) accordingly. HEK293T cells were pelleted by centrifugation at 800 × g for 10 min and resuspended in 20 mL TBS (150 mM NaCl, 20 mM Tris, pH 8.0). Sodium deoxycholate (10% stock solution in water; Sigma) was added to a final concentration of 0.5%, along with benzonase nuclease (50 U/mL; Sigma). The suspension was incubated at 37 °C for 1 h and clarified by centrifugation at 3,000 × g for 15 min.

The resulting supernatant was loaded onto a HiTrap heparin column (GE Healthcare) pre-equilibrated with 10 mL TBS using a peristaltic pump (Gibson MP4). After virus binding, the column was washed sequentially with 20 mL 100 mM TBS (peristaltic pump), followed by 1 mL 200 mM TBS and 1 mL 300 mM TBS (syringe). Virus was eluted with 1.5 mL 400 mM TBS, 3.0 mL 450 mM TBS, and 1.5 mL 500 mM TBS. Eluted fractions were concentrated using a 15 mL centrifugal filter unit (100 kDa MWCO; Amicon) at 2,000 × g for ~1 min to a final volume of ~500 µL. The sample was further concentrated to ~100 µL using a 1.5 mL centrifugal filter unit (Amicon).

For titration, 2 µL of concentrated virus was incubated with 1 µL DNase I (NEB; 2 U/µL), 4 µL DNase I buffer, and 33 µL nuclease-free $H_2O$ at 37 °C for 30 min. DNase I was inactivated by heating at 75 °C for 15 min. Five microliters of the DNase-treated sample was mixed with 14 µL $H_2O$ and 1 µL Proteinase K (20 mg/mL; Thermo Fisher), followed by incubation at 50 °C for 30 min. Proteinase K was inactivated at 98 °C for 10 min. Two microliters of the resulting digest was used as template in qPCR reactions containing 5 µL SYBR Green master mix (2×; Thermo Fisher), 0.06 µL of each primer (0.3 µM final), and 2.88 µL nuclease-free $H_2O$.

Primers specific to the bovine growth hormone polyadenylation were used:

bGH poly(A) signal -forward: 5′- AAATGAGGAAATTGCATCGC

bGH poly(A) signal -reverse: 5′- TGCTATTGTCTTCCCAATCC

Standard curves were generated using purified, linearized AAV DNA plasmids containing bGH poly(A) signal sequences at input amounts of 0.05 ng, 0.1 ng, and 0.2 ng. Viral genome titers were calculated relative to the appropriate standard curve as described previously[24].

**FlexID and TurboID labeling in rat cortical neuron cultures**

All procedures were approved and carried out in compliance with the Stanford University Administrative Panel on Laboratory Animal Care, and all experiments were performed in accordance with relevant guidelines and regulations. Before dissection, plates were coated with 0.001% (w/v) poly-l-ornithine (Sigma-Aldrich) in DPBS (Gibco) at room temperature overnight, washed twice with DPBS, and subsequently coated with 5 μg/mL of mouse laminin (Gibco) in DPBS at 37 °C overnight. Cortical neurons were extracted from embryonic day 18 Sprague Dawley rat embryos (Charles River Laboratories, strain 400) by dissociation in Hank's balanced salt solution (Gibco) supplemented with 1 mM d-glucose and 10 mM HEPES pH 7.2 (Thermo Life Sciences). Cortical tissue was digested in papain according to the manufacturer's protocol (Worthington), then plated onto 12 well plates in neuronal plating medium at 37 °C under 5% $CO_2$. The neuronal plating medium is CNB media: neurobasal (Gibco), supplemented with 2% (v/v) B27 supplement (Life Technologies), 0.5% (v/v) fetal bovine serum, 1mM Sodium Pyruvate (Gibco), 1% GlutaMAX (Gibco), 1% penicillin-streptomycin (VWR, 5 units per ml penicillin, 5 μg/ml streptomycin). Every 3 days after plating, half of the media was removed from each well and replaced with neuronal growth medium. At 4 days in vitro (DIV), each well was infected with 200 μL of supernatant AAV1/2 along with a media change. At DIV11, cells were treated with a final concentration of 200 µM biotin or 200 µM BCN for 5 min in the incubator. Cells were lysed or fixed for western blot analysis or immunofluorescence detection as described above.

**FlexID labeling in the mouse brain**

All experimental and surgical protocols were approved by the University of California, Davis, Institutional Animal Care and Use Committee. Here, 5-7 week-old C57BL/6J mice (Jackson Laboratory Strain 000664) were used for all experiments. Mice were maintained on a 12-h reverse light–dark cycle (lights on at 21:00) at 22 °C and 40–60% humidity, group-housed with same-sex cage mates and given ad libitum access to food and water. For mouse stereotaxic surgeries, mice were maintained under anesthesia with 1.5–2% isoflurane and placed in a stereotaxic apparatus (RWD) on a heating pad. The fur on the top of the skull was removed and antiseptic iodine and 70% alcohol were used in alternation to clean the scalp. Sterile ocular lubricant (Dechra) was administered to the eyes of the mice to protect them from drying out. A midline scalp incision was made, and 0.1% hydrogen peroxide was applied to the skull. A craniotomy was made above the injection site. Virus was then injected into the targeted region using a 33-gauge beveled needle (WPI) and a 10 µl Hamilton syringe controlled by an injection pump (WPI). For all surgeries, 500 nl of virus ($5.9 \times 10^{11}$ viral genomes per ml titer) was bilaterally injected into the targeted mPFC brain region (coordinates: ML, ±0.5; AP, +1.98; DV, −2.25) at a rate of 150 nl/min.

Mice were allowed to recover for one week. Mice were then placed under anesthesia, injected with 500 nl of 10 mM BCN or vehicle in PBS solution (pH = 7) at the same stereotaxic coordinates. After 30 minutes of labeling, mice were perfused with ice-cold PBS and their brains were immediately dissected and cut in a roughly 1 mm coronal section using a brain block and razor blades on ice. Both hemispheres were pooled for each sample and dissected brain tissues were flash frozen in 1.5 mL tubes.

Frozen brain tissues were resuspended with 150 µl of ice-cold buffer, 50 mM Tris/HCl, pH 7.5, 150 mM NaCl, 1 mM EDTA supplemented with protease inhibitor cocktail and PMSF. A tissue homogenizer was then used to process the tissue. The samples were lysed by adding 200 µl of ice-cold buffer containing 0.4% SDS, 2% TrixonX-100, and 2% sodium deoxycholate. Samples were incubated with benzonase (50 U/ml) at 4 °C for 30 min and briefly sonicated for 25 times (20% Amp, 1 sec pulses on/off). Lysates were cleared via centrifugation at 20,000g at 4 °C for 30 min, and then processed for in gel fluorescence and western blot analysis as described above.

**Generation of proteomic samples**

Stable HEK293T cell lines were generated with doxycycline (dox)-inducible expression of ERM-APEX, NES-APEX, ERM-FlexID, NES-FlexID, ERM-TurboID, and NES-TurboID. To normalize expression and labeling efficiency, dox titrations were performed at 33 ng/mL, 100 ng/mL, and 333 ng/mL. The optimized concentrations were: FlexID-NES, 333 ng/mL; FlexID-ERM, 100 ng/mL; APEX-NES, 333 ng/mL; APEX-ERM, 33 ng/mL; TurboID-NES, 333 ng/mL; and TurboID-ERM, 100 ng/mL. After overnight induction (19–20 hours), cells reached ~80% confluency. Labeling conditions were as follows: FlexID samples were treated with 200 µM BCN for 5 min at 37 °C; TurboID samples were treated with 200 µM biotin for 5 min at 37 °C; and APEX2 samples were pretreated with 500 µM biotin-phenol for 30 min at 37 °C, followed by 1 min of $H_2O_2$ labeling. FlexID and TurboID samples were quenched with three washes of ice-cold DPBS. APEX2 samples were quenched by adding an equal volume of 2× quenching buffer (20 mM ascorbate, 10 mM Trolox, 20 mM sodium azide in DPBS), followed by two additional washes with 1× quenching buffer, followed by three additional washes of ice-cold DPBS. All samples were harvested by scraping, pelleted at 300 × g for 3 min, flash frozen in liquid nitrogen, and stored at −80 °C.

Frozen pellets were lysed in 50 mM Tris-HCl (pH 7.4), 150 mM NaCl, 0.25% sodium deoxycholate, 1% NP-40, 1 mM EDTA, 1× protease inhibitor cocktail (Sigma-Aldrich), and 1 mM PMSF by gentle pipetting, followed by a 10-min incubation at 4 °C. Lysates were clarified by centrifugation at 20,000 × g for 10 min at 4 °C. Protein concentrations were measured with the Pierce BCA Protein Assay Kit (Thermo Fisher) and normalized to 1.6 mg/mL. FlexID lysates were reacted with 4 μM mTz-biotin for 2 h at 4 °C with shaking (850 rpm), followed by incubation with 40 μL TCO-agarose for 2 h at 4 °C with rotation. TCO-agarose was removed by centrifugation at 20,000 × g for 10 min at 4 °C. For enrichment of biotinylated proteins, 60 μL streptavidin-coated magnetic beads (Pierce) were prewashed twice with RIPA buffer, then incubated with 1.5 mg total eluates for 24 h at 4 °C with rotation. Beads were washed sequentially with: RIPA buffer (2×), 1 M KCl (1×), 0.1 M $Na_2CO_3$ (1×), 2 M urea in 10 mM Tris-HCl (pH 8.0) (1×), and RIPA buffer (2×). For Western blotting and silver staining, 10% of enriched proteins were eluted by boiling in 3× protein loading buffer supplemented with 20 mM DTT and 2 mM biotin. For proteomic analysis, the beads were then resuspended in 1 mL of fresh RIPA lysis buffer and transferred to a new tube. The beads were resuspended in 60 μL of digestion buffer and flash frozen in liquid nitrogen and stored at −80 °C. All the procedures were carried out in lo-bind tubes.

The enriched proteins with magnetic beads were washed twice with 1 mL of 25 mM ammonium bicarbonate (ABC), and the resulting beads were resuspended in 500 μL of 25 mM ABC containing 6 M urea. The proteins were reduced with 10 mM dithiothreitol (DTT) at 35 °C for 30 minutes and then alkylated with 20 mM iodoacetamide (IAA) at 37 °C for 30 minutes in the dark. Following this, the beads were washed eight times with 1 mL of 25 mM ABC to remove urea, resuspended in 200 µL of 25 mM ABC buffer containing 10 ng/µL trypsin, and digested at 37 °C overnight on a thermomixer. The resulting peptides were collected in fresh microcentrifuge tubes, and the beads were washed twice with 100 µL of 25 mM ABC; the supernatants were combined with the peptide solution, and 0.1% formic acid was added before drying with speedvac. Dried peptides were shipped to Wei Qin's laboratory (Tsinghua University) with cold packs for further processing and preparation for LC-MS/ MS analysis.

**Liquid chromatography and mass spectrometry (for 5 minute FlexID experiment)**

Peptides were first loaded onto a trap column (PepMap Neo C18, 5 µm, 300 µm × 5 mm; Thermo Fisher Scientific, 174500) and subsequently separated on an in-house packed analytical column (100 µm × 20 cm, ReproSil-Pur 120 C18-AQ, 1.9 µm; Dr. Maisch GmbH, r119.aq). Chromatographic separation was performed on a Vanquish™ Neo UHPLC system (Thermo Fisher Scientific) at a flow rate of 0.5 µL/min. The mobile phases consisted of solvent A (water with 0.1% formic acid) and solvent B (80% acetonitrile with 0.1% formic acid). The gradient was programmed as follows: 0–4 min, 4–5% B; 4–109 min, 5–20% B; 109–150 min, 20–35% B; 150–159 min, 35–99% B; held at 99% B for 7 min, followed by column re-equilibration. Eluted peptides were analyzed on Q Exactive-Plus Orbitrap mass spectrometers (Thermo Fisher Scientific) equipped with an EASY-Spray ion source operated at a spray voltage of +2.2 kV and a capillary temperature of 320 °C. Full MS survey scans were acquired over an m/z range of 350–1800 at a resolution of 70, 000 with an AGC target of $3 \times 10^6$. For FlexID site-mapping samples, data-dependent acquisition (DDA) was employed with Top20 MS/MS scans per cycle. MS/MS spectra were acquired at a resolution of 17,500 with an AGC target of $1 \times 10^5$. Monoisotopic precursor selection was enabled, using a default precursor charge state of 2. Fragmentation was performed by higher-energy collisional dissociation (HCD) with a normalized collision energy of 28% and an isolation window of 1.6 m/z. For PL ERM/NES comparision samples, data-independent acquisition (DIA) was performed. MS/MS spectra were acquired at a resolution of 17, 500 with an AGC target of $1 \times 10^6$. Fragmentation was performed by HCD with a normalized collision energy of 28%, using a fixed first mass of 200 m/z, a default precursor charge state of 3.

**Liquid chromatography and mass spectrometry**

Desalted peptides were resuspended in 9 μL of 3% MeCN/0.1% FA and analyzed by online nanoflow liquid chromatography tandem mass spectrometry (LC-MS/MS) using a Thermo Orbitrap Exploris 480 MS (ThermoFisher Scientific) coupled on-line to a Proxeon Easy-nLC 1200 (ThermoFisher Scientific). Four microliters of each sample were loaded onto a microcapillary column (360 μm outer diameter × 75 μm inner diameter) containing an integrated electrospray emitter tip (10 μm), packed to approximately 30 cm with ReproSil-Pur C18-AQ 1.5 μm beads (Dr. Maisch GmbH) and heated to 50 °C. Each fraction was run on a 110min-method, including a linear 84 min gradient from 94.6% solvent A (0.1% formic acid) to 27% solvent B (99.9% acetonitrile, 0.1% formic acid), followed by a linear 9 min gradient from 27% solvent B to 54% solvent B.
The Thermo Orbitrap Exploris 480 operated in a data-dependent acquisition mode, measuring MS1 with 60,000 resolution, 300% normalized AGC target, and m/z range from 350 to 1800. MS2 spectra of the top 20 most abundant ions per cycle were acquired at 45,000 resolution, 30% AGC target, 0.7 m/z isolation window, and 34 normalized collision energy. The dynamic exclusion time was set to 20 s, and the peptide match and isotope exclusion functions were enabled.

**Mass spectrometry data processing**

MS/MS data were processed using DIA-NN (v2.2.0). A reference spectral library was generated from the UniProt human proteome (UniProt ID: UP000005640, 20,663 gene entries with a release date of June 18, 2025), which was subjected to in silico digestion with deep learning–based prediction. The library generation was performed with the following parameters: 1% FDR (0.01), precursor m/z range of 300–1800, fragment ion m/z range of 200–1800, precursor charge range of 1–4, and peptide lengths of 7–30 amino acids. Trypsin/P was specified as the protease, allowing up to one missed cleavage. Fixed and variable modifications included N-terminal methionine excision, carbamidomethylation of cysteine, N-terminal acetylation, and methionine oxidation, with a maximum of one variable modification per peptide.

Experimental .raw files were processed in DIA-NN using the predicted reference library. For the ERM vs. NES analysis, negative control samples were excluded due to their substantially lower peptide content compared with PL-enzyme–labeled samples. Mass accuracy was optimized automatically. Protein quantification was performed with match-between-runs (MBR) enabled, and protein inference was applied. Proteotypic peptides were annotated using the "Reannotate" option, and common contaminants from the Cambridge Centre for Proteomics (CCP) database were automatically excluded from quantification. The scoring mode was set to peptidoforms, with all other parameters left at default values. DIA-NN outputs were filtered at a precursor q-value < 1% to ensure accuracy. For the ERM vs. Negative control groups analysis, all samples were included and processed as described above, with the exception that cross-run normalization was disabled. Normalized QuantUMS quantities from the protein groups ('pg_matrix') were used for further analysis.

We used the Proteomics Toolset for Integrative Data Analysis (Protigy, v1.1.8, Broad Institute, https://github.com/broadinstitute/protigy) to calculate moderated t-test P-values for regulated proteins. For the ERM vs. NES comparison, no imputation was applied. For the ERM vs. Negative control comparison, missing values in the negative control groups were addressed using MinDet imputation to account for missing-not-at-random (MNAR) patterns[25]. When a protein was absent in all control runs, the imputed value was set to 90% of the lowest protein abundance observed across the run. When a protein was absent in only a subset of runs, the imputed value was set to 90% of the lowest abundance of that protein observed across the remaining runs[26].

For the ERM versus NES comparison, normalized QuantUMS intensity values for each protein ID were $\log_2$-transformed and subjected to median normalization. For the ERM versus negative control comparison, normalized QuantUMS intensity values for each protein ID were also $\log_2$-transformed; however, median normalization was performed across predefined groups to account for systematic differences: Group 1 included all PL-labeled samples, Group 2 included all untransduced samples, and Group 3 included TurboID-positive samples lacking biotin supplementation. A two-sample moderated t test was performed on the data to compare experimental groups using an internal R-Shiny package based in the limma library. p-values associated with every protein were adjusted using the Benjamini–Hochberg FDR approach (https://doi.org/10.1111/j.2517-6161.1995.tb02031.x).

**Generation of proteomic lists for the ER membrane**

Complete mass spectrometry data for the ER membrane (ERM) proteomic experiments are shown in **Extended Data Table 4.** To select cutoffs for proteins labeled by the indicated ERM-ligase over proteins labeled by the corresponding cytosol-targeted ligase, we classified the detected proteins into three groups:

1. ERM proteins (**Extended Data Table 3, Tab 1**; true positive list of 90 well-established ERM proteins[6]).
2. Non-secretory proteins (**Extended Data Table 3, Tab 3**; false positive list of 6,465 human proteins that are not predicted to be secretory by Phobius[27], the Human Protein Atlas[28] (protein localized to endoplasmic reticulum, Golgi apparatus, plasma membrane, vesicles, nuclear membrane, cell junctions; or predicted membrane proteins and predicted secreted proteins), the Plasma Proteome Database (2021 build)[29], literature (reference cited in table), or are not annotated with the following Gene Ontology terms[30,31]: GO:0005783, GO:0005789, GO:0007029, GO:0030867, GO:0048237, GO:0061163, GO:0016320, GO:0030868, GO:0006983, GO:0000139, GO:0051645, GO:0031985, GO:0005796, GO:0005795, GO:0005794, GO:0007030, GO:0090168, GO:0005886, GO:0007009, GO:1903561, GO:0070062, GO:0005576, GO:0031012, GO:0005615, GO:0005769, GO:0035646, GO:0005765, GO:0090341, GO:0090340, GO:0005635, GO:0007084, GO:0007077, GO:0006998, GO:0051081, GO:0005641, GO:0031965, GO:0005637, GO:0071765, GO:0048471, GO:1905719, GO:0031982, GO:0006906, GO:0048278, GO:0032587, GO:0016021, GO:0005887, GO:0005768, GO:0071816, GO:0031526, GO:0005913, GO:0072546, GO:1990440, GO:0030968, GO:1902236, GO:1990441, GO:0034976, GO:0005788, GO:0005790, GO:1902237, GO:0070059, GO:0005786, GO:0005793, GO:0044322, GO:0098554, GO:0005791, GO:1902010, GO:0043001, GO:0005802, GO:0006888, GO:0006890, GO:0005801, GO:0012510, GO:0006892, GO:0042147, GO:0034499, GO:0032588, GO:0006895, GO:0030140, GO:0051684,

GO:0000042, GO:0032580, GO:0030173, GO:0006891, GO:0030198, GO:0031668, GO:0010715, GO:0035426, GO:1903053, GO:1903551, GO:0005578, GO:1903055, GO:0001560, GO:0022617, GO:0006887, GO:0012505)

3. All other proteins.

To calculate optimal cut-offs, the proteins were first ranked in a descending order according to ERM/NES enrichment ratio. Common contaminants were removed. We then calculated the true positive rate (TPR) and false positive rate (FPR) we would obtain if we retained only proteins above that enrichment ratio. We defined TPR as the fraction of class (1) proteins above the enrichment ratio in question, and FPR as the fraction of class (2) above the enrichment ratio in question. A receiver operating characteristic (ROC) curve was plotted accordingly for each comparison. For the 5min data, partial area under the curve of each ROC curve at different FPR cutoff was computed using the trapezoid rule in Prism (GraphPad). After filtering with adjusted p-values ($q < 0.05$), we selected enrichment ratios that maximize the difference between TPR and FPR as our cutoffs (**Extended Data Figures 7C, 7E, 7F**) to produce the final proteomes (**Extended Data Table 4**). Alternatively, we selected enrichment ratios that resulted in a fixed FPR = 0.10 as our cutoffs to produce the final proteomes for the DIA dataset (**Extended Data Table 4**).

To assess the specificity of our proteomes (**Figure 3K**, **Extended Data Figure 7H**), we report the percentage of proteins present in **Extended Data Table 3**, a list of 14,198 human proteins with secretory annotation according to Phobius, the Human Protein Atlas (protein localized to endoplasmic reticulum, Golgi apparatus, plasma membrane, vesicles, nuclear membrane, cell junctions; or predicted membrane proteins and predicted secreted proteins), the Plasma Proteome Database(2021 build), literature (reference cited in table), or are not annotated with the following Gene Ontology terms: GO:0005783, GO:0005789, GO:0007029, GO:0030867, GO:0048237, GO:0061163, GO:0016320, GO:0030868, GO:0006983, GO:0000139, GO:0051645, GO:0031985, GO:0005796, GO:0005795, GO:0005794, GO:0007030, GO:0090168, GO:0005886, GO:0007009, GO:1903561, GO:0070062, GO:0005576, GO:0031012, GO:0005615, GO:0005769, GO:0035646, GO:0005765, GO:0090341, GO:0090340, GO:0005635, GO:0007084, GO:0007077, GO:0006998, GO:0051081, GO:0005641, GO:0031965, GO:0005637, GO:0071765, GO:0048471, GO:1905719, GO:0031982, GO:0006906, GO:0048278, GO:0032587, GO:0016021, GO:0005887, GO:0005768, GO:0071816, GO:0031526, GO:0005913, GO:0072546, GO:1990440, GO:0030968, GO:1902236, GO:1990441, GO:0034976, GO:0005788, GO:0005790, GO:1902237, GO:0070059, GO:0005786, GO:0005793, GO:0044322, GO:0098554, GO:0005791, GO:1902010, GO:0043001, GO:0005802, GO:0006888, GO:0006890, GO:0005801, GO:0012510, GO:0006892, GO:0042147, GO:0034499, GO:0032588, GO:0006895, GO:0030140, GO:0051684, GO:0000042, GO:0032580, GO:0030173, GO:0006891, GO:0030198, GO:0031668, GO:0010715, GO:0035426, GO:1903053, GO:1903551, GO:0005578, GO:1903055, GO:0001560, GO:0022617, GO:0006887, GO:0012505).

The specificity of the 'entire human proteome' reported in **Figure 3K, and Extended Data Figure 7H** was calculated as the percentage of human proteins that are not present in category (2) non-secretory proteins, as defined above; i.e., the proteins that are present in the list of 14,198 human proteins with secretory annotation in **Extended Data Table 3**. To assess the recall/sensitivity of our ERM proteomes, we utilized a list of 90 true positive ERM proteins (**Figure 3L**, **Extended Data Figures 7I, Extended Data Table 3**).

**Evaluating PL enzyme specificity without using cytosolic spatial reference**

To assess the spatial specificity of PL enzyme labeling, we performed receiver operating characteristic (ROC) analyses. Proteins labeled by the indicated ERM-ligase were compared against non-specific bead binders detected in untransduced negative controls. Proteins were ranked in descending order based on the adjusted p values and then ERM-to-untransduced enrichment ratio. True positive rates (TPR) and false positive rates (FPR) were calculated as in the ERM vs. NES comparison, using a reference set of 90 well-established ERM proteins and an additional 25 ERM proteins that are identified by all three proximity labeling enzymes (true positives, **Extended Data Table 3, Tab 8**) and 6,465 non-secretory proteins (false positives). ROC curves were then generated for each comparison.

ERM-targeted TurboID labels ER membrane proteins in HEK293T cells even without exogenous biotin supplementation, due to the low basal levels of biotin present in the culture medium. Consequently, the apparent spatial specificity of TurboID labeling is higher than that of FlexID. To account for potential differences in ER membrane protein labeling over extended labeling periods, we performed receiver operating characteristic (ROC) analyses. Proteins labeled by ERM-TurboID were compared against NES-TurboID in the absence or presence of 5 min of 200 μM Biotin treatment. Proteins were ranked in descending order based on the ERM-to-NES enrichment ratio. True positive rates (TPR) and false positive rates (FPR) were calculated as described for the ERM vs. NES comparison, using a reference set of 90 well-established ERM proteins (true positives) and 6,465 non-secretory proteins (false positives). ROC curves were then generated for each comparison.

For the ROC analysis of ERM spatial specificity by normalization to HEK bulk lysate protein abundance, ERM raw intensity scores were ratioed against raw intensity scores from (Guzman, Martinez-Val, and Ye et al.) measurements of HEK lysate by DIA proteomics[4]. Proteins were ranked in descending order based on the ERM-to-total lysate enrichment ratio. True positive rates (TPR) and false positive rates (FPR) were calculated as above, and ROC analysis enabled comparison between datasets.

**Generation and characterization of CRBN-FlexID Jurkat cells**

CRBN-FlexID Jurkat cells were generated by introducing doxycycline (dox)-inducible expression of FlexID-CRBN into $CRBN^{-/-}$ Jurkat cells, using the lentiviral transduction protocol described above. To validate CRBN-FlexID expression, CRBN-FlexID Jurkat cells ($0.5 \times 10^6$ cells/mL) were induced with 1 µg/mL doxycycline overnight. Cells were lysed in RIPA buffer supplemented with PIC and PMSF on ice for 10 minutes at 4 °C. Lysates were cleared by centrifugation at 20,000 × g for 10 minutes at 4 °C, and protein concentrations were normalized using the BCA Protein Assay Kit (Pierce), adjusting all samples to 1.5 mg/mL. Equal amounts of protein were resolved on 4–12% SDS–PAGE gels and transferred to PVDF membranes. Membranes were stained with Ponceau S (Sigma) to verify equal loading, then blocked in 5% BSA in TBST for 1 hour at room temperature. Blots were incubated overnight at 4 °C with Rabbit anti-CRBN antibody (1:1000) and mouse anti–β-actin antibody (1:5000) diluted in blocking buffer. After three 5-minute washes with TBST, membranes were incubated for 1 hour at room temperature with anti-mouse-IRDye680-conjugated secondary antibodies and anti-rabbit-IRDye800-conjugated secondary antibodies (1:10,000) diluted in blocking buffer. Following three additional 5-minute washes, membranes were imaged using an Odyssey CLx imager (LI-COR). Antibody sources and dilutions are provided in the Supplementary Information.

**PLF assay**

*Cell treatment:* FlexID-expressing cells were treated with 200 μM BCN in complete medium. Labeling was quenched by washing cells three times with cold PBS. Cells were lysed in IP lysis buffer (25 mM Tris pH 7.4, 150 mM NaCl, 1% NP-40, 1 mM EDTA, 5% glycerol) supplemented with 1× protease inhibitor cocktail (PIC) and 1 mM PMSF, followed by high-speed centrifugation to remove debris. Protein concentration of the cleared lysates was determined using the BCA Protein Assay Kit (Pierce) and normalized to 1 mg/mL.

*Antibody and bead preparation:* IP-grade antibodies (1 μg) were incubated with 1 μL of Pierce Protein A or G beads in 10 μL of IP lysis buffer for 1 h at room temperature with rotation. Antibody-conjugated beads were washed twice with IP lysis buffer and diluted into buffer supplemented with either mTz-biotin or mTz-BODIPY.

*Immunoprecipitation and flow analysis:* 0.05-0.1 μL of antibody-conjugated Protein A or G beads were incubated overnight at 4 °C with 10–25 μL of cleared lysate under constant mixing (1300 rpm or end-over-end rotation). Phycoerythrin-based detection: Beads were washed three times with PBST (0.05% Tween-20) and incubated with streptavidin–phycoerythrin (1:500, Jackson ImmunoResearch, 016-110-084) for 1 h at room temperature with mixing. After three additional PBST washes, beads were resuspended in PBST and analyzed by flow cytometry at 4 °C. Samples were agitated for 5 s prior to acquisition and run at a flow rate of 1 μL/s on a ZE5 Cell Analyzer (Bio-Rad) until 50, 000 beads were collected. Single beads were gated on forward scatter area (FSC-A) versus side scatter area (SSC-A) to identify the main bead population (P1).

P1 was further gated on FSC-A versus FSC-H to exclude aggregates (P2). Phycoerythrin signal intensity (median fluorescence intensity) was then quantified from P2. Data were analyzed using FlowJo v10 (BD Biosciences). BODIPY-based detection: Beads were washed twice with IP lysis buffer, twice with high-salt wash buffer (25 mM Tris-HCl pH 7.6, 1 M NaCl, 1% NP-40, 0.1% SDS, 0.5% sodium deoxycholate), and twice more with IP lysis buffer. Beads were resuspended in IP lysis buffer and analyzed by flow cytometry at 4 °C using the same gating strategy as above. BODIPY signals were quantified from the P2 population.

The following gating strategy was used for the PLF assay:

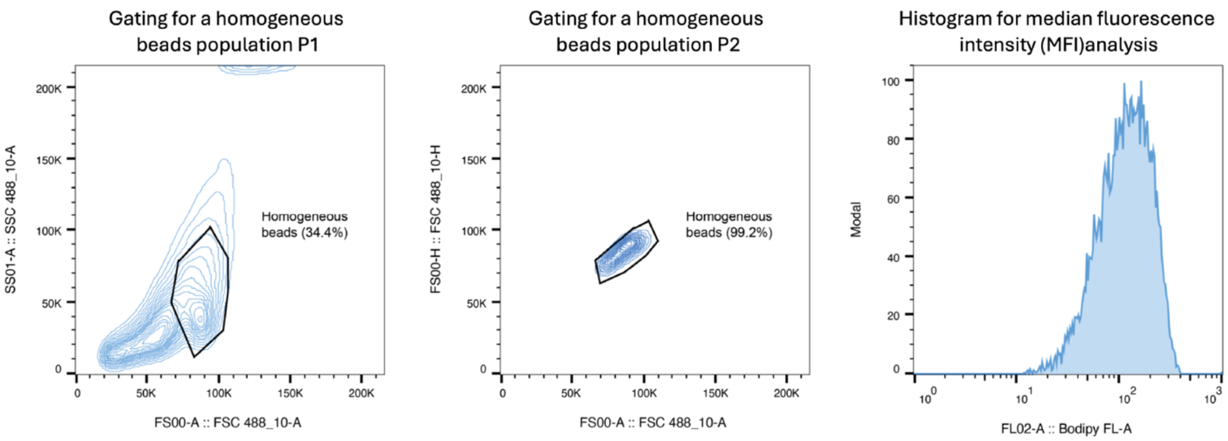


**Rapamycin dose-response**

HEK293T cells were transiently transfected with FRB-Flag-FlexID and APEX-V5-FKBP12 plasmids (1:1 ratio) and incubated overnight. The following day, cells were treated with a 3-fold serial dilution of Rapamycin (final concentrations: 300 nM to 0) for 2 h at 37 °C. Cells were then labeled with 200 µM BCN for 5 min at 37 °C, washed three times with cold PBS, and lysed in IP lysis buffer with 1× PIC and 1 mM PMSF. Lysates were clarified by centrifugation at 20,000 × g for 10 min. For bead preparation, 5 µg of V5 antibody (Invitrogen, R960-25) was incubated with 5 µL of Pierce Protein G beads in 50 µL IP lysis buffer for 1 h at room temperature with rotation.

Antibody-conjugated beads were washed twice with IP lysis buffer and resuspended in 50 μL of the same buffer.

For immunoprecipitation, 50 μL of cleared lysate was incubated overnight at 4 °C with 0.5 μL of diluted antibody–bead conjugates and 4 μM mTz-BODIPY under end-over-end rotation. Beads were washed twice with IP lysis buffer, twice with high-salt wash buffer, and twice again with IP lysis buffer. Beads were resuspended in IP lysis buffer and analyzed by flow cytometry at 4 °C following the gating strategy described in the PLF assay section.

**Pomalidomide and HRZ-98-1 dose-response**

CRBN-FlexID–expressing Jurkat cells (0.5 × $10^6$ cells/mL) were induced with 1 μg/mL doxycycline overnight, then pretreated with 5 μM MLN4924 for 2 h at 37 °C. After MLN4924 treatment, 200 μL of the cell suspension (2 × $10^6$ cells/mL) was added to 10 μL of the 21× compound solution, mixed by pipetting at least five times, and incubated for 1 h at 37 °C. This yielded final compound concentrations of 0, 1, 5, 25, 125, 625, and 3125 nM. Subsequently, 10 μL of 4.4 mM BCN was added, mixed thoroughly, and incubated for 30 min at 37 °C. Cells were pelleted at 500 × g for 3 min, the supernatant was removed, and pellets were washed three times with cold PBS. Cells were lysed with 40 μL of total IP lysis buffer with 1x PIC and 1 mM PMSF, followed by centrifugation at 3300 × g for 30 min to remove debris.1.5 μg of Wee1 antibody (Cell Signaling Technology, D10D2), IKAROS antibody (Invitrogen, #PA5-85570), and CSNK1A1 antibody (Proteintech, 55192-1-AP) were incubated with 1.5 μL Pierce Protein A beads in a 30 μL IP lysis buffer for 1h at room temperature, respectively. Antibody-conjugated beads were washed twice with IP lysis buffer and diluted into 300 μL lysis buffer supplemented with 8 μM mTz-biotin.

For immunoprecipitation, protein A beads were incubated overnight at 4 °C with 10 μL of cleared lysate and 10 μL of diluted beads at 1300 rpm. Beads were then washed three times with PBST and incubated with streptavidin–phycoerythrin (1:500 dilution, Jackson ImmunoResearch, 016-110-084) for 1 h at room temperature with shaking at 1300 rpm. After three additional PBST washes, beads were resuspended in 90 μL PBST and analyzed by flow cytometry at 4 °C.

**PLF screen of molecular glue library**

The screen was performed following the procedure described in the Molecular Glue Dose-Response (pomalidomide and HRZ-98-1) section, with minor modifications to the antibody and bead preparation step. Specifically, 5.5 μg of Wee1 antibody (Cell Signaling Technology, D10D2), IKAROS antibody (Invitrogen, #PA5-85570), or CSNK1A1 antibody (Proteintech, 55192-1-AP) was incubated separately with 5.5 μL of Pierce Protein G beads in 30 μL of IP lysis buffer for 1 h at room temperature. The antibody-conjugated beads were then washed twice with IP lysis buffer and diluted into 300 μL of lysis buffer supplemented with 8 μM mTz-biotin.

**PLF analysis of protein subcellular localization**

HEK293T cells were transfected with FlexID constructs targeted to the ER membrane (458 ng DNA/well) or mitochondrial matrix (1042 ng DNA/well) in 12-well plates at ~80% confluency. After 20 h of expression, cells were labeled with 200 μM BCN for 5 min at 37 °C, washed three times with cold DPBS, pelleted at 500 × g for 5 min, and flash frozen. Cell pellets were lysed in IP buffer with protease inhibitors and PMSF for 25 min on ice. Lysates were clarified by centrifugation (20,000 × g, 10 min, 4 °C) and normalized to 2 mg/mL using the BCA assay. Antibodies against calnexin (ER marker), MTCO1 (mitochondrial marker), GRSF1 (mitochondrial marker), Rabbit IgG (control IgG) were conjugated to Pierce Protein A magnetic beads (0.7 μg antibody per 0.7 μL beads, 1 h, room temperature, rotation), washed twice in IP buffer, and resuspended in 14 μL IP buffer. Lysates at a final concentration of 1 mg/mL were incubated with 1 μL diluted antibody–bead conjugates, 5% control rabbit IgG (Fortis Life Sciences, P120-101), and 4 μM mTz-BODIPY overnight at 4 °C with end-over-end mixing. Beads were washed five times with PBST (0.05% Tween-20), resuspended in 100 μL PBST, and analyzed by flow cytometry at 4 °C following the gating strategy described above. Background from IgG control beads is subtracted for each sample. The flowthroughs from the overnight incubation were collected and analyzed by in-gel fluorescence to assess FlexID labeling efficiency and signal quality.

**BCN dose-response**

HEK293T cells were transiently transfected with ERM-FlexID plasmid and incubated overnight. The following day, cells were treated with a 3-fold serial dilution of BCN (final concentrations: 200 μM to 0) for 5 min at 37 °C. Cell were washed three times with cold PBS and lysed in IP lysis buffer with 1× PIC and 1 mM PMSF. Lysates were clarified by centrifugation at 20,000 × g for 10 min. For bead preparation, 15 μg of Calnexin antibody was incubated with 15 μL of Pierce Protein G beads in 300 μL IP lysis buffer for 1 h at room temperature with rotation. Antibody-conjugated beads were washed twice with IP lysis buffer and resuspended in 150 μL of the same buffer.

For immunoprecipitation, 50 μL of cleared lysate was incubated overnight at 4 °C with 0.5 μL of diluted antibody–bead conjugates and 4 μM mTz-BODIPY under end-over-end rotation. Beads were washed twice with IP lysis buffer, twice with high-salt wash buffer, and twice again with IP lysis buffer. Beads were resuspended in IP lysis buffer and analyzed by flow cytometry at 4 °C following the gating strategy described in the PLF assay section.

**Quantification and data analysis**

All graphs were created using GraphPad Prism or matplotlib (Python). Error bars represent standard deviation unless otherwise noted. For comparison between two groups, p-values were determined using two-tailed Student's t tests. For multiple comparisons, p-values were calculated using the statistical methods specified in the corresponding figure legends or Methods section.
